# Integrative protein modeling in RosettaNMR from sparse paramagnetic restraints

**DOI:** 10.1101/597872

**Authors:** Georg Kuenze, Richard Bonneau, Julia Koehler Leman, Jens Meiler

**Author notes:** To whom correspondence should be addressed., Georg Kuenze, Ph.D., Vanderbilt University, Center for Structural Biology, 465 21st Ave S, 5140 MRB3, Room 5154E, Nashville, TN 37235.

## Abstract

Computational methods to predict protein structure from nuclear magnetic resonance (NMR) restraints that only require assignment of backbone signals hold great potential to study larger proteins and complexes. Additionally, computational methods designed to work with sparse data add atomic detail that is missing in the experimental restraints, allowing application to systems that are difficult to investigate. While specific frameworks in the Rosetta macromolecular modeling suite support the use of certain NMR restraint types, use of all commonly measured restraint types together is precluded. Here, we introduce a comprehensive framework into Rosetta that reconciles CS-Rosetta, PCS-Rosetta and RosettaNMR into a single framework, that, in addition to backbone chemical shifts and nuclear Overhauser effect distance restraints, leverages NMR restraints derived from paramagnetic labeling. Specifically, RosettaNMR incorporates pseudocontact shifts, residual dipolar couplings, and paramagnetic relaxation enhancements, measured at multiple tagging sites. We further showcase the generality of RosettaNMR for various modeling challenges and benchmark it on 28 structure prediction cases, eight symmetric assemblies, two protein-protein and three protein-ligand docking examples. Paramagnetic restraints generated more accurate models for 85% of the benchmark proteins and, when combined with chemical shifts, sampled high-accuracy models (≤ 2Å) in 50% of the cases.

**Significance Statement:** Computational methods such as Rosetta can assist NMR structure determination by employing efficient conformational search algorithms alongside physically realistic energy functions to model protein structure from sparse experimental data. We have developed a framework in Rosetta that leverages paramagnetic NMR data in addition to chemical shift and nuclear Overhauser effect restraints and extends RosettaNMR calculations to the prediction of symmetric assemblies, protein-protein and protein-ligand complexes. RosettaNMR generated high-accuracy models (≤ 2Å) in 50% of cases for a benchmark set of 28 monomeric and eight symmetric proteins and predicted protein-protein and protein-ligand interfaces with up to 1Å accuracy. The method expands Rosetta’s rich toolbox for integrative data-driven modeling and promises to be broadly useful in structural biology.

## Introduction

Structural biology increasingly focuses on large and complex protein systems, such as membrane proteins, multi-subunit assemblies or cellular machineries, seeking insights into their role in health and disease. For such challenging targets for protein structure determination, the major technologies X-ray crystallography, nuclear magnetic resonance (NMR) spectroscopy, and cryo-electron microscopy (cryo-EM) struggle to obtain experimental data that unambiguously defines atomic detail. Limitations in resolution, accuracy and coverage of individual experimental methods can be bridged by data-driven, integrative modeling that considers multiple types of information simultaneously. Programs like Rosetta (1), HADDOCK (2), the Integrative Modeling Platform (IMP) (3), or BCL::Fold (4) are pioneering applications that integrate information from multiple biophysical and biochemical sources and have led to structures of complex protein systems (5, 6).

NMR data are one of the most useful sources of structural information for integrative modeling. Advances in solution-state NMR such as protein deuteration (7), relaxation-optimized spectroscopy (TROSY (8)), methyl labeling (9), ^13^C direct detection (10), and non-uniform sampling (11) have improved sensitivity and spectral resolution and pushed the limit of protein NMR to molecular weights of 50-100 kDa. Nonetheless, NMR datasets from large proteins remain sparse and of low resolution, making 3D structure determination challenging. To facilitate structure determination, computational modeling tools are required that can leverage limited NMR data and translate them into accurate structural models.

A powerful strategy to supplement chemical shifts (CSs) and sparse nuclear Overhauser effect (NOE) data is the use of long-range (up to 40 Å) paramagnetic NMR restraints, arising from interactions of protein nuclear spins with paramagnetic metal ions or nitroxide spin-labels. Metal ions can be introduced via metal binding sites or attached to the protein using coordinating tags. Advances in chemical synthesis of metal ion-binding tags (12) and strategies for their covalent attachment (13) have broadened the applicability of paramagnetic NMR to many biomolecular systems (14, 15).

Three types of paramagnetic NMR data are commonly used for protein structure determination: paramagnetic relaxation enhancements (PRE), residual dipolar couplings (RDC), and pseudocontact shifts (PCS). PREs can be detected in any paramagnetic system and provide long-range distance restraints. PCSs and RDCs require anisotropic magnetic susceptibility of the paramagnetic ion (e.g. lanthanides), giving rise to a non-vanishing Δχ-tensor and leading to partial alignment of the protein in the external magnetic field. The Δχ-tensor defines a coordinate system in the protein that is centered on the metal ion and relates the observed PCS and RDC to the polar coordinates of a nuclear spin or spin pair. The PCS is a particularly rich structural restraint because it combines both distance and angular information (16).

The Rosetta suite can use different types of NMR data, mainly for protein structure prediction. The original RosettaNMR method used backbone CSs to find structurally similar peptide fragments in the Protein Databank (PDB), which are assembled *de novo* guided by limited NOE restraints (17). This was later expanded to RDCs from conventional (non-paramagnetic) alignment media (18). In 2003, Meiler et al. showed that RosettaNMR can be used for structure determination of small proteins to atomic detail from unassigned NMR spectra predicting CSs, NOEs, and RDCs on the fly (19). Critical for this approach was the iterative filtering of models through comparison of predicted with experimental CSs (20). This concept has been extended with CS-Rosetta for structure determination of larger proteins from backbone-only CS and RDC data (21) up to 25 kDa molecular weight (22). Incorporating sparse side-chain NOEs from deuterated protein samples and improving conformational sampling algorithms increased the application limit of CS-Rosetta to proteins up to 40 kDa (23) and enabled prediction of helical membrane protein structures (24). Further advances to CS-Rosetta came from the inclusion of structural templates from homologous proteins (25). Rosetta was also combined with PCS data (PCS-Rosetta), achieving model qualities comparable to CS-Rosetta (26). This approach was extended to PCS data from multiple tagging sites and applied to the solution structure of ERp29, which uncovered flaws in a previous NOE-derived NMR structure (27).

These prior results were obtained with customized frameworks in Rosetta that do not interface with each other and are difficult to integrate into various protocols. For example, previous implementations of PCSs in Rosetta were tailored to protein structure prediction but cannot be used for protein-protein or protein-ligand docking. Moreover, while specific combinations of restraints can be used in Rosetta, it is currently impossible to generally exploit the complementary nature of long-range and local restraints by combining PCSs, RDCs and PREs with CSs and NOEs. However, combining these restraints will presumably improve the accuracy, precision and completeness of a model, especially where individual restraint types are too sparse or erroneous. Further, other integrative methods are either tailored to a specific application (e.g. HADDOCK for protein-protein docking) or cannot handle the wide array of NMR data (e.g. the IMP does not support paramagnetic NMR data). Therefore, generalizing the frameworks in Rosetta for integrated use of various types of experimental data is highly beneficial for modeling larger and more sophisticated biomolecular systems.

Here we reconcile RosettaNMR with CS-Rosetta and PCS-Rosetta to create a unified framework that combines long-range paramagnetic data with CSs and NOEs and integrates them into a variety of Rosetta applications. This required substantial refactoring and generalization of pre-existing NMR scoring methods. A new PRE energy method was added to account for conformational flexibility of the spin-label via an ensemble rotamer approach. We show that this framework can be easily combined with popular modeling protocols such as RosettaAbinitio (28), RosettaDock (29), RosettaLigand (30) and RosettaSymmetry (31) for a variety of structure determination tasks. Extensive documentation (https://www.rosettacommons.org/docs/latest/application_documentation/RosettaNMR-with-Paramagnetic-Restraints) and protocol captures in the supplementary material allow ease-of-use and facilitate reproducibility for non-expert Rosetta modelers. Our framework vastly improves the extensibility and accuracy when modeling from sparse NMR restraints, therefore expanding Rosetta’s rich toolbox for incorporating experimental data into an integrative modeling pipeline. RosettaNMR is available as part of Rosetta which is free of charge for academic, government, and non-profit users (https://www.rosettacommons.org/software).

## Results

### RosettaNMR offers a versatile platform for integrative modeling with paramagnetic NMR restraints

The success of protein structure modeling from limited NMR data can be markedly improved if different types of NMR data are combined in one calculation. To realize such an integrative modeling approach, we extend the suite of NMR tools in Rosetta, which we collectively term RosettaNMR, by paramagnetic NMR data in combination with CS and NOE data in one single framework (Fig 1).

**Figure 1:**
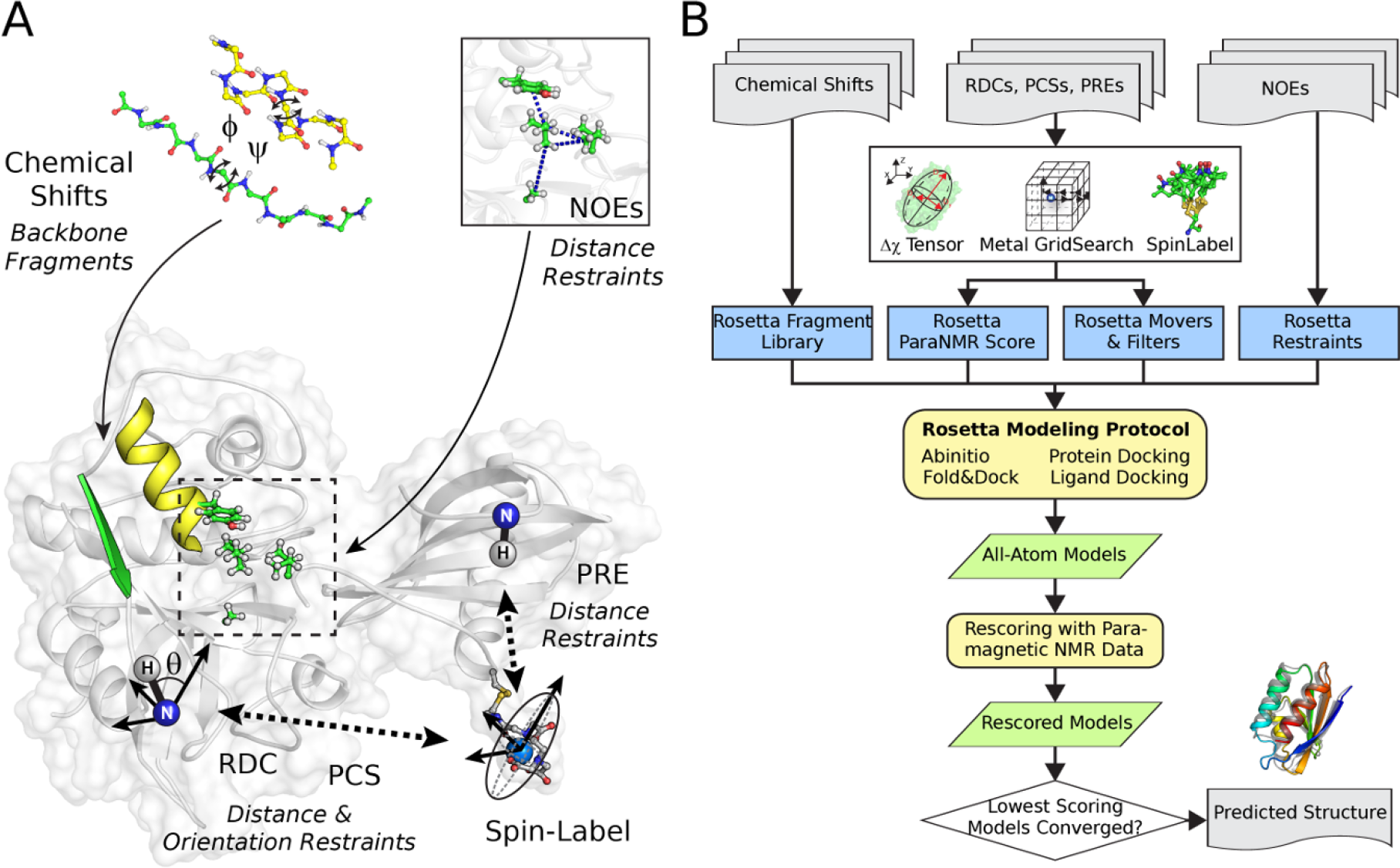
Macromolecular modeling with RosettaNMR using different types of restraints. **(A)** NMR data types accessible through RosettaNMR: CSs aid in the selection of short peptide fragments with known backbone conformations to assemble protein structures *de novo*. NOE-derived distances restrain the protein’s tertiary structure and interface with binding partners. Long-range paramagnetic NMR effects provide an orthogonal set of restraints on the position and orientation of internuclear vectors within the macromolecular system. **(B)** Overview of the experimental workflow of integrative NMR-guided protein structure modeling with Rosetta. NMR data can be incorporated at different stages in the Rosetta protocol to guide conformational sampling or modify the energy landscape.

We focused our implementation on PCS, RDC and PRE data which are most informative in terms of the protein structure. The PCS and RDC are manifested as change in the chemical shift or spin coupling constant originating from molecular alignment induced by the metal’s Δχ-tensor. Whereas the PCS depends both on the distance and orientation to the metal ion, the RDC provides an angular restraint for inter-atomic vectors (Fig 1A). The PRE is the contribution to the spin relaxation rate and can be converted into a long-range distance restraint (Fig 1A).

Each data type was implemented as a restraint energy penalty. During model scoring, the PCS score is calculated by computing the Δχ-tensor by singular value decomposition coupled with a grid search over the metal ion coordinates to find the best possible fit. The RDC score is computed analogously by determining the molecular alignment tensor, which is related to the Δχ-tensor by a constant pre-factor if alignment originates solely from the paramagnetic metal ion but is different for an external anisotropic medium (e.g. bicelles). To generalize score calculation for any para- or diamagnetic data, we decided to fit PCSs and RDCs separately. In addition, Δχ-tensor values obtained by fitting RDCs tend to be smaller than those calculated from PCSs, because RDCs are more sensitive to local protein motions. Considering structural flexibility becomes even more important when interpreting PREs due to the many degrees of freedom (DOFs) of the spin-label; we have adapted the ensemble averaging approach from Iwahara and Clore for computation of the PRE score (32). The spin-label is modeled as an ensemble of side-chain conformations which are approximated from a rotamer library stored in the Rosetta database. An effective distance <*r*^−6^> is calculated between the protein nuclear spin and all spin-label conformations, whose structural variability is further represented by an order parameter *S*^2^.

Rosetta allows manipulating the protein through modeling objects accessible from high-level scripting languages such as PyRosetta (33) and Rosetta Scripts (34). This allows the user to develop new customized protocols by mixing and matching different strategies (Fig 1B). The main modeling tools fall into two categories: Movers and Filters. Movers change the conformation of a biomolecule and Filters decide whether the given conformation should go into the next stage. One can easily devise new strategies of how paramagnetic restraints can be used to guide conformational sampling or model filtering. We exemplify this by applying PCSs to ligand docking for which we have designed a new PCS ligand scoring grid method and a rigid-body sampling Mover. Furthermore, RosettaNMR allows new spin-labels to be added to the chemical database in a straightforward manner to accommodate new spin-label designs.

### Paramagnetic NMR data enhance sampling of native-like models in de novo structure prediction

We tested RosettaAbinitio (28) on 28 monomeric proteins in our benchmark set (**Tab S1**) using different types and combinations of NMR data: CSs, RDCs, PCSs and PREs. For three proteins, both PCSs and RDCs were available and could be tested. Table 1 summarizes the RMSD_100_ of the best 1% of models ranked by either RMSD or score. Score-vs-RMSD plots of selected targets are displayed in Fig 2, and for all targets in **Fig S1** and **S2**. Noticeably, paramagnetic NMR data greatly improve sampling of near-native models, i.e. they bias the fragment assembly toward the native structure. In 29 out of 34 test cases, the best 1% of models ranked by RMSD had a lower RMSD_100_ to the native structure when paramagnetic NMR data were included (Fig 3A+F), and the average RMSD_100_ improved from 4.5 Å to 3.5 Å (Tab 1; see **Fig S3A** for statistical analysis of RMSD improvement).

**Table 1:**
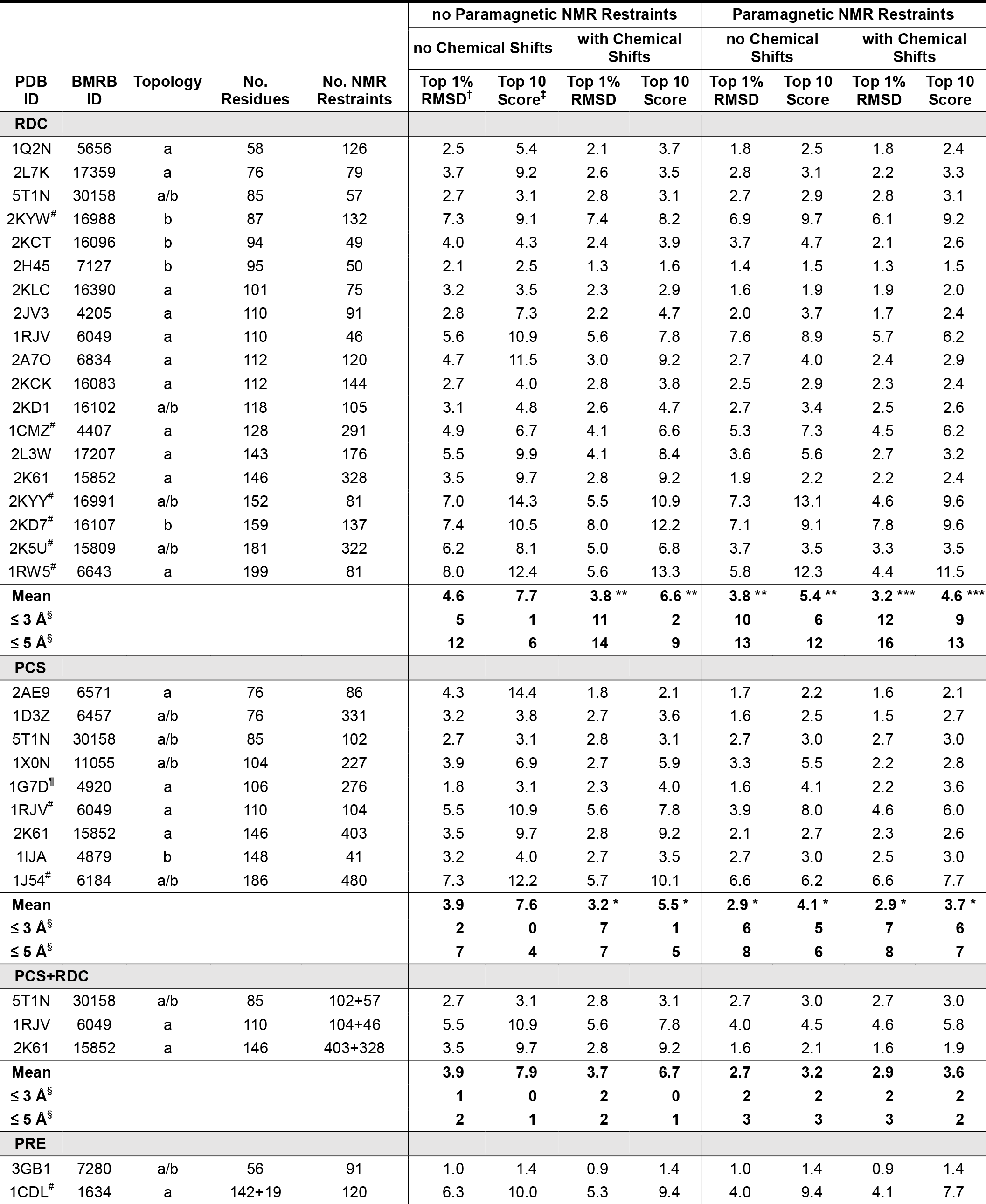
Structure prediction benchmark of monomeric proteins with paramagnetic NMR restraints and chemical shifts.

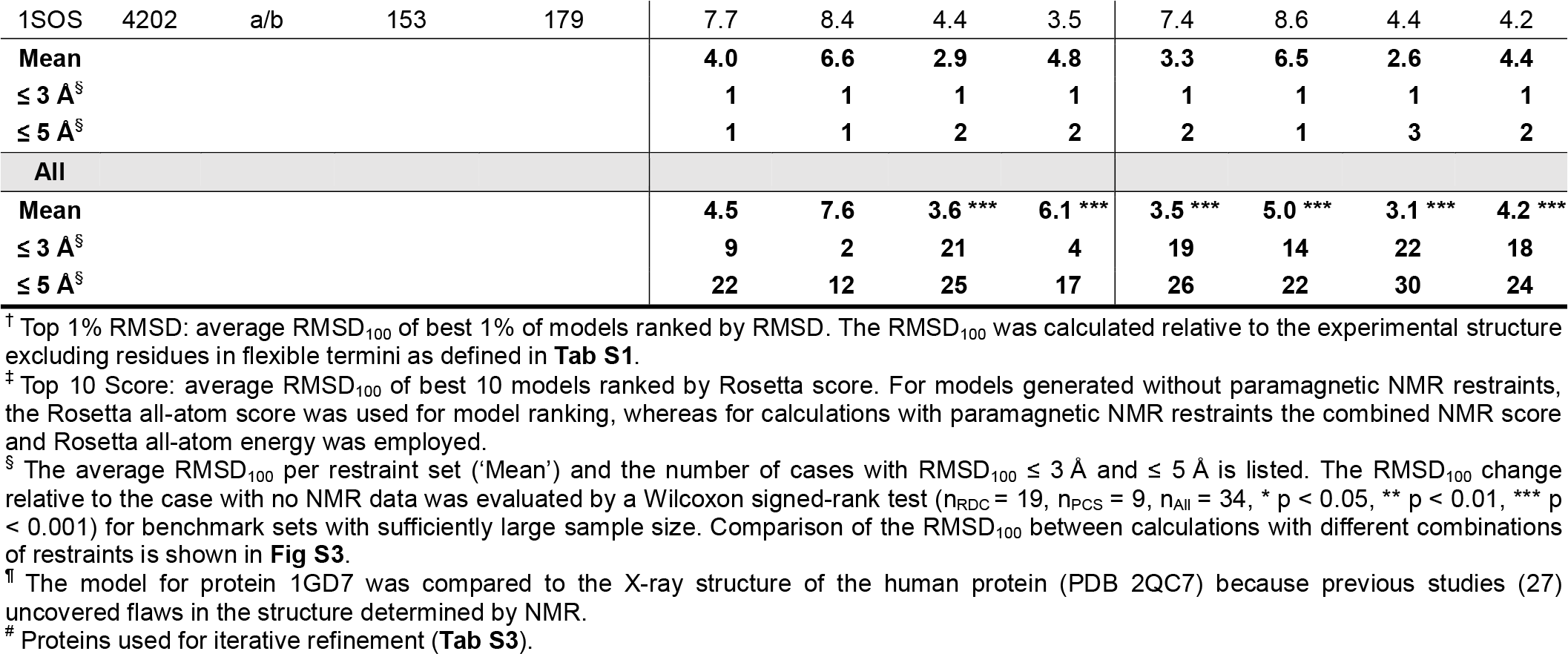

**Figure 2:**
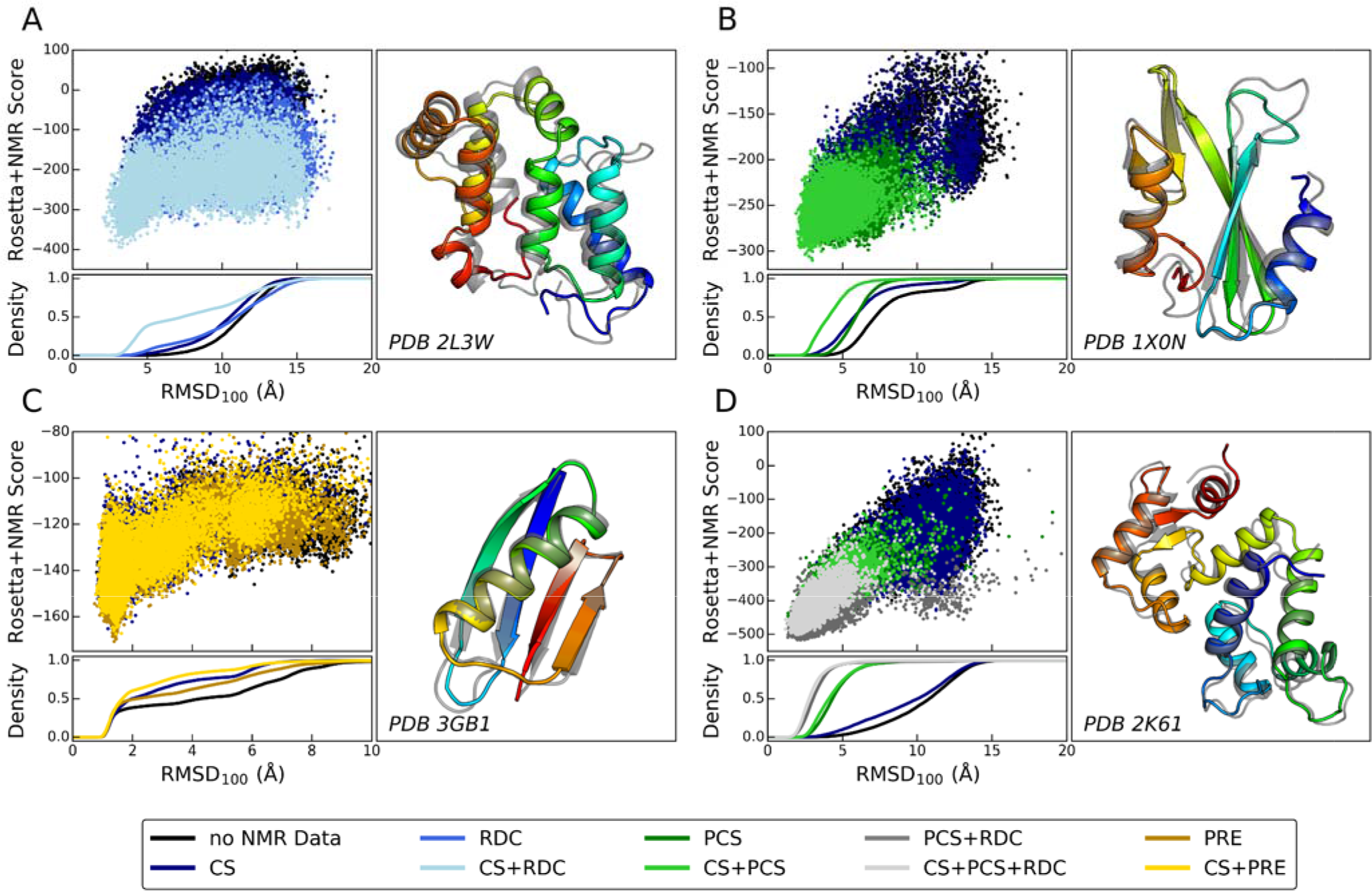
*De novo* structure prediction of monomeric proteins with paramagnetic and CS NMR data. Four selected benchmark proteins for which prediction utilized either **(A)** RDCs (phycobilisome rod linker protein from S. elongatus, PDB 2L3W), **(B)** PCSs (Grb2 SH2 domain, PDB 1X0N), **(C)** PREs (B1 domain of streptococcal protein G, PDB 3GB1) or **(D)** RDCs and PCSs (calmodulin, PDB 2K61), respectively. A complete list of the results of all prediction targets is given in **Fig S1** and **S2**. For each protein, the combined Rosetta and paramagnetic NMR score and the cumulative fraction of models versus the models’ RMSD_100_ relative to the experimental structure is shown. The paramagnetic NMR score was the **(A)** RDC, **(B)** PCS, **(C)** PRE or **(D)** combined PCS+RDC restraint score. The largest increase in the density of models with low RMSD_100_ is observed when CSs and paramagnetic NMR restraints are applied together. The lowest-scoring model from this experiment is depicted as ribbon diagram (colored in rainbow) and compared to the experimentally determined structure shown in gray. The superposition was optimized for residues in ordered regions as defined in the footnote to **Tab S1** and flexible termini are not shown for clarity.

**Figure 3:**
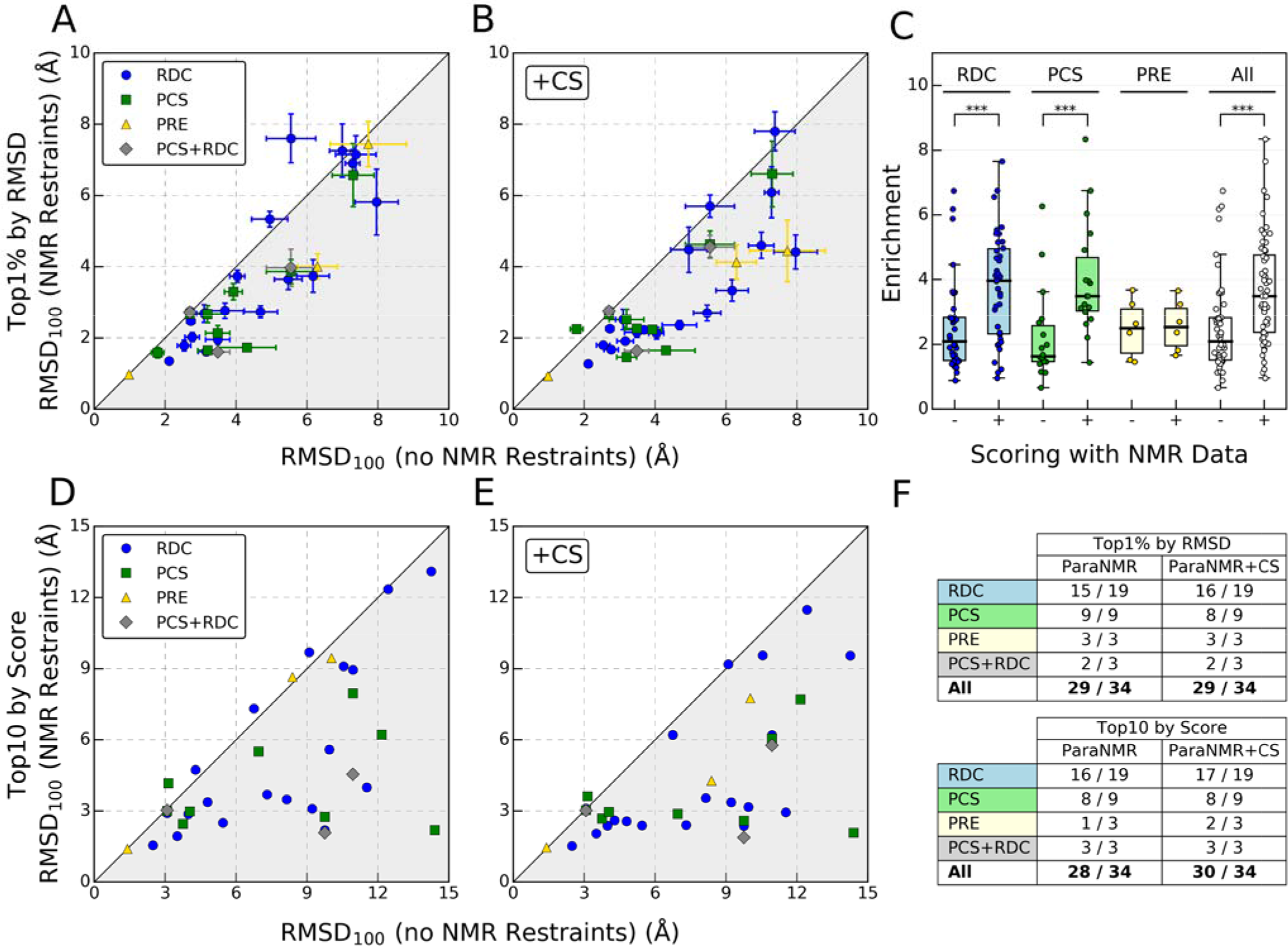
Effect of paramagnetic and CS data on sampling and scoring of *de novo* predicted models. Comparison of the average RMSD_100_ (± S.D.) of the top 1% of models ranked by RMSD between Rosetta models predicted without NMR data (x-axis) and **(A)** with only paramagnetic NMR restraints (y-axis) or **(B)** when all CS and paramagnetic NMR data were used. Gray areas indicate an RMSD_100_ improvement. **(C)** Model enrichment calculated when models were scored with the standard Rosetta all-atom energy function (−) or an energy function which included an additional RDC, PCS or PRE score term (+), respectively. Box-and-Whisker plot with whiskers drawn from the lowest and highest enrichment value still within 1.5 IQR of the lower and upper quartile. Comparison was made by two-tailed Wilcoxon signed-rank test (n_RDC_ = 38, n_PCS_ = 18, n_All_ = 56, * p < 0.05, ** p < 0.01, *** p < 0.001). **(D)** and **(E)** Comparison of average the RMSD_100_ among the top ten scoring models rom structure predictions with paramagnetic NMR data only or in combination with CSs. Gray areas indicate improved model accuracy. **(F)** Summary of the fraction of cases in which NMR-restrained structure prediction yielded more accurate models, which were identified either after ranking by RMSD or score.

Comparing models generated with either paramagnetic NMR restraints or CSs (**Fig S4**), we find that the former had a slightly better performance on model sampling: for 72% of the proteins paramagnetic NMR data led to a lower RMSD_100_. However, both restraint types improved model quality compared to unrestrained *de novo* prediction: With CSs the average RMSD_100_ for the best 1% models over all proteins dropped from 4.5 Å to 3.6 Å, improving the RMSD_100_ for 24 out of 34 cases (Tab 1, **Fig S3A** and **S4**). With paramagnetic NMR restraints the average RMSD_100_ dropped to 3.5 Å, improving for 29 of 34 cases.

Model sampling was even more enhanced when CSs were combined with RDCs, PCSs and PREs (Fig 3B), showing that these data have an orthogonal beneficial effect on the conformational search. In 29 of 34 cases, the best 1% of models ranked by RMSD had a lower RMSD_100_ to the native structure when paramagnetic NMR and CS data were used (Fig 3F), and the average RMSD_100_ decreased from 4.5 Å to 3.1 Å (Tab 1; see also **Fig S3A** for statistical analysis of RMSD change).

The gradual improvement in model sampling by including CSs, paramagnetic restraints or a combination of both can be seen in the score-vs-RMSD and density plots in Fig 2, **S1** and **S2**. The fraction of proteins for which a high-accuracy model (RMSD_100_ ≤ 2 Å) could be generated, increased from 9% (no NMR data) to 35% (CSs), 35% (paramagnetic restraints) and 50% (CSs and paramagnetic restraints). Medium-accuracy models (RMSD_100_ ≤ 3 Å) could be sampled in 62% (no NMR data), 68% (CSs), 76% (paramagnetic restraints) and 76% (CSs and paramagnetic restraints) of the cases.

### Paramagnetic NMR data enrich for native-like models in scoring

A similar improvement in model quality was observed when they were ranked by score (Tab 1, Fig 3D+E). This is more realistic since in a blind structure prediction models must be identified predominantly by score. Here, we calculated the score as combination of the Rosetta all-atom energy and the RDC, PCS or PRE score with a weighting factor that was optimized for each NMR dataset and protein as described under Methods. For those cases where PCSs and RDCs were used, both the PCS and RDC score was added to the Rosetta energy with weights proportional to the logarithm of the number of PCSs and RDCs.

For 28 of 34 and 30 of 34 targets the ten best-scoring models had a lower average RMSD_100_ when paramagnetic restraints were used individually or combined with CSs (Fig 3D+E). The average RMSD_100_ of the ten lowest-scoring models over all benchmark proteins improved from 7.6 Å (no NMR data) to 5.0 Å (paramagnetic restraints) and 4.2 Å (CSs and paramagnetic restraints) (Tab 1, **Fig S3B**). The number of proteins for which the RMSD_100_ of the top ten models was ≤ 3 Å increased from two (no NMR data) to 14 (paramagnetic restraints) and 18 (CSs and paramagnetic restraints). Moreover, 22 (paramagnetic restraints) and 24 (CSs and paramagnetic restraints) proteins fell within a 5 Å cutoff.

To investigate the ability of the RDC, PCS and PRE score to recognize near-native models we calculated model enrichment (maximum value is 10, see **Method S4**) displayed in Fig 3C. To this end, models built without paramagnetic NMR data were rescored with RDCs, PCSs or PREs and the all-atom energy function. The average enrichment over the benchmark set increased from 2.5 to 3.7 (RDCs: 2.5 → 3.8; PCSs: 2.2 → 4.1; RDCs + PCSs: 1.9 → 4.2). No significant increase in enrichment was seen for the PRE score (2.5 → 2.6), showing that it failed to further improve model selection with Rosetta’s all-atom energy function. Although we only have experimental PRE datasets from three proteins, the observations match those made with simulated PRE data for symmetric proteins (see below). We speculate that the lower discriminative power of the PRE score is due to the fact that PREs are purely distance-dependent and have a narrow dynamic range as they decrease with r^−6^ to the spin-label. This is in contrast to PCSs which are sensitive to orientation and distances and cover a larger range due to their r^−3^ dependence. Consequently, the PCS score drastically improves model discrimination when included in Rosetta’s all-atom energy function.

### Paramagnetic NMR data facilitate protein model selection

Experimental data can be compared to back-calculated data from predicted structural models for model selection and validation. The top RosettaNMR models satisfy their experimental data remarkably well (Fig 4A-C for proteins 2K61, 5T1N and 1CDL, Fig 4D and **Tab S2**). A direct measure of the agreement provides the NMR Q factor (see **Method S4**), which we calculated for the ten best-scoring models of each target in the benchmark set. We saw a steady improvement of the Q-factor with the inclusion of more NMR data, and combining CSs and paramagnetic data yielded Q-factors comparable to native structures (Fig 4D).

**Figure 4:**
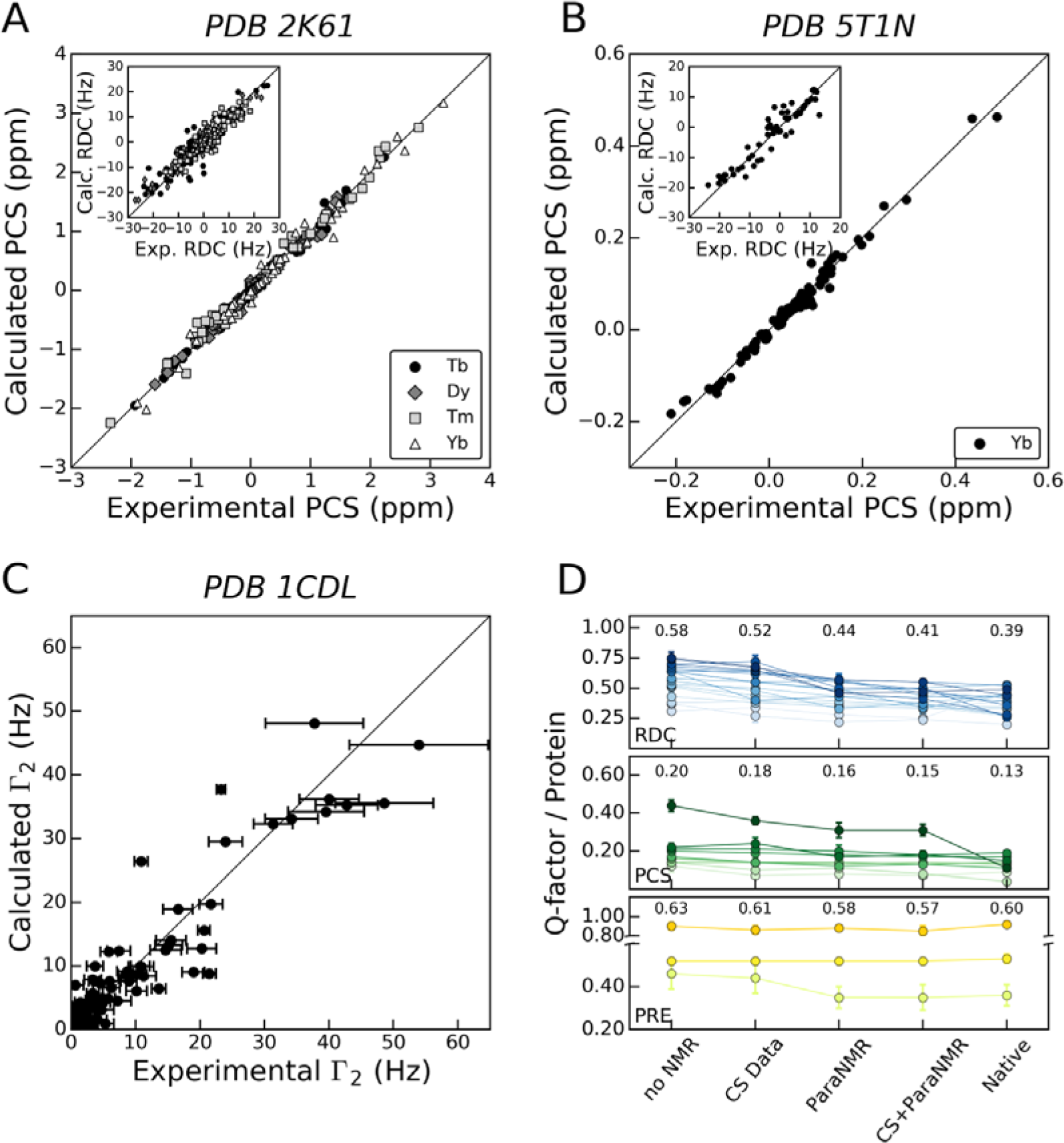
Agreement of experimental with model-predicted paramagnetic restraints. Correlation between experimental PCSs and RDCs for **(A)** calmodulin (PDB 2K61) and **(B)** NPr (PDB 5T1N) with PCS and RDC values back-calculated from their respective lowest-scoring model. **(C)** Correlation of experimental and calculated PREs for the lowest-scoring model of MLCK-peptide-bound calmodulin (PDB 1CDL). In each case, the lowest scoring model obtained with CSs and paramagnetic NMR data was used for the analysis. **(D)** Average NMR Q-factor (± S.D.) of the ten lowest-scoring models for each *de novo* structure prediction target versus the amount and type of NMR data: no NMR – without NMR data, CS – with CSs, ParaNMR – with RDC, PCS or PRE data, CS+ParaNMR – with CSs and RDC, PCS or PRE data, native – experimental structure after constrained minimization in Rosetta. The individual proteins modeled with either RDC, PCS or PRE restraints are represented by using different shades of blue, green or yellow, respectively, and lines between data points are used for easier visualization. The average Q-factor across all proteins is given on the top of each column. A complete list of Q-factor values is given in **Tab S2** and **S4**.

We used two measures to assess the quality and estimate success of structure prediction: a) low Q-factors confirm good agreement with the experimental data and b) model convergence indicates that a topology is repeatedly sampled and satisfies the combined Rosetta and NMR score. We defined convergence as the average RMSD between the best-scoring model and the next nine low-scoring models. We found that paramagnetic restraints and CSs were highly beneficial for model convergence and a steady improvement in the RMSD was observed when including more NMR data (**Fig S5A**). In 28 of 34 test cases (82%), the model created with CSs and paramagnetic restraints had a convergence ≤ 4 Å (**Tab S2** and **Fig S5A**). This RMSD cutoff is employed in the calculation of other protein structure similarity metrics (e.g. GDT-TS and MaxSub) and models below this cutoff usually share the same global topology. The other six proteins (2KYW, 2KYY, 2KD7, 1RW5, 1CDL, 1J54) had limited convergence, identifying their model as unreliable and allowing for their rejection (see **Fig S6** for more discussion).

### Iterative resampling with paramagnetic NMR data further improves model quality

Inspection of the six prediction failures according to our convergence criterion (2KYW, 2KYY, 2KD7, 1RW5, 1CDL, 1J54) revealed no high-accuracy model (≤ 2 Å) and only a small number of intermediate-accuracy models (≤ 3 Å) after *de novo* prediction, even when combining CSs with paramagnetic restraints. Insufficient sampling was also noticed for another three proteins in the benchmark set (1CMZ, 1RJV, 2K5U) with consequently high RMSD_100_ for their top-scoring models. To test if enhanced sampling could further improve model accuracy, we applied the Rosetta Iterative Hybridize refinement protocol, incorporating paramagnetic restraints. As previously described (35), Iterative Hybridize generates new hybrid conformations by recombining templates obtained from low-energy structures together with *de novo* fragment insertion. A pool of low-energy conformations is maintained, and the worst scoring models are periodically replaced with new models.

We ran 30 rounds of Iterative Hybridize and selected the offspring generation of models at each step by their combined Rosetta and NMR score. The RMSD_100_ steadily improved for eight of nine proteins; the RMSD_100_ of the ten lowest-scoring models decreased by 3.2 Å on average, improving the number of proteins with RMSD_100_ ≤ 5 Å for the ten lowest-scoring models from one of nine to six of nine (**Tab S3** and **Fig S7**). For two of nine proteins at least one high-accuracy model was present in the pool after the final refinement step and for four of nine proteins more than 1% of models had an RMSD_100_ better 3 Å.

### RosettaNMR with paramagnetic NMR data improves structure prediction of symmetric proteins

We further applied RosettaNMR to symmetric proteins, focusing on systems with cyclical (Cn) and dihedral (Dn) symmetry as they are most abundant in nature. We hypothesized that RDCs, PCSs and PREs provide sufficient information to fold the asymmetric monomer and sample symmetric rigid body DOFs toward low-energy and experimentally relevant conformations. We incorporated point symmetry into the RDC, PCS and PRE scoring methods, allowing the use of experimental data collected on the whole protein assembly through symmetric tagging. Hence, time-consuming asymmetric tagging of single subunits to break the symmetry for the ease of assignment is not required.

We combined RosettaNMR and the Fold-and-Dock protocol (36), which simultaneously models the asymmetric monomer and symmetric complex, applicable to interleaved and non-interleaved topologies. The protocol was tested with published RDCs and simulated PCSs and PREs (Tab 2 and **Tab S1**), and the type of symmetry was assumed to be known. We created 5,000 models for each protein and restraint set. In addition, we employed a two-step protocol for proteins 1RJJ and 2M89 for which Fold-and-Dock failed to build models < 10 Å to the native structure (see Methods).

**Table 2:** Structure prediction benchmark of symmetric proteins with paramagnetic restraints.

| PDB ID | BMRB ID | Symmetry | No. Residues | No. NMR Restraints* | no NMR Restraints |  | RDC |  | PCS |  | PRE |  |
| --- | --- | --- | --- | --- | --- | --- | --- | --- | --- | --- | --- | --- |
|  |  |  |  |  | Top 1% RMSD<sup>†</sup> | Top 10 Score<sup>‡</sup> | Top 1% RMSD | Top 10 Score | Top 1% RMSD | Top 10 Score | Top 1% RMSD | Top 10 Score |
| 2KBY | 25958 | D2 | 4x50 | 154,115,69 | 5.6 | 9.2 | 1.6 | 2.1 | 3.1 | 3.1 | 4.0 | 6.6 |
| 2KO8 | 16492 | C3 | 3x53 | 43,131,125 | 7.7 | 12.0 | 5.3 | 4.6 | 3.9 | 2.4 | 6.2 | 10.1 |
| 2JWK | 15459 | C2 | 2x74 | 263,233,168 | 5.9 | 10.0 | 2.3 | 3.6 | 2.3 | 2.2 | 5.4 | 7.6 |
| 2L01 | - | C2 | 2x77 | 97,223,174 | 5.5 | 9.3 | 3.1 | 2.7 | 1.6 | 1.6 | 5.4 | 8.8 |
| 2LTD | 18469 | C2 | 2x80 | 102,224,188 | 3.3 | 7.0 | 4.7 | 9.6 | 7.0 | 2.6 | 2.6 | 2.6 |
| 2KOD | 16555 | C2 | 2x88 | 134,232,173 | 3.2 | 8.8 | 1.9 | 2.2 | 4.4 | 9.3 | 1.9 | 2.9 |
| 1RJJ | 6011 | C2 | 2x111 | 105,536,359 | 6.6 | 11.6 | 6.1 | 6.0 | 4.3 | 2.7 | 4.7 | 8.3 |
| 2M89 | 19235 | C2 | 2x134 | 92,657,437 | 3.5 | 17.3 | 3.4 | 15.5 | 3.3 | 10.9 | 3.6 | 9.6 |

\* Number of restraints in the following order: RDCs, PCSs, PREs. RDCs were experimental data (**Tab S1**). PCSs and PREs were simulated as described in **Method S1** and **S2**.
<sup>†</sup> The definition of the Top 1% RMSD and Top 10 Score metric is the same as in **Tab 1**. For models generated without paramagnetic NMR restraints, the Rosetta all-atom score was used for ranking, whereas the best-scoring models generated with paramagnetic NMR restraints were identified using the combined NMR score and Rosetta all-atom energy.

Table 2 summarizes the average RMSD_100_ values of the best 1% of models by RMSD and of the ten best-scoring models. Score-vs-RMSD plots of selected proteins are displayed in Fig 5 and for all targets in **Fig S8**. We find a clear improvement in model sampling and scoring with paramagnetic NMR data. The number of cases with a lower RMSD_100_ among the best 1% of models was seven (RDCs), six (PCSs) and seven (PREs) out of eight (considering the symmetric docking results for 1RJJ and 2M89 in which the correct topology could be sampled), and the average RMSD_100_ value over the best 1% of models for all proteins decreased from 5.2 Å without NMR data to 3.6 Å (RDCs), 3.8 Å (PCSs) and 4.2 Å (PREs). The sampling density of low-RMSD models improved considerably for many targets; the number of cases with at least one high-accuracy model (≤ 2 Å) improved from one without NMR data to five, six and two with RDC, PCS and PRE data, respectively. As in our analysis for monomeric proteins, paramagnetic NMR data had a beneficial effect on scoring and convergence. The average enrichment increased from 2.3 to 2.7 (RDCs), 3.1 (PCSs) and 2.4 (PREs) (Fig 5E), and the mean convergence over all targets improved from 5.4 Å to 4.2 Å (RDCs), 2.9 Å (PCSs) and 5.1 Å (PREs) (**Fig S5B**). Using a combination of convergence and NMR Q-factor as criterion to assess prediction reliability, five (RDCs), six (PCSs) and three (PREs) out of eight modeling cases could be classified as successful (**Fig S6B**).

**Figure 5:**
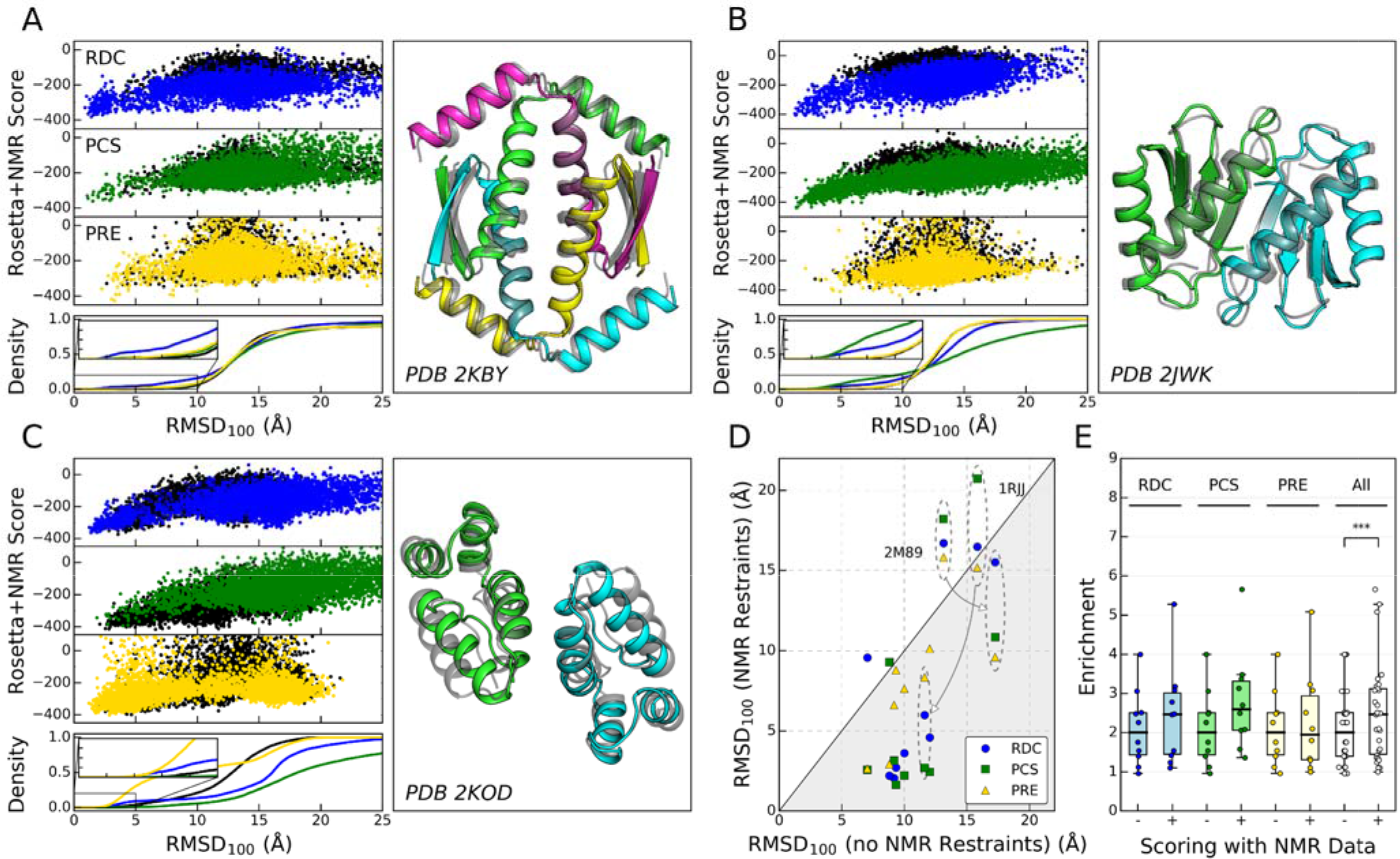
*De novo* structure prediction of symmetric proteins with paramagnetic NMR data. Three selected test cases for symmetric protein structure prediction with experimental RDC and simulated PCS and PRE data: **(A)** tetramerization domain of human p73 (PDB 2KBY), **(B)** periplasmic domain of TolR from H. influenzae, and **(C)** C-terminal domain of HIV-1 capsid protein (PDB 2KOD). Complete results of all symmetric protein targets are given in **Fig S8**. For each protein, the combined Rosetta and paramagnetic NMR score and the cumulative fraction of models versus the models’ RMSD_100_, is shown. The NMR score was either the RDC (blue), PCS (green) or PRE (yellow) score. We compare structure prediction without NMR data (black curve) and using RDCs (blue), PCSs (green) or PREs (yellow), respectively. The lowest-scoring model of 2KBY with RDCs **(A)**, of 2JWK with PCSs **(B)** and of 2KOD with PREs **(C)** is displayed as ribbon diagram for residues in ordered regions (defined in **Tab S1**) and superimposed onto the experimental structure in gray. **(D)** Average RMSD_100_ of the ten best-scoring models for structure prediction with and without paramagnetic NMR data. In the first case, the combined Rosetta and NMR score was used to identify the lowest-scoring models, whereas in the latter case models were ranked by only the Rosetta score. The gray area indicates improvement of RosettaNMR over the unrestrained calculation. The sampling of two proteins for which paramag etic NMR restraints led to a higher RMSD_100_ value, 2M89 and 1RJJ, can be significantly improved by dividing the modeling protocol into two steps: (i) folding of the monomer and (ii) docking of the symmetric dimer. The improvement in the RMSD_100_ is illustrated by gray arrows. **(E)** Improvement in model enrichment by scoring models with an energy function including the respective NMR restraint score compared to the standard Rosetta energy function. Box-and-Whisker plot; comparison was made by two-tailed Wilcoxon signed-rank test (n_All_ = 30, * p < 0.05, ** p < 0.01, *** p < 0.001).

### Pseudocontact shifts are highly beneficial to identify native interfaces in protein-protein docking

PCSs are particularly attractive for protein-protein docking because they combine distance and angular information and are easy to interpret in terms of the coordinates of a nuclear spin within the Δχ-tensor frame. PCSs from multiple tagging sites can remove the ambiguity related to the symmetry of the ΔX-tensor, allowing prediction of the binding interface. The validity of this approach was first corroborated by PCS-based rigid-body docking (37, 38). Later, Schmitz et al. implemented a PCS energy term into the docking program HADDOCK (39). Here, we combined PCSs in RosettaNMR with RosettaDock (29) and show that protein-protein interfaces can be accurately identified using PCS data as the only source of experimental information.

The method performance was assessed on two systems with experimental PCSs (Tab 3): (a) a ternary complex of FKBP12 (FK506-binding protein), rapamycin and the FKBP12-rapamycin-binding (FRB) domain of FRAP (PDB 1FAP) (Fig 6A+B) and (b) a homodimer of the PB1 domain of p62 (PDB 2KTR) (**Fig S9**). For FKBP12-rapamycin-FRB (40), we obtained a best-scoring model with an interface RMSD of 0.5 Å to the X-ray structure (Fig 6A). The final ensemble of ten models was highly converged with an average interface RMSD of 0.6 ± 0.2 Å to the native structure and was in excellent agreement with the PCS data (Q = 0.07). In contrast, docking without PCS data failed to converge; the final ensemble had an average RMSD of 13.2 ± 6.6 Å (Fig 6B).

**Figure 6:**
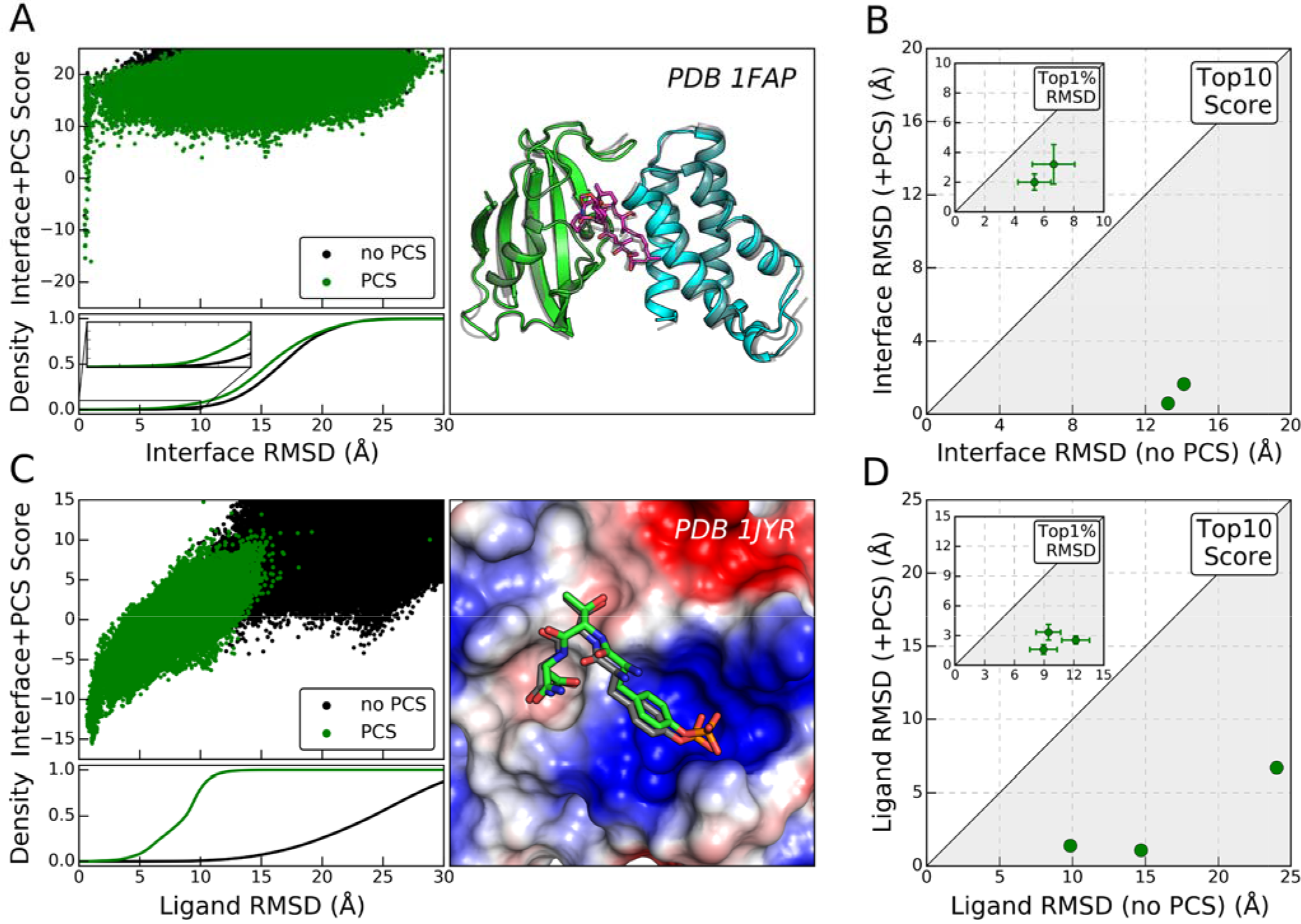
Protein-protein and protein-ligand docking with PCS data. **(A)** Results of docking the FRB domain of FRAP to the FKBP12-rapamycin complex. The combined Rosetta interface and PCS score and the cumulative fraction of models are plotted versus the models’ interface RMSD (PDB 1FAP). The lowest-scoring model obtained with PCS-restrained docking is represented as ribbon diagram with FKBP12 in green, FRB in cyan and rapamycin depicted as magenta sticks, and superimposed onto the X-ray structure colored in gray. **(B)** Average interface RMSD among the ten lowest scoring models obtained for the two docking test cases with or without PCS restraints. The inset compares the average interface RMSD (± S.D.) among the top 1% of models ranked by RMSD. Lower gray triangles indicate RMSD improvement with PCSs. **(C)** Results of docking a pYTN tripeptide ligand to the SH2 domain of Grb2. The combined Rosetta interface and PCS score and the cumulative fraction of models are plotted versus the ligand RMSD. As comparison served the X-ray structure of a phosphorylated peptide with the same three-residue motif bound to Grb2 SH2 (PDB 1JYR). The lowest scoring solution of the ligand obtained with PCS-restrained docking is represented as sticks with atoms colored by chemical identity (C: green, O: red, N: blue, P: orange), and compared with the ligand in the X-ray structure colored gray. **(D)** Average ligand RMSD among the ten lowest-scoring models obtained for the three ligand docking cases without and with PCSs. The inset compares the average interface RMSD (± S.D.) of the best 1% of models ranked by RMSD. Complete results for protein-protein and protein-ligand docking are given in **Fig S9**.

**Table 3:** Structure prediction of protein-protein and protein-ligand complexes with PCS data.

| PDB ID | BMRB ID | Stoichiometry | No. Residues | No. PCS Restraints | no PCS |  | PCS |  |
| --- | --- | --- | --- | --- | --- | --- | --- | --- |
|  |  |  |  |  | Top 1% RMSD<sup>†</sup> | Top 10 Score<sup>‡</sup> | Top 1% RMSD | Top 10 Score |
| Protein Docking |  |  |  |  |  |  |  |  |
| 2KTR | 16361 | 1:1 | 100+100 | 1042 | 5.3 | 14.1 | 2.0 | 1.7 |
| 1FAP | 11471 | 1:1 | 107+95 | 404 | 6.6 | 13.2 | 3.2 | 0.6 |
| Ligand Docking |  |  |  |  |  |  |  |  |
| 1JYR | 11055 | 1:1 | 100+1 | 227+15* | 8.9 | 14.7 | 1.6 | 1.1 |
| 1X0N | 11055 | 1:1 | 100+1 | 227+55 | 9.4 | 9.8 | 3.3 | 1.4 |
| 3U1I | - | 1:1 | 247+1 | 223+18 | 12.2 | 24.0 | 2.6 | 6.7 |

\* Number of PCSs for protein and ligand.
<sup>†</sup> Average RMSD of best 1% models ranked by RMSD.
<sup>‡</sup> Average RMSD of ten best-scoring models. The interface and ligand RMSD as defined in Methods was used as similarity measure for protein and ligand docking, respectively. Models were ranked by their combined Rosetta energy and PCS restraint score.

For the p62 PB1 homodimer (41), docking started from the unbound conformation of the p62 PB1 because no high-resolution structure of the dimer was available and the experimental structure (PDB 2KKC) was reported as rigid-body docking structure. Although docking of unbound structures is notoriously difficult, our calculations converged to a best-scoring model with an interface RMSD of 1.3 Å (**Fig S9**) compared to the published model by Saio et al. that is very similar to the X-ray structure of a related p62-PKCζ heterodimer (PDB 4MJS). The final ensemble of ten lowest-scoring models had an average RMSD of 1.7 ± 0.3 Å and was in very good agreement with the experimental PCSs with a Q-factor of 0.17. In contrast, docking without PCSs failed to converge and the interface RMSD of the best ten models was 14.1 ± 2.4 Å (Fig 6B) (Q = 0.21).

### Pseudocontact shifts efficiently guide protein-ligand docking to native-like complexes

Paramagnetic NMR has been successful in compound screening for drug design (42) and in modeling of protein-ligand interactions (43). PCS restraints play a special role because they can be measured for ligands with a broad range of affinities and provide atomic detail information about the binding mode. Here, we have tested the performance of RosettaNMR with RosettaLigand (30) to leverage PCSs for modeling protein-ligand interactions. Using PCSs as the only source of experimental information for three protein-ligand systems with different affinities and no prior knowledge about where the ligand binds, our method was able to recover the native binding mode with an accuracy of up to 1.1 Å ligand RMSD.

We employed a two-step protocol for global protein-ligand docking. A search of the ligand’s six rigid body DOFs in a grid around the protein identifies positions with low PCS score, which are then used as input into RosettaLigand. For our three test cases, we found that after the initial grid search the ligand clustered in a space corresponding to an RMSD of around 6 to 8 Å relative to the native ligand binding mode. Simultaneous ligand translation, rotation and conformer replacement further explores the space around the starting position, and subsequent high-resolution docking refines the structure to atomic detail. Furthermore, we implemented a PCS scoring grid (44) into RosettaLigand, which evaluates the ligand position given the experimental PCS data and a Δχ-tensor as input. The components of the Δχ-tensor(s) can be calculated from the proteins’ PCS values prior to docking.

We tested this method on three protein-ligand complexes (Tab 3): the SH2 domain of Grb2 bound to a low-affinity phosphorylated pYTN tripeptide ligand as well as a high-affinity macrocyclic inhibitor (45), and a complex of dengue virus NS2B-NS3 protease (DENpro) with a high-affinity ligand (46). PCS-assisted docking of the two SH2 ligands was converged (see score-vs-RMSD plots in Fig 6C and **Fig S9**). The models of the pYTN ligand (Fig 6C) and the macrocyclic inhibitor (**Fig S9**) with the lowest Rosetta interface and PCS score had a ligand RMSD of 1.1 Å in both cases, and were in very good agreement with the NMR data (Q = 0.14, Q = 0.23). The average RMSD of the ten best models was 1.1 ± 0.1 Å and 1.4 ± 0.6 Å, respectively (Fig 6D). In contrast, ligand docking without PCSs failed to identify the binding pocket (ligand RMSDs were 14.7 ± 6.9 Å and 9.8 ± 4.5 Å). Similar improvements were found for the third system, DENpro with a high-affinity ligand. Our lowest scoring model had a PCS Q-factor of 0.11 and a RMSD of 2.1 Å to the model by Chen et al. (**Fig S9**) which bears high similarity to the X-ray structure of a related peptide-DENpro complex (PDB 3U1I) (47). Due to the presence of a second low-scoring binding mode (**Fig S9**), the average RMSD of the ten best models (6.7 ± 2.3 Å) was larger and less converged than for the other two test cases, but still considerably lower than for docking without PCSs (24.0 ± 8.1 Å) (Fig 6D).

## Discussion

The utility of paramagnetic NMR restraints has long been recognized (16) but they are often too sparse to allow structure determination from these data exclusively. Moreover, interpretation of PCS and RDC data usually requires that the alignment tensor or metal Δχ-tensor is calculated from an existing structural model and simultaneous computations of the tensor and structure (48) have remained difficult or are practical only for small proteins. Consequently, PCSs, RDCs and PREs have mainly been used in addition to NOEs in structure calculations or refinement (49, 50). However, computational methods can leverage limited experimental data, and integrative modeling approaches have proven successful in assisting structure determination of larger proteins from limited NMR data (51, 52).

We present a strategy for protein structure modeling from paramagnetic restraints as part of the Rosetta biomolecular modeling suite. Our approach revives and extends the RosettaNMR framework and allows modeling from long-range PCS, RDC and PRE restraints, in combination with CSs and NOEs. RosettaNMR was designed to be integrable with other modeling protocols such as RosettaAbinitio, RosettaDock, RosettaLigand and RosettaSymmetry in order to tackle various modeling challenges for proteins and protein interactions.

Structures predicted by our method yield comparable accuracy as previous CS- (21), PCS- (26) and CS-RDC-Rosetta (22) protocols and improved accuracy compared to BCL::Fold (4) (**Tab S5**). Our analysis shows that paramagnetic restraints greatly enrich for near-native models by improving both model sampling and scoring. When evaluated on the monomeric protein test set, paramagnetic restraints allowed building more accurate models for 85% of the proteins. Compared to CS-Rosetta calculations on the same protein set, paramagnetic restraints yielded lower RMSD_100_ in 72% of the cases. Combining CSs with paramagnetic restraints sampled at least one high-accuracy model (RMSD_100_ ≤ 2 Å) in 50% of the cases, and a medium-accuracy model (RMSD_100_ ≤ 3 Å) for 75% of the proteins. In contrast, calculations without NMR data rarely arrived at high-accuracy models; for only 9% of the proteins a model ≤ 2 Å could be generated. This clearly illustrates that CSs and paramagnetic restraints complement each other in protein structure determination: CSs guide the fragment search through local conformation bias while paramagnetic restraints guide the global assembly into a 3D fold.

Our results for *de novo* protein structure prediction are further confirmed by those obtained on the symmetric protein benchmark set and for modeling protein-protein and protein-ligand complexes. Paramagnetic restraints improve model quality of up to 80% of the symmetric proteins and help to recapitulate native protein and ligand binding poses with an RMSD as low as 1 Å. Moreover, paramagnetic restraints proved highly beneficial for model selection. Energy differences between native-like and non-native conformations can be small due to a simplified protein representation in Rosetta and inaccuracies in the energy function, making identification of the correct fold difficult. Supplementing the Rosetta energy function with a score reporting the PCS, RDC and/or PRE restraint violation significantly improves model enrichment by a factor of ~2.

Predicting the likelihood of success for structure prediction and refinement remains difficult, as various aspects (fold complexity, secondary structure content, number and quality of restraints and spin-label sites) play an important, and protein-specific, role. In general, paramagnetic restraints from multiple tagging sites are expected to improve structure prediction by resolving the degeneracy of solutions that equally fulfill the experimental data. For proteins 1D3Z and 1G7D in our benchmark set which had PCS data from multiple tagging sites available the largest RMSD improvements were found after combining all datasets, and individual tagging sites were found not equally valuable in sampling low-RMSD models.

Additional improvements to integrative modeling with sparse paramagnetic NMR data in Rosetta are expected to come from enhanced sampling algorithms such as iterative fragment assembly (53), iterative hybridization (35) or RASREC (54). These techniques can improve modeling for larger and more complex topologies and make more efficient use of NMR data. Further improvements can be expected from incorporation of alternative experimental data, for instance from SAXS (55) or mass-spectroscopy (56) or from co-evolution data (57). Evolutionary coupling-NMR (58) incorporates evolutionary contacts during and after NMR data interpretation and NOE assignment. With the expansion of genomic databases and increased accuracy in protein contact prediction algorithms, evolutionary constraints provide increasingly viable orthogonal structural information to assist NMR-guided protein structure prediction.

In conclusion, we present the development and application of a structural biology framework for protein modeling from sparse paramagnetic NMR restraints as part of the Rosetta software suite. The improved RosettaNMR framework integrates the capabilities of CS-Rosetta and PCS-Rosetta and allows the use of all combinations of CS, NOE, PCS, RDC, and PRE restraints. We applied RosettaNMR to structure prediction of monomeric and symmetric proteins, protein-protein docking and protein-ligand docking. Our approach enables structure determination from sparse paramagnetic NMR datasets where other types of data are difficult or time-consuming to acquire. Approaches like these will open the door for structure determination of complex biomolecular assemblies and ultimately our understanding of their biological function.

## Supporting information

RosettaNMR Protocol Capture

## Supplemental Information

Supplemental information includes a detailed description of the PCS, RDC and PRE scoring methods, eleven supporting figures and five supporting tables (**File S1**), and a protocol capture on RosettaNMR with complete command lines (**File S2**) as well as input files for running the protocol capture (**File S3**).

## Acknowledgements

This work was supported by NIH grant R01 GM080403. GK was supported by fellowships from the German Research Foundation (KU 3510/1-1) and the American Heart Association (18POST34080422). JKL and RB were funded by the Flatiron Institute as part of the Simons Foundation. This work was conducted using the resources of the Advanced Computing Center for Research and Education (ACCRE) at Vanderbilt University. The authors also like to thank the members of the RosettaCommons for their helpful discussions and support.

## Author Contributions

GK, JKL and JM conceived the idea and GK and JKL designed the framework. GK wrote the code and performed the calculations and analyzed the data with guidance from JKL and JM. GK and JKL wrote the paper with input from RB and JM. The final manuscript has been approved by all authors.

## Methods

### Selection of benchmark proteins

For *de novo* structure prediction, 28 monomeric and eight symmetric proteins with experimental NMR data were selected (**Tab S1**). The monomeric protein benchmark set included 16 proteins with RDC data, six proteins with PCS data, three proteins with PRE data and three proteins for which both PCS and RDC data were available. Because only a small number of proteins with paramagnetically-induced RDC data are reported in the literature to the best of our knowledge, we included additional proteins with RDC data induced by conventional alignment media; their computational treatment is identical. In addition, CSs were available for all proteins. These included CSs of backbone atoms ^1^H^N^, ^1^H^α^, ^13^C^α^, 13C’, ^13^C^β^, ^15^N^H^ for almost all proteins. ^13^C’ shifts were not available for proteins 2H45, 2A7O, 1CMZ, 5T1N, 1X0N and 1RJV, ^13^C^β^ shifts were missing for 1G7D and 1CDL and ^1^H^α^ shifts were absent for 2K61 and 1J54.

To compare with previous structure prediction studies, our benchmark set included 14 targets from the BCL::Fold benchmark (4), seven targets from previous CS-Rosetta benchmarks (21, 22) and 4 targets from the PCS-Rosetta benchmark (26). Furthermore, proteins were selected to have a diverse set of α, β and α/β topologies, and ranged from 56 to 199 residues.

The symmetric protein benchmark set comprised six targets with C2 symmetry, one target with C3 symmetry and one target with D2 symmetry. All targets had experimental RDC data, and PCSs and PREs were simulated (**Method S1+S2**). Three targets were taken from an earlier CS-RDC-Rosetta study (59).

### Calculation of the paramagnetic NMR score

PCSs, RDCs and PREs are evaluated by a Rosetta scoring method in low- and high-resolution mode and computed on the whole protein structure. The score represents the sum of squared errors between the experimental and predicted NMR values. Computation of each of the three score terms is described in detail in **Method S3**. Detailed description of the format of PCS, RDC and PRE input files, the choice of critical parameters and complete command lines are provided with the protocol capture in **File S2**.

Analogous to previous Rosetta protocols for NMR structure determination (21, 22, 26), we decided to add each paramagnetic NMR score to the Rosetta energy function using a different weight (*w_ParaNMR_*), which was adjusted such that the ranges of the NMR and Rosetta score were approximately equal. The weight was determined by first generating 1000 models without NMR data and then rescoring them with the respective paramagnetic NMR restraints. The weight *w_ParaNMR_* was calculated as

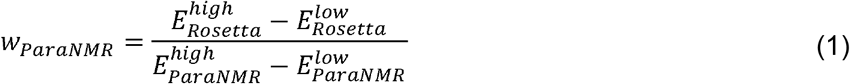

where 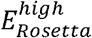 and 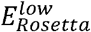 are the average of highest and lowest 10% of the values of the Rosetta score, and 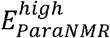 and 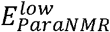 are the average of the highest and lowest 10% of the values of the respective paramagnetic NMR score, respectively. For *de novo* structure prediction, *E_Rosetta_* represented the Rosetta low-resolution score which is obtained with the score3 weight set. For scoring all-atom models, *E_Rosetta_* was calculated with the Rosetta ref2015 energy function (60). In case *E_ParaNMR_* comprised more than one type of paramagnetic NMR data, *w_ParaNMR_* was split between those, and we chose a ratio proportional to the logarithm of the number of restraints in each data set.

### RosettaNMR de novo structure prediction with paramagnetic and chemical shift NMR data

Structure prediction was accomplished using Rosetta’s fragment assembly protocol (28) followed by all-atom refinement via FastRelax (61). Paramagnetic NMR data were used as follows: CSs for fragment selection, PCS, RDC and PRE data for low-resolution fragment assembly, high-resolution refinement, model scoring and final model selection.

The fragment search was carried out with the Rosetta3 fragment picker (62) and two libraries were generated for each protein. For the first library, only sequence-based secondary structure information derived from PSIPRED (63) and Jufo9D (64) was used. The second library was created using backbone CSs and TALOS+ (65) secondary structure assignments. In all cases, homologous proteins were excluded according to a sequence similarity criterion (PSI-BLAST E-value < 0.5).

To guide fragment assembly with RDCs, PCSs or PREs, the score function was supplemented with the score term for the respective paramagnetic NMR restraint, using a weighting factor *w_ParaNMR_* optimized against Rosetta’s low-resolution score as described above. Following high-resolution refinement, models were rescored with the Rosetta all-atom energy function (60) combined with the respective paramagnetic NMR score term.

For each benchmark target, 10,000 all-atom models were generated, and computations took 0.25 – 0.5 hrs/model for RDCs, 0.5 – 1.0 hrs/model for PREs and 0.5 – 2.0 hrs/model for PCSs.

The performance of RosettaNMR structure prediction was evaluated by calculating model RMSD_100_ to the native structure, model convergence and NMR Q-factor (see **Method S4**). The effect of the PCS, RDC and PRE score on model scoring was assessed using the enrichment metric.

### RosettaNMR symmetric protein modeling with paramagnetic NMR data

Structure prediction of symmetric proteins was accomplished using the Fold-and-Dock protocol (36) which carries out simultaneous folding and docking of multi-chain homo-oligomers. We adjusted the Fold-and-Dock protocol to include RDC, PCS and PRE data for fragment assembly, rigid body sampling and final model scoring and selection.

In a symmetric complex, the PCSs and PREs are calculated as the sum over equivalent spins in symmetric subunits. For evaluating RDCs, we expect that one axis of the alignment tensor is collinear with the symmetry axis of the system. Thus, a copy of the RDC values is assigned to each subunit, and simultaneous search of all rigid body DOFs finds the orientation of the alignment tensor that best aligns with the symmetry axis.

We used published RDCs (**Tab S1**) and simulated PCSs and PREs (**Method S1+S2**). The type of symmetry was assumed to be known. For unknown symmetry groups, all eligible point groups can be tested under the assumption that models with correct symmetry will score better with higher convergence.

A total of 5,000 models were generated for each protein, which took only slightly longer than structure prediction for monomeric proteins of similar sizes. The respective paramagnetic NMR score was computed for all models, multiplied by the weighting factor *w_ParaNMR_* and added to the Rosetta all-atom score for final model selection.

An additional two-step protocol (59) was conducted for proteins 1RJJ and 2M89, for which Rosetta failed to produce models with RMSD_100_ < 10 Å. The structure of the monomer was first modeled without NMR data using *de novo* prediction and subsequently docked together using RDC, PCS or PRE data. An ensemble of the 50 lowest-scoring monomer models was used as starting conformations for symmetric docking (31). This step yielded a total of 50,000 models which were ranked by the Rosetta all-atom interface score together with their paramagnetic NMR score.

### RosettaNMR protein-protein docking guided by PCS data

We adapted RosettaDock (29) to use PCS data both for rigid body sampling and high-resolution refinement. The PCS score for every tagging site is minimized over all intra- and intermolecular PCSs to yield the Δχ-tensor with the best fit to a given docking pose. Thus, incorrect binding poses with a large number of PCS violations are highly penalized. Moreover, rigid body DOFs and backbone torsion angles are optimized during high-resolution refinement under the restraint of the PCS target function.

PCS-guided Rosetta docking was tested for two protein complexes with experimental PCS data: a p62 PB1 homodimer (PDB 2KTR) and a ternary complex of FKBP12 (FK506-binding protein), rapamycin and the FKBP12-rapamycin-binding (FRB) domain of FRAP (PDB 1FAP). RosettaDock calculations were performed as global docking with random initial orientations of the two binding partners at the beginning of each simulation. The magnitude of translational and rotational rigid body Monte Carlo moves was set to 0.3 Å and 5°, respectively. For the ternary FKBP12-rapamycin-FRB complex, the rapamycin ligand was kept fixed with respect to FKBP12 and the FRB domain was docked to the FKBP12-rapamycin complex. Simultaneous optimization of protein side-chain rotamers and rapamycin conformers during high-resolution docking allowed equal adjustment of all protein-protein and protein-ligand interfaces.

A total of 100,000 models were generated for each target with and without PCS data and final models were selected by their combined Rosetta interface and PCS score. Model accuracy was evaluated by the RMSD of all backbone heavy atoms in the interface after superposition. The interface was defined as all residues within 8 Å of any other residue on the other protein binding partner.

### RosettaNMR protein-ligand docking guided by PCS data

To explore the full potential of PCSs for ligand docking without prior knowledge of the ligand binding site, we made the following adaptations to the RosettaLigand protocol (30): First, to identify ligand positions with low PCS score, an additional pre-transform step runs a grid search of the ligand’s rigid body DOFs in a shell around the protein (with adjustable resolution; 5 Å and 20° in this study). These positions are then used as starting points for low-resolution docking to find a favorable non-clashing ligand position in the binding pocket. For comparison, docking without PCS data was initiated from positions uniformly distributed in a shell around the protein with a width corresponding to half the ligand neighbor radius and a spacing of 5 Å. Second, the cartesian grid-based energy function that is used for ligand scoring in the low-resolution stage (44) was extended by a new PCS scoring grid, which represents the 3D field of the expected ligand PCS given a Δχ-tensor as input. Thus, the PCS ligand grid score examines the agreement of the ligand position with the localization space defined by the PCS. The Δχ-tensor components can be determined from the protein PCS dataset. The dimensions of the grid were set to 30 Å × 30 Å × 30 Å with a spacing of 0.25 Å. Initially, the grid is centered at the position of the ligand neighbor atom but is recomputed if the ligand moves outside of the grid.

After low-resolution docking the algorithm proceeds with its high-resolution phase using the Rosetta all-atom energy function including the PCS score. Six cycles of alternating protein side-chain and ligand conformer packing and small perturbations to the ligand are performed, followed by a final gradient minimization of the interface. A total of 100,000 models were generated, and final models were selected by their combined interface and PCS score. Model accuracy was assessed by the RMSD of all ligand heavy atoms without superposition.

We applied PCS-guided RosettaLigand to three systems for which experimental PCS data had been published: two complexes of the Src homology 2 (SH2) domain of the growth factor receptor-bound protein 2 (Grb2) with a low-affinity pYTN tripeptide and a high-affinity inhibitor (4-[(10S,14S,18S)-18-(2-amino-2-oxoethyl)-14-(1-naphthylmethyl)-8,17,20-trioxo-7,16,19-triazaspiro[5.14]icos-11-en-10-yl]benzylphosphonic acid) (45), and a complex between the dengue virus NS2B-NS3 protease with a high affinity ligand (46).

## References

1. Leaver-Fay A, et al. (2011) ROSETTA3: an object-oriented software suite for the simulation and design of macromolecules. Methods Enzymol 487:545–574.

2. Dominguez C, Boelens R, & Bonvin AM (2003) HADDOCK: a protein-protein docking approach based on biochemical or biophysical information. J Am Chem Soc 125(7):1731–1737.

3. Russel D, et al. (2012) Putting the pieces together: integrative structure determination of macromolecular assemblies. PLoS Biol 10(1):e1001244.

4. Weiner BE, et al. (2014) BCL::Fold--protein topology determination from limited NMR restraints. Proteins 82(4):587–595.

5. Demers JP, et al. (2014) High-resolution structure of the Shigella type-III secretion needle by solid-state NMR and cryo-electron microscopy. Nat Commun 5:4976.

6. Robinson PJ, et al. (2015) Molecular architecture of the yeast Mediator complex. Elife 4.

7. LeMaster DM (1990) Deuterium labelling in NMR structural analysis of larger proteins. Q Rev Biophys 23(2):133–174.

8. Pervushin K, Riek R, Wider G, & Wuethrich K (1997) Attenuated T2 relaxation by mutual cancellation of dipole–dipole coupling and chemical shift anisotropy indicates an avenue to NMR structures of very large biological macromolecules in solution. Proc. Natl. Acad. Sci. U. S. A. 94:12366–12371.

9. Tugarinov V & Kay LE (2005) Methyl groups as probes of structure and dynamics in NMR studies of high-molecular-weight proteins. Chembiochem 6(9):1567–1577.

10. Matzapetakis M, Turano P, Theil EC, & Bertini I (2007) 13C- 13C NOESY spectra of a 480 kDa protein: solution NMR of ferritin. J Biomol NMR 38(3):237–242.

11. Paramasivam S, et al. (2012) Enhanced sensitivity by nonuniform sampling enables multidimensional MAS NMR spectroscopy of protein assemblies. J Phys Chem B 116(25):7416–7427.

12. Graham B, et al. (2011) DOTA-amide lanthanide tag for reliable generation of pseudocontact shifts in protein NMR spectra. Bioconjug Chem 22(10):2118–2125.

13. Loh CT, et al. (2013) Lanthanide tags for site-specific ligation to an unnatural amino acid and generation of pseudocontact shifts in proteins. Bioconjug Chem 24(2):260–268.

14. Crick DJ, et al. (2015) Integral membrane protein structure determination using pseudocontact shifts. J Biomol NMR 61(3–4):197–207.

15. Rathinavelan T, et al. (2014) NMR model of PrgI-SipD interaction and its implications in the needle-tip assembly of the Salmonella type III secretion system. J Mol Biol 426(16):2958–2969.

16. Koehler J & Meiler J (2011) Expanding the utility of NMR restraints with paramagnetic compounds: background and practical aspects. Prog Nucl Magn Reson Spectrosc 59(4):360–389.

17. Bowers PM, Strauss CEM, & Baker D (2000) Denovo protein structure determination using sparse NMR data. J. Biomol. NMR 18:311–318.

18. Rohl CA & Baker D (2002) De novo determination of protein backbone structure from residual dipolar couplings using Rosetta. J Am Chem Soc 124(11):2723–2729.

19. Meiler J & Baker D (2003) Rapid protein fold determination using unassigned NMR data. Proc Natl Acad Sci U S A 100(26):15404–15409.

20. Meiler J (2003) PROSHIFT: Protein Chemical Shift Prediction Using Artificial Neural Networks. J. Biomol. NMR 26(1):25–37.

21. Shen Y, et al. (2008) Consistent blind protein structure generation from NMR chemical shift data. Proceedings of the National Academy of Sciences of the United States of America 105(12):4685–4690.

22. Raman S, et al. (2010) NMR structure determination for larger proteins using backbone-only data. Science 327(5968):1014–1018.

23. Lange OF, et al. (2012) Determination of solution structures of proteins up to 40 kDa using CS-Rosetta with sparse NMR data from deuterated samples. Proc Natl Acad Sci U S A 109(27):10873–10878.

24. Reichel K, et al. (2017) Systematic evaluation of CS-Rosetta for membrane protein structure prediction with sparse NOE restraints. Proteins 85(5):812–826.

25. Shen Y & Bax A (2015) Homology modeling of larger proteins guided by chemical shifts. Nat Methods 12(8):747–750.

26. Schmitz C, Vernon R, Otting G, Baker D, & Huber T (2012) Protein structure determination from pseudocontact shifts using ROSETTA. J Mol Biol 416(5):668–677.

27. Yagi H, et al. (2013) Three-dimensional protein fold determination from backbone amide pseudocontact shifts generated by lanthanide tags at multiple sites. Structure 21(6):883–890.

28. Simons KT, Kooperberg C, Huang E, & Baker D (1997) Assembly of Protein Tertiary Structures from Fragments with Similar Local Sequences using Simulated Annealing and Bayesian Scoring Functions. J. Mol. Biol. 268:209–225.

29. Gray JJ, et al. (2003) Protein-protein docking with simultaneous optimization of rigid-body displacement and side-chain conformations. J Mol Biol 331(1):281–299.

30. Meiler J & Baker D (2006) ROSETTALIGAND: protein-small molecule docking with full side-chain flexibility. Proteins 65(3):538–548.

31. Andre I, Bradley P, Wang C, & Baker D (2007) Prediction of the structure of symmetrical protein assemblies. Proc Natl Acad Sci U S A 104(45):17656–17661.

32. Iwahara J, Schwieters CD, & Clore GM (2004) Ensemble approach for NMR structure refinement against ^1^H paramagnetic relaxation enhancement data arising from a flexible paramagnetic group attached to a macromolecule. J Am Chem Soc 126(18):5879–5896.

33. Chaudhury S, Lyskov S, & Gray JJ (2010) PyRosetta: a script-based interface for implementing molecular modeling algorithms using Rosetta. Bioinformatics 26(5):689–691.

34. Fleishman SJ, et al. (2011) RosettaScripts: A Scripting Language Interface to the Rosetta Macromolecular Modeling Suite. PLoS One 6(6):e20161.

35. Ovchinnikov S, et al. (2017) Protein structure determination using metagenome sequence data. Science 355(6322):294–298.

36. Das R, et al. (2009) Simultaneous prediction of protein folding and docking at high resolution. Proc Natl Acad Sci U S A 106(45):18978–18983.

37. Ubbink M, Ejdeback M, Karlsson BG, & Bendall DS (1998) The structure of the complex of plastocyanin and cytochrome f, determined by paramagnetic NMR and restrained rigid-body molecular dynamics. Structure 6(3):323–335.

38. Pintacuda G, Park AY, Keniry MA, Dixon NE, & Otting G (2006) Lanthanide labeling offers fast NMR approach to 3D structure determinations of protein-protein complexes. J Am Chem Soc 128(11):3696–3702.

39. Schmitz C & Bonvin AM (2011) Protein-protein HADDocking using exclusively pseudocontact shifts. J Biomol NMR 50(3):263–266.

40. Kobashigawa Y, et al. (2012) Convenient method for resolving degeneracies due to symmetry of the magnetic susceptibility tensor and its application to pseudo contact shift-based protein-protein complex structure determination. J Biomol NMR 53(1):53–63.

41. Saio T, Yokochi M, Kumeta H, & Inagaki F (2010) PCS-based structure determination of protein-protein complexes. J Biomol NMR 46(4):271–280.

42. Jahnke W (2002) Spin labels as a tool to identify and characterize protein-ligand interactions by NMR spectroscopy. Chembiochem 3(2-3):167–173.

43. Guan JY, et al. (2013) Small-molecule binding sites on proteins established by paramagnetic NMR spectroscopy. J Am Chem Soc 135(15):5859–5868.

44. DeLuca S, Khar K, & Meiler J (2015) Fully Flexible Docking of Medium Sized Ligand Libraries with RosettaLigand. PLoS One 10(7):e0132508.

45. Saio T, et al. (2011) An NMR strategy for fragment-based ligand screening utilizing a paramagnetic lanthanide probe. J Biomol NMR 51(3):395–408.

46. Chen WN, et al. (2016) Sensitive NMR Approach for Determining the Binding Mode of Tightly Binding Ligand Molecules to Protein Targets. J Am Chem Soc 138(13):4539–4546.

47. Noble CG, Seh CC, Chao AT, & Shi PY (2012) Ligand-bound structures of the dengue virus protease reveal the active conformation. J Virol 86(1):438–446.

48. Bryson M, Tian F, Prestegard JH, & Valafar H (2008) REDCRAFT: a tool for simultaneous characterization of protein backbone structure and motion from RDC data. J Magn Reson 191(2):322–334.

49. Gaponenko V, et al. (2004) Improving the accuracy of NMR structures of large proteins using pseudocontact shifts as long-range restraints. J Biomol NMR 28(3):205–212.

50. Gabel F, et al. (2008) A structure refinement protocol combining NMR residual dipolar couplings and small angle scattering restraints. J Biomol NMR 41(4):199–208.

51. Baldwin AJ, et al. (2011) The polydispersity of alphaB-crystallin is rationalized by an interconverting polyhedral architecture. Structure 19(12):1855–1863.

52. Loquet A, et al. (2012) Atomic model of the type III secretion system needle. Nature 486(7402):276–279.

53. Blum B, Jordan MI, & Baker D (2010) Feature space resampling for protein conformational search. Proteins 78(6):1583–1593.

54. Lange OF & Baker D (2012) Resolution-adapted recombination of structural features significantly improves sampling in restraint-guided structure calculation. Proteins: Structure, Function, and Bioinformatics 80(3):884–895.

55. Rossi P, et al. (2015) A hybrid NMR/SAXS-based approach for discriminating oligomeric protein interfaces using Rosetta. Proteins 83(2):309–317.

56. Aprahamian ML, Chea EE, Jones LM, & Lindert S (2018) Rosetta Protein Structure Prediction from Hydroxyl Radical Protein Footprinting Mass Spectrometry Data. Anal Chem 90(12):7721–7729.

57. Hopf TA, et al. (2014) Sequence co-evolution gives 3D contacts and structures of protein complexes. Elife 3.

58. Tang Y, et al. (2015) Protein structure determination by combining sparse NMR data with evolutionary couplings. Nat Methods 12(8):751–754.

59. Sgourakis NG, et al. (2011) Determination of the structures of symmetric protein oligomers from NMR chemical shifts and residual dipolar couplings. J Am Chem Soc 133(16):6288–6298.

60. Alford RF, et al. (2017) The Rosetta All-Atom Energy Function for Macromolecular Modeling and Design. J Chem Theory Comput 13(6):3031–3048.

61. Tyka MD, et al. (2011) Alternate states of proteins revealed by detailed energy landscape mapping. J Mol Biol 405(2):607–618.

62. Vernon R, Shen Y, Baker D, & Lange OF (2013) Improved chemical shift based fragment selection for CS-Rosetta using Rosetta3 fragment picker. J Biomol NMR 57(2):117–127.

63. Jones DT (1999) Protein Secondary Structure Prediction Based on Position-specific Scoring Matrices. J. Mol. Biol. 292(2):195–202.

64. Koehler J, Woetzel N, Staritzbichler R, Sanders CR, & Meiler J (2009) A unified hydrophobicity scale for multispan membrane proteins. Proteins 76(1):13–29.

65. Shen Y, Delaglio F, Cornilescu G, & Bax A (2009) TALOS+: a hybrid method for predicting protein backbone torsion angles from NMR chemical shifts. J Biomol NMR 44(4):213–223.

