## Supplemental Information for "Integrative protein modeling in RosettaNMR from sparse paramagnetic restraints"

### Supporting Methods

#### Method S1: Simulation of PCS data for symmetric proteins

For each symmetric protein, two spin-label sites were chosen focusing on solvent-exposed residues in the middle of secondary structure elements because it was assumed that those sites infer a reduced flexibility on the spin-label and have limited effect on protein stability which would render them possible candidates for experimental lanthanide ion tagging. The lanthanide ion was placed 6 Å away from the Cβ atom of the selected residue in the direction of the Cα-Cβ bond vector, consistent with the approximate side chain length of small lanthanide binding tags. $^1H^N$ PCS data were generated for two lanthanides ($Tb^{3+}$, $Tm^{3+}$) using Δχ-tensors with magnitudes commonly found in the literature (1) and random values for Euler angles α, β and γ. A uniform random error in the range of ± 0.1 ppm was added and all residues with a magnitude of the PCS value greater than 2.0 ppm were excluded to account for the fact that large PCS effects of residues close to the metal ion are usually not detected in an experimental setting because the PRE effect broadens their NMR signal beyond detection.

#### Method S2: Simulation of PRE data for symmetric proteins

Two spin-label sites per symmetric protein were selected using the same criteria as for simulating PCSs, i.e. a solvent-exposed residue located in a continuous stretch of secondary structure. In addition, protein 2M89 offers a solvent-exposed cysteine residue which was also selected as a spin-label site. $^1H^N$-$\Gamma_2$ PREs were simulated by placing all distinct, non-clashing side-chain conformations of the MTSL residue at the designated spin-label site and assigning them a weight proportional to their Boltzmann-weighted rotamer energy. The PRE was calculated from this spin-label ensemble using a modified version of the PRE equations for dipole-dipole relaxation (2) as well as theoretical values of s, g and $\tau_e$ appropriate for a nitroxide radical (3), and values of 14.1 T and 298 K for $B_0$ and T. An internal correlation time of $10^{-3}$ ns of the spin-label was assumed and the rotational correlation time of the protein was estimated from its molecular mass using the Stokes-Einstein relationship. A uniform random error in the range between 0-20 Hz was added and all residues with a PRE rate > 200Hz (this amounts to an approximate $^1H^N$-radical distance of < 10-12 Å) were excluded, to account for the fact that NMR signals of residues close to the spin-label are usually undetectable because their NMR line is broadened beyond detection.

#### Method S3: Calculation of the PCS, RDC and PRE score

***Rosetta PCS Score:*** The PCS (in ppm) induced by a paramagnetic metal ion *M* on a nuclear spin *i* in the molecular frame *f* can be calculated as:

$$\delta_i^{PCS} = \frac{1}{4\pi r^3} \cdot \left[ (\Delta\chi_{zz} - \overline{\Delta\chi}) \cdot \frac{2z_i^2 - x_i^2 - y_i^2}{2r_{ij}^2} + (\Delta\chi_{xx} - \Delta\chi_{yy}) \cdot \frac{x_i^2 - y_i^2}{r_{ij}^2} + \Delta\chi_{xy} \cdot \frac{2x_i y_i}{r_i^2} + \Delta\chi_{xz} \cdot \frac{2x_i z_i}{r_i^2} + \Delta\chi_{yz} \cdot \frac{2y_i z_i}{r_i^2} \right], \text{with } \overline{\Delta\chi} = Tr(\Delta\chi)/3 \qquad \text{(S1)}$$

where $r_i$ is the distance between the spin *i* and the metal ion *M*, $x_i$, $y_i$, $z_i$ are the cartesian coordinates of the metal-spin connection vector, and $\Delta\chi_{xx}$, $\Delta\chi_{yy}$, $\Delta\chi_{zz}$, $\Delta\chi_{xy}$, $\Delta\chi_{xz}$ and $\Delta\chi_{yz}$ are the components of the magnetic susceptibility tensor, Δχ, in the frame *f* (as $\Delta\chi_{zz} = -\Delta\chi_{xx} - \Delta_{xy}$, there are only five independent Δχ-tensor values). The PCS score, $E_{PCS}$, over all paramagnetic centers $M_k$ and spins *i* is calculated as:

$$E_{PCS} = \sum_k \sqrt{\sum_i \left(\delta_i^{PCS,calc}(M_k) - \delta_i^{PCS,exp}(M_k)\right)^2} \qquad \text{(S2)}$$

where $\delta_i^{PCS,calc}(M_k)$ and $\delta_i^{PCS,exp}(M_k)$ are the calculated and experimental PCS values of spin *i* induced by metal ion $M_k$. For calculating $E_{PCS}$, we have adapted a computational procedure previously described for the original PCS-Rosetta algorithm (4). At each scoring step, the Δχ-tensor is determined by singular value decomposition (SVD) along with a grid search over the cartesian coordinates $x_M$, $y_M$, $z_M$ to find the metal ion position. The node with the lowest value of $E_{PCS}$ is then used as starting point for a full optimization of the metal coordinates and Δχ-tensor values. The grid search of the metal position is centered at a point near the spin-label site and can be controlled with parameters for the step size between neighboring nodes and the inner and outer cutoff radius. This allows adjustment of the grid resolution and limits the search to a volume not too close or far away from its center. The grid search parameters are chosen based on prior knowledge about the approximate position of the spin-label or metal ion binding site and the geometry of the metal binding tag. The described procedure makes it possible to evaluate the PCS score in the absence of an explicit molecular model of the spin-label, with which the experimental PCS data were collected. However, the metal grid search considerably slows down the PCS-Rosetta algorithm, since approximately a few hundred matrix decomposition operations, one per grid node, are carried out at each scoring step. To avoid recurrent calculations of the Δχ-tensor, we implemented an alternative method for determining the metal ion coordinates using an explicit representation of the metal binding tag. To this end, a pre-generated library of side-chain conformations (rotamers) of the spin-label residue is modeled onto the spin-label site and the Δχ-tensor fit is restrained to the coordinates occupied by the metal ion in those conformations. Individual rotamers are treated independently and can overlap with each other but not with neighboring residues. Rotamers with steric clashes are identified prior to the tensor determination by a distance-based criterion (low resolution centroid stage) or an energy cutoff for the interaction with all neighboring backbone and side-chain atoms (high resolution all-atom stage) and are removed from the spin-label ensemble. The final Δχ-tensor and coordinates of *M* are read from the rotamer with the lowest value of $E_{PCS}$ and can be subjected to an additional step of unrestrained optimization to further minimize $E_{PCS}$. It must be pointed out that the described method requires an atomic model of the paramagnetic tag which is not always available. Development of rotamer libraries for a variety of spin-labels will be focus of our future research.

***Rosetta RDC Score:*** The RDC (in Hz) between two protein spins *i* and *j* induced by molecular alignment originating in the interaction between a paramagnetic metal ion *M* and the magnetic field $B_0$ can be calculated as:

$$D_{ij} = -3 \cdot \frac{B_0^2}{15kT} \cdot \frac{\hbar\gamma_i\gamma_j}{8\pi^2 r_{ij}^3} \cdot \left[ (\Delta\chi_{zz} - \overline{\Delta\chi}) \cdot \frac{2z_{ij}^2 - x_{ij}^2 - y_{ij}^2}{2r_{ij}^2} + (\Delta\chi_{xx} - \Delta\chi_{yy}) \cdot \frac{x_{ij}^2 - y_{ij}^2}{r_{ij}^2} + \Delta\chi_{xy} \cdot \frac{2x_{ij}y_{ij}}{r_{ij}^2} + \Delta\chi_{xz} \cdot \frac{2x_{ij}z_{ij}}{r_{ij}^2} + \Delta\chi_{yz} \cdot \frac{2y_{ij}z_{ij}}{r_{ij}^2} \right], \text{with } \overline{\Delta\chi} = Tr(\Delta\chi)/3 \qquad \text{(S3)}$$

where $r_{ij}$ is length of the internuclear connection vector between spins *i* and *j*, $\gamma_i$ and $\gamma_j$ are their respective gyromagnetic ratios, $x_{ij}$, $y_{ij}$ and $z_{ij}$ are the cartesian coordinates of the vector between spins *i* and *j* in the molecular frame *f*, and $\Delta\chi_{xx}$, $\Delta\chi_{yy}$, $\Delta\chi_{zz}$, $\Delta\chi_{xy}$, $\Delta\chi_{xz}$ and $\Delta\chi_{yz}$ are the components of the Δχ-tensor in the frame *f*. It should be noted that, although very similar to the equation of the PCS (eq. (S1)), the RDC does not depend on the distance from the paramagnetic center; the distance between spins *i* and *j* is fixed. Furthermore, RDC eq. (S3) is valid irrespective of the source of partial alignment which can also arise from diamagnetic external alignment media such as liquid crystals or bicelles. If molecular alignment originates only in the anisotropic magnetic susceptibility of the paramagnetic metal ion, the alignment tensor is related to the Δχ-tensor by:

$$\mathbf{D} = \frac{B_0^2}{15k_BT}\boldsymbol{\Delta\chi} \quad \text{(S4)}$$

where $B_0$ is the magnetic field, and $k_B$ and $T$ are Boltzmann's constant and temperature. Thus, the degree of alignment increases with the magnetic field strength and the magnetic susceptibility. In order to make our scoring method generally applicable to RDCs measured under para- and diamagnetic alignment conditions we decided to report the alignment tensor D by its alignment order $D_a$ (in Hz) and rhombicity $R$ instead of $\Delta\chi$.

The alignment tensor is determined each time the RDC score, $E_{RDC}$, is evaluated using singular value decomposition as developed by Losonczi and coworkers (5). $E_{RDC}$ is calculated over all alignment media (or metal ions in case of paramagnetically-induced alignment) $M_k$:

$$E_{RDC} = \sum_k \sqrt{\sum_{\mathrm{ij}} \left(D_{ij}^{calc}(M_k) - D_{ij}^{exp}(\mathrm{M}_k)\right)^2} \quad \text{(S5)}$$

where $D_{ij}^{calc}(M_k)$ and $D_{ij}^{exp}(M_k)$ are the calculated and experimental RDC value of spin pair *ij* induced by medium (metal ion) $M_k$.

***Rosetta PRE Score:*** For metal ions with an isotropic magnetic susceptibility (e.g. $Mn^{2+}$, $Cu^{2+}$, $Gd^{3+}$) and for nitroxide radicals the predominant contribution to the PRE is dipole-dipole (DD) relaxation whereas Curie and DD-Curie cross-correlated relaxation (CCR) play only a minor role (3). These paramagnetic substances have become very popular for PRE measurements and strategies for converting PREs into quantitative distance restraints with reasonable accuracy have been devised (2, 6). On the contrary, lanthanides which have a significant anisotropic magnetic susceptibility have not gained high popularity for PRE measurements because a derivation of reliable distance restraints is more difficult and error prone. PREs are compromised by CCR effects which can be difficult to separate from Curie relaxation, the major component of the PRE for lanthanides. Furthermore, exchange contributions to the nuclear relaxation rate arising from conformational dynamics can be substantially different for the diamagnetic and paramagnetic states owing to the PCS, and thus, the PRE cannot simply be measured as the difference between the relaxation rates in both states (7). For those reasons, we have concentrated on the former group of paramagnetic substances when implementing our PRE scoring method with the aim to simplify the analysis and speed up the calculation. We have adapted the theoretical framework formerly developed by Iwahara and Clore (2), that allowed for accurate interpretation of PRE data and which we briefly revisit here.

For metal ions with isotropic magnetic susceptibility, the PRE arises mostly from dipolar relaxation and its contribution to the longitudinal and transverse relaxation rates of a spin *i* is described by the Solomon-Bloembergen equations:

$$\Gamma_{1,i} = \frac{2}{5}\left(\frac{\mu_0}{4\pi}\right)^2 \gamma_i^2 g^2 \mu_B^2 s(s+1) J(\omega_i) \quad \text{(S6)}$$

$$\Gamma_{2,i} = \frac{1}{15}\left(\frac{\mu_0}{4\pi}\right)^2 \gamma_i^2 g^2 \mu_B^2 s(s+1)\{4J(0) + 3J(\omega_i)\} \quad \text{(S7)}$$

where $\gamma_i$ is the gyromagnetic ratio of spin *i*, *s* is the electron spin quantum number, *g* is the electron g-factor, $\mu_0$ is the permeability of vacuum, $\mu_B$ is the magnetic moment of the free electron, $\omega_i$ is the Lamor frequency of spin *i* and $J(\omega)$ is the spectral density function. The Solomon-Bloembergen theory assumes that the dipole-dipole interaction vector is rigid in the molecular frame; an assumption that breaks down if the space sampled by a paramagnetic group is quite large. To evaluate the effect of the internal dynamics of the spin-label onto the PRE, a "model-free" formalism (8, 9) is incorporated assuming that internal motion and overall tumbling of the protein are uncoupled. In the "model-free" formalism $J(\omega)$ is expressed as:

$$J(\omega) = \langle r^{-6} \rangle \left\{ \frac{S^2 \tau_c}{1 + \omega^2 \tau_c^2} + \frac{(1 - S^2)\tau_t}{1 + \omega^2 \tau_t^2} \right\} \tag{S8}$$

where $\langle r^{-6} \rangle$ is an ensemble-averaged quantity for the proton-metal distance *r* and $S^2$ is the generalized order parameter as defined by Brüschweiler et al. (10). $\tau_c$ represents the correlation time defined as $(\tau_r^{-1} + \tau_e^{-1})$, with $\tau_r$ being the rotational correlation time of the protein and $\tau_e$ the electron relaxation time of the metal ion, and $\tau_t$ represents the total correlation time defined as $(\tau_r^{-1} + \tau_e^{-1} + \tau_i^{-1})$ where $\tau_i$ is the correlation time for internal motion. Thus, two effects are taken into account for evaluating PREs arising from a flexible paramagnetic group: an ensemble effect related to $\langle r^{-6} \rangle$ and a motional effect related to $S^2$ and $\tau_i$. For calculating $\langle r^{-6} \rangle$ and $S^2$, we use a pre-generated library of discrete side-chain conformations of the spin-label residue as already described for the PCS method. In addition, each spin-label rotamer of the ensemble is assigned a weight proportional to its Boltzmann-weighted Rosetta energy with which it contributes to the $\langle r^{-6} \rangle$ and $S^2$ calculation. Finally, correlation times $\tau_c$ and $\tau_t$ are optimized by non-linear least-squares fitting within a predefined range (between $\tau_c^{max}$ and $\tau_c^{min}$).

Using our PRE scoring method, we obtained reasonably good fits to experimental PRE profiles that were reproduceable with PRE Q-factors in the range from 0.36 to 0.57 (compare with **Figure S11**). Furthermore, around 5 – 10 spin-label conformations represent a good compromise between speed and accuracy of the PRE calculation (see **Figure S12**).

In analogy to PCSs and RDCs, the total PRE score is defined as sum of squared deviation between the experimental ($\Gamma_i^{exp}(M_k)$) and calculated ($\Gamma_i^{calc}(M_k)$) PRE values over all metal ions $M_k$:

$$E_{PRE} = \sum_{\mathrm{k}} \sqrt{\sum_{\mathrm{i}} \left(\Gamma_i^{calc}(\mathrm{M_k}) - \Gamma_i^{exp}(\mathrm{M_k})\right)^2} \tag{S9}$$

Finally, for all three score terms described above equations to compute the derivative of the Rosetta energy with respect to the corresponding degrees of freedom (i.e. the coordinates of the vector connecting spin *i* with metal ion *M* in case of PCSs and PREs, or with spin *j* in case of RDCs) were implemented into RosettaNMR. This facilitates using paramagnetic NMR restraints for gradient-based minimization as employed during high-resolution refinement with the Rosetta FastRelax protocol.

#### Method S4: Performance metrics for protein structure prediction

For assessing the performance of paramagnetic NMR data for protein structure modeling with Rosetta, several evaluation metrics were used. The prediction accuracy was quantified by using the Cα-$RMSD_{100}$ which represents a normalized RMSD relative to a protein size of 100 amino acids (11) and was calculated over all residues in ordered regions of the protein as defined in **Table S1** relative to the experimental structure. For proteins determined by NMR the first model of the ensemble was used as reference structure. The $RMSD_{100}$ measure is useful when comparing proteins of varying size such as those used in this benchmark. For symmetric proteins, the Cα-$RMSD_{100}$ was evaluated on the oligomeric model, and for proteins with more than two chains was calculated by testing all possible chain orderings because the order of chains in the Rosetta output models is arbitrary.

The effect of the PCS, RDC and PRE score term on scoring of protein structure models was assessed by the enrichment metric, defined as $(TP/(TP + FP) \times (P + N)/P)$. Therefore, models were sorted according to either their Rosetta score or their combined Rosetta and NMR score. Models that fell within the top 10% by score were counted as “positive” (*P*) and the remaining models were counted as “negative” (*N*). The positives were then sorted by $RMSD_{100}$ and those models that fell within the top 10% by $RMSD_{100}$ were labeled “true positives”

(*TP*); all other models were considered “false positives” (*FP*). Since the $(P+N)/P$ ratio is set to 10, the maximum possible enrichment is also limited to a value of 10.

For a blind structure prediction, it is important to discern whether the final model is reliable or not. As a discrimination measure of a structure prediction we used a combination of two criteria which were evaluated on the 10 lowest-scoring models: backbone convergence and NMR Q-factor. A structure calculation was considered converged if the Cα-RMSD between the lowest-scoring model and the next nine low-scoring models computed for all DSSP-assigned secondary structure (12) regions was less than 4 Å on average. A similar RMSD cutoff is employed in the calculation of other protein structure similarity metrics (e.g. GDT-TS and MaxSub). The agreement with the experimental NMR data was assessed by computing the NMR Q-factor according to Cornilescu et al. (13), $Q=\sqrt{\sum(c_{calc}-c_{exp})^2/\sum(c_{exp})^2}$, where $c_{calc}$ and $c_{exp}$ correspond to the calculated and experimental value of the PCS ($c=\delta_i^{PCS}$), RDC ($c=D_{ij}$) or PRE ($c=\Gamma_i$), respectively. A Q-factor below 0.25 in case of PCSs, and below 0.5 for RDCs and PREs indicates a good structural model consistent with the experimental data.

### Supporting Figures

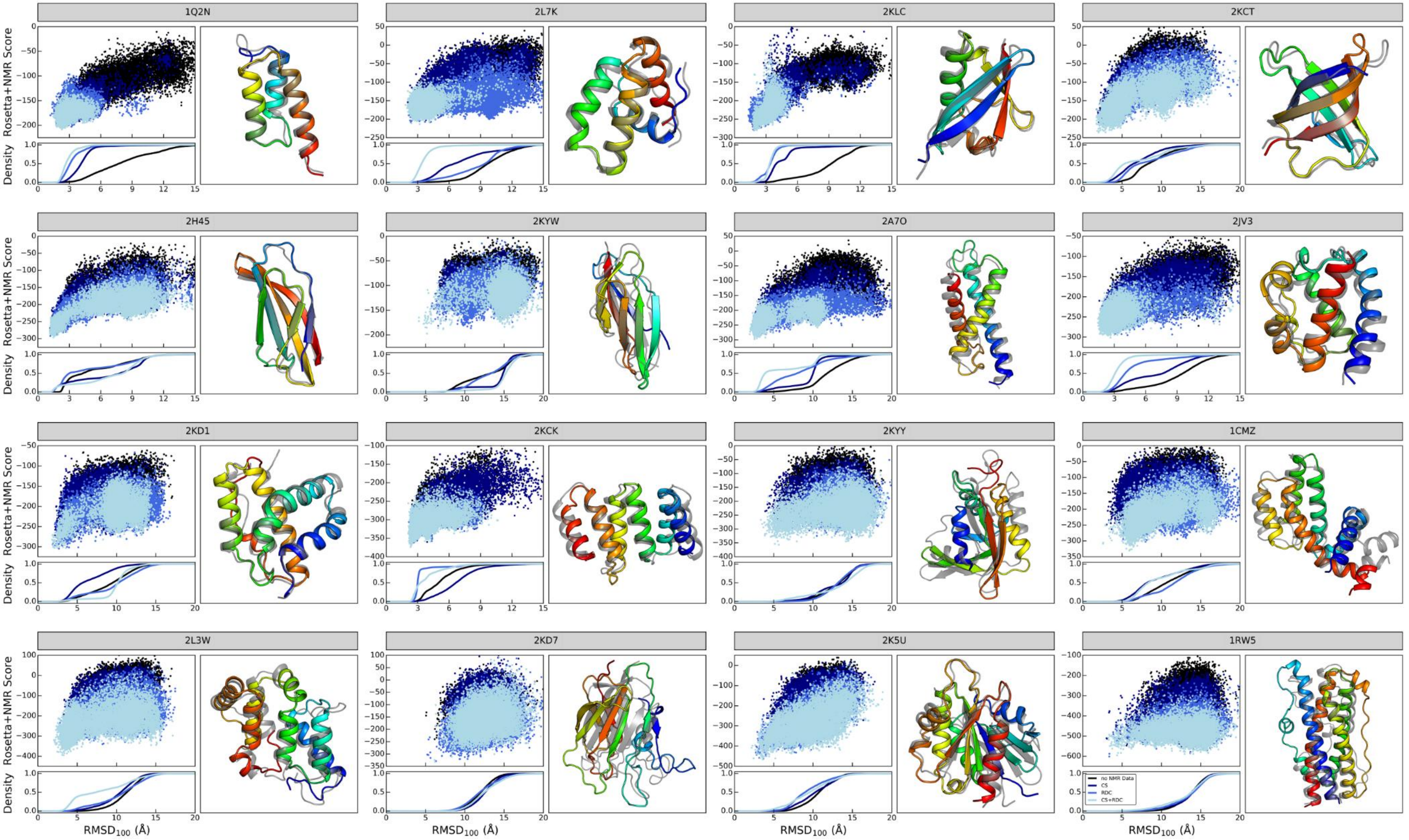


**Figure S1: *De novo* structure prediction of monomeric proteins with chemical shift and RDC data; related to Figure 2 and 3.** For each modeling target, the combined Rosetta and RDC score and the cumulative model density is plotted versus the $RMSD_{100}$ relative to the experimental structure. The models' energy landscape and fraction at low-$RMSD_{100}$ for the unrestrained case (black) is compared with the prediction result when chemical shift data (navy), RDCs (blue), or chemical shift and RDC data (light blue) were used. The lowest scoring model from the calculation with chemical shifts and RDCs is depicted as ribbon diagram and superimposed on the experimental structure in gray. The superposition is optimized for ordered residues as defined in **Table S1** and flexible termini are omitted.

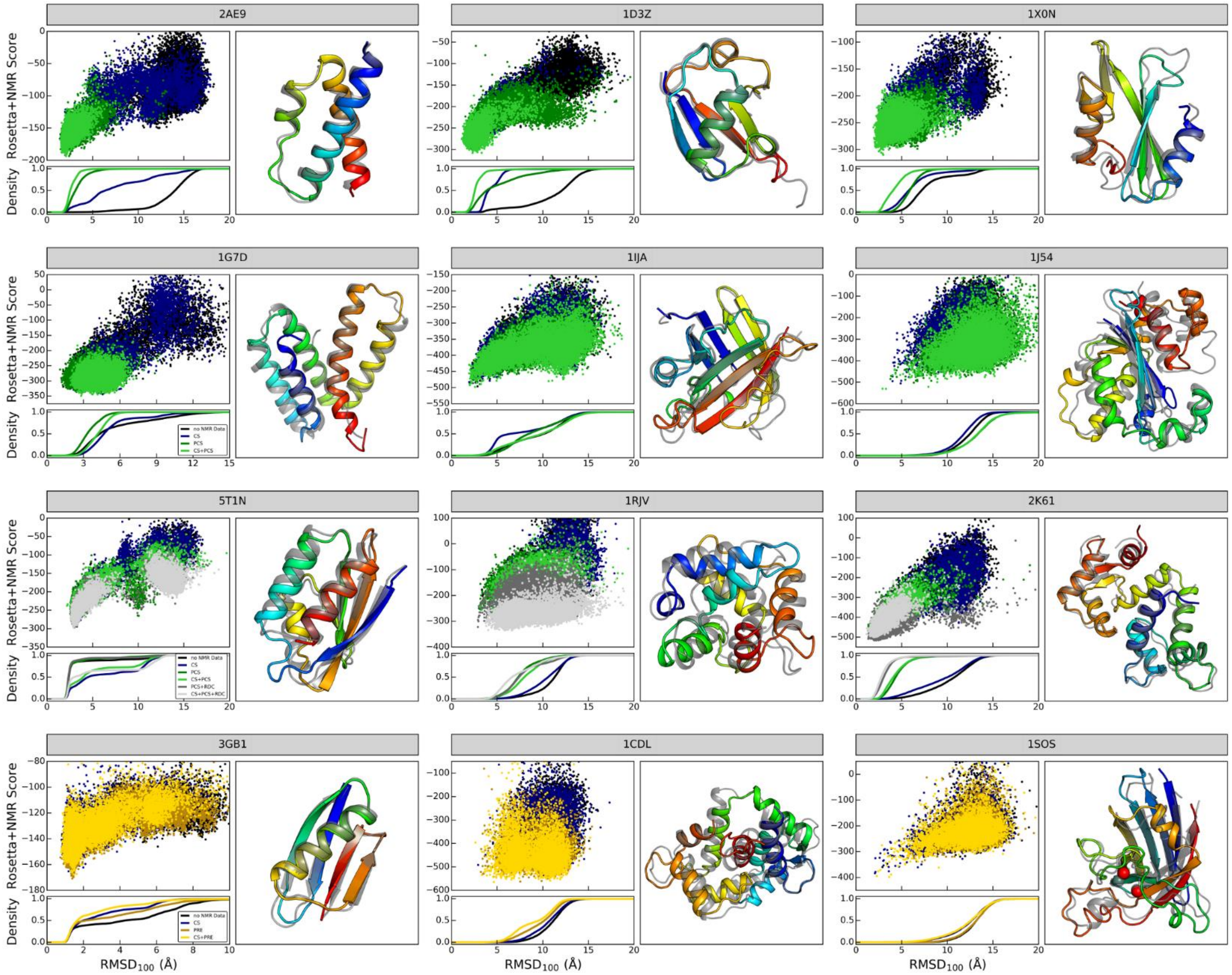


**Figure S2: *De novo* structure prediction of monomeric proteins with chemical shifts, PCS and PRE data; related to Figure 2 and 3.** For each modeling target, the combined Rosetta and paramagnetic NMR score as well as the cumulative model density is plotted versus the $RMSD_{100}$ relative to the experimental structure. For targets 2AE9, 1D3Z, 1X0N, 1G7D, 1IJA and 1J54 the NMR score is calculated from PCSs only, whereas it is composed of the PCS and RDC score term for proteins 5T1N, 1RJV and 2K61, and corresponds to the PRE score in case of 3GB1, 1CDL and 1SOS. For each protein target, the lowest scoring model from the calculation with chemical shifts and the respective paramagnetic NMR data is depicted as ribbon diagram and compared with the experimental structure shown in gray. The superposition is optimized for ordered residues as defined in **Table S1** and flexible termini are omitted for clarity. The colors refer to the following restraint sets: black – no NMR data, navy – only chemical shifts, dark green – PCSs, light green – chemical shifts and PCSs, gray – PCSs and RDCs, light gray – chemical shifts, PCSs and RDCs, olive – PREs, yellow – chemical shifts and PREs.

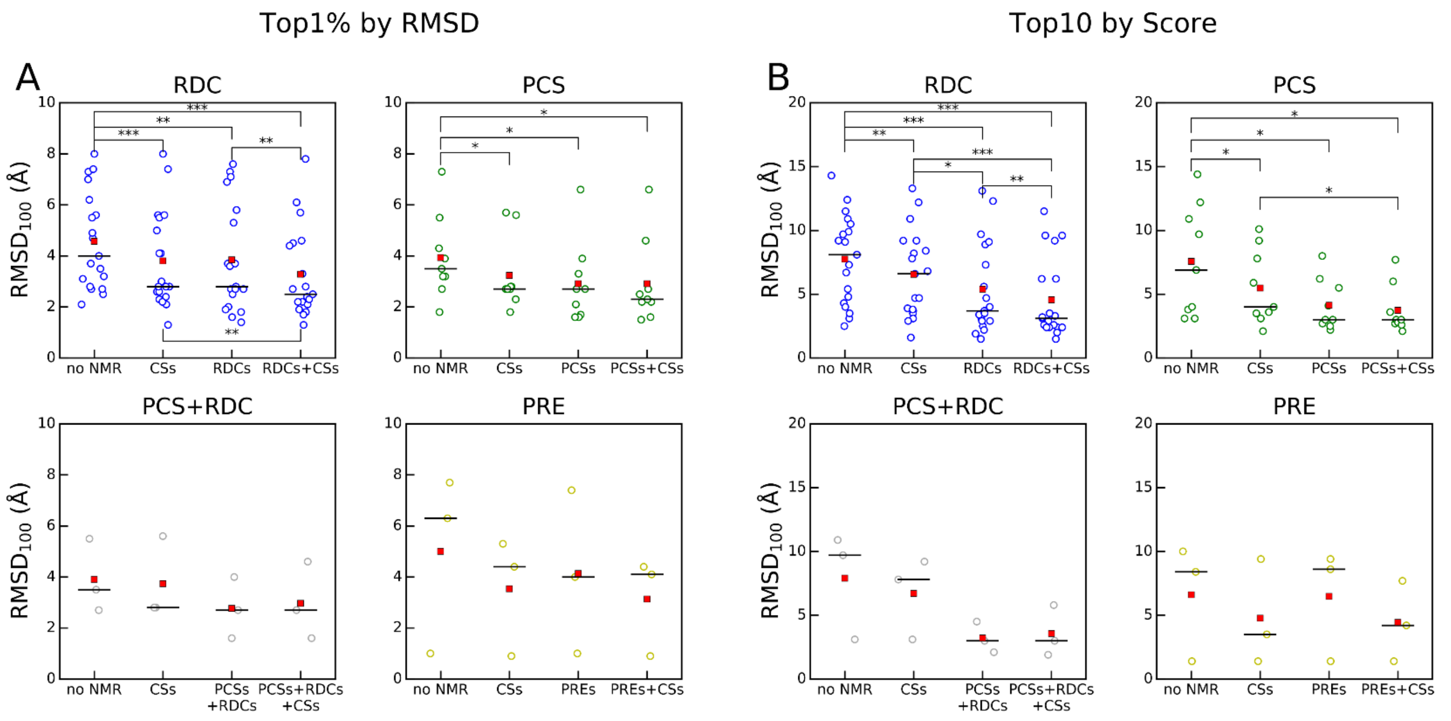


**Figure S3: Comparison of the average $RMSD_{100}$ of the (A) top 1% models by RMSD and (B) the top ten models by score between Rosetta calculations with no NMR data and with different types and combinations of NMR data; related to Figure 3 and Table 1.** For each type of restraint (RDCs: blue, PCSs: green, RDCs + PCSs: gray, PREs: yellow), the $RMSD_{100}$ for every protein target from the benchmark set is plotted, and the distribution median (—) and average (■) are marked. Statistical significance of the $RMSD_{100}$ change was calculated by a two-tailed Wilcoxon signed rank test for restraint types with sufficiently large sample size ($n_{RDC} = 19$, $n_{PCS} = 9$, * $p < 0.05$, ** $p < 0.01$, *** $p < 0.001$).

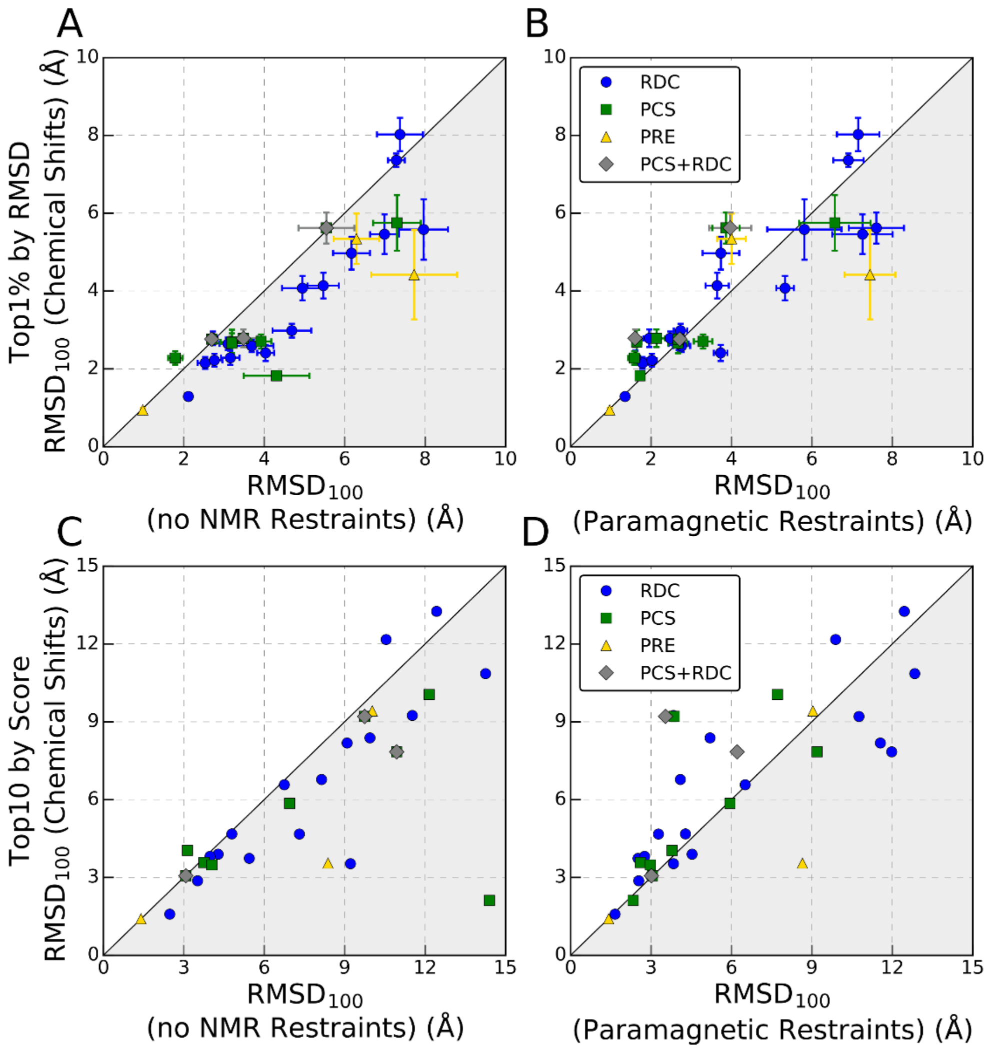


**Figure S4: Accuracy of CS-Rosetta models vs. Rosetta models generated with no NMR data or with paramagnetic NMR restraints; related to Figure 3 and Table 1. (A)** Comparison of the average $RMSD_{100}$ (± S.D.) of the best 1% of models ranked by RMSD between Rosetta models generated with fragments which were selected without or with chemical shifts. The marker symbols indicate the restraint set that each protein belonged to. **(C)** Average $RMSD_{100}$ of the ten lowest-scoring models generated with or without chemical shifts. **(B)** Comparison of the average $RMSD_{100}$ (± S.D.) of the best 1% models identified by RMSD between the Rosetta *de novo* method performed with either chemical shift-selected fragments or with paramagnetic NMR restraints but fragments picked without chemical shifts. **(D)** The average $RMSD_{100}$ of the ten lowest-scoring models derived from the same Rosetta calculations as in **(B)** is shown.

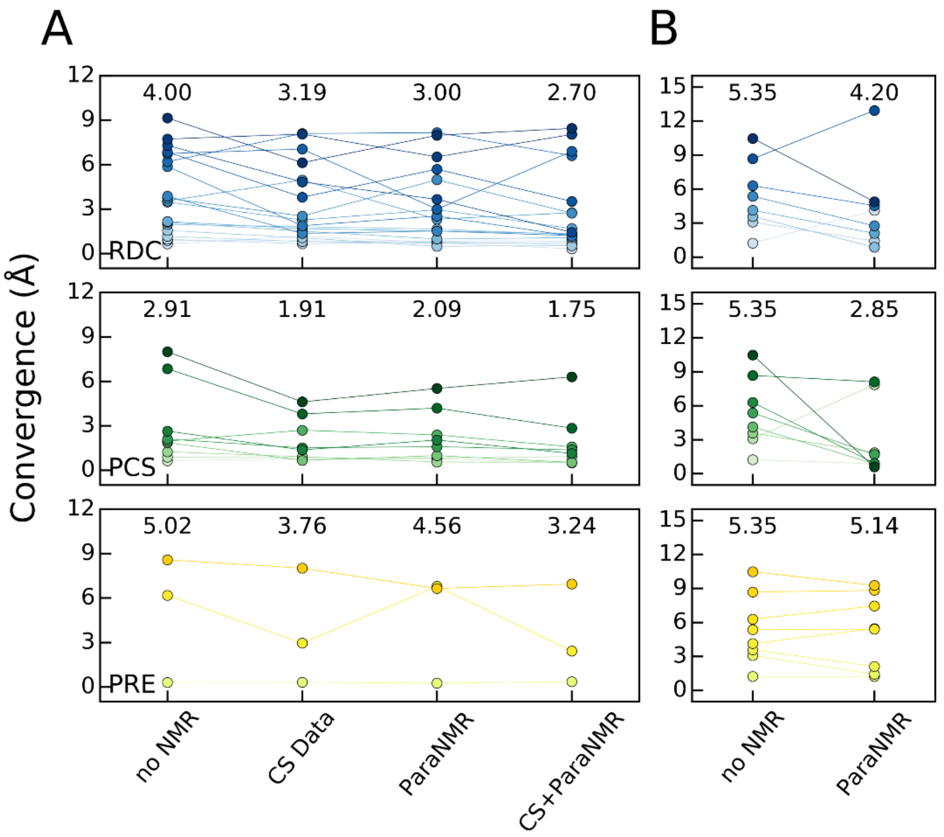


**Figure S5: Effect of NMR restraints on the convergence of Rosetta models; related to Figure 3, 4 and 5. (A)** The RMSD convergence among the ten lowest-scoring models of *de novo* predicted monomeric proteins is plotted as a function of the type and amount of NMR data used in the calculation (no NMR – without NMR data, CS – with chemical shifts, ParaNMR – with RDC, PCS or PRE data, CS+ParaNMR – chemical shifts and RDC, PCS or PRE data). The individual protein targets modeled with RDC, PCS or PRE restraints are represented by using different shades of blue, green or yellow, respectively, and lines between data points are used to guide the eye. The average convergence value across all proteins modeled with a certain restraint set is given on the top of each column. **(B)** The same type of analysis as in **(A)** but for symmetric proteins is shown. The convergence values of all modeled monomeric and symmetric proteins are listed in **Tables S2** and **S4**, respectively, and also shown in **Figure S6**.

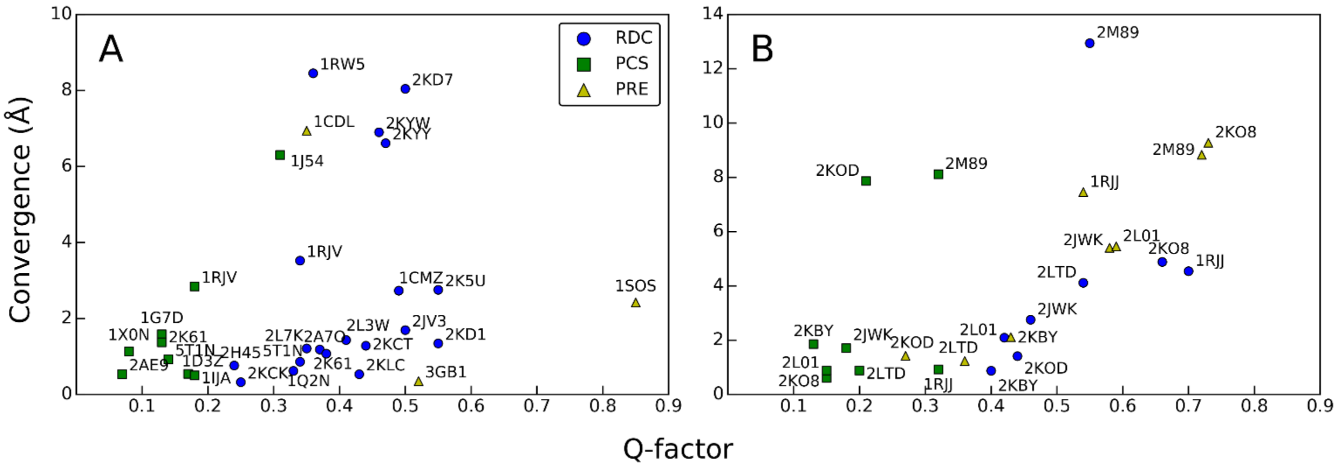


**Figure S6: Identification of successful structure predictions from model convergence and NMR quality factor; related to Figure 3 and 5.** The correlation between model convergence (as defined in Method Details) and the NMR Q-factor (as defined by (13)) is plotted for *de novo* predicted models of **(A)** monomeric and **(B)** symmetric proteins. All proteins are labeled with their respective PDB IDs and their convergence and Q-factor values are listed in **Tables S2** and **S4**. A reliable structure prediction can be identified by high convergence, i.e. a low RMSD deviation between the top-scoring models, and a low NMR Q-factor which reports the agreement between experimental and structure-predicted NMR data. RMSD deviations below 4 Å indicate convergence of the computational protocol, and a Q-factor below 0.25 for PCSs, and below 0.5 for RDCs and PREs indicates good agreement with experimental data. Thus, the convergence and Q-factor can be combined to evaluate success of a structure prediction calculation. Based on this criterion, the predictions for monomeric proteins 1J54, 1RW5, 1CDL, 2KD7, 2KYW, 2KYY and 1SOS are rejected. Proteins 1J54 (ε-subunit of *E. coli* DNA polymerase III) and 1RJV (parvalbumin) are either large or have only a sparse PCS dataset from one lanthanide ion. 2KYW (NSGC target PtR41O), 2KYY (NSGC target NeR70A) and 2KD7 (NSGC target BtR324B) have mostly or exclusively β-sheet content. *De novo* prediction assembled a sheet with incorrect strand order, a topological feature that is insufficiently restrained by RDCs. Prediction of 1SOS (Cu, Zn superoxide dismutase) converged on a single solution but the experimental dataset appeared to contain several erroneous PREs incompatible with native X-ray structure (Q=0.92) which explains the observed high Q-factor of the Rosetta model (Q=0.85). 1RW5 (human prolactin) was the largest protein in the benchmark set (199 residues) but had only 81 RDCs from one alignment medium (<0.5 RDC per residue) likely being insufficient to guide the fragment assembly. Moreover, two 20-25 residue long loops with little secondary structure content further complicated the structure calculation.

In case of symmetric proteins, modeling of 2M89 (with PCSs, RDCs, PREs), 2KO8 (with RDCs, PREs), 1RJJ (with RDCs, PREs), 2KOD (with PCSs), 2JWK and 2L01 (both with PREs) was unsuccessful. 1RJJ (2x 111 residues) and 2M89 (2x 134 residues) were the two largest proteins in the benchmark set. Although low-RMSD models were sampled during symmetric docking, structure prediction failed to converge, as seen by the lack of a docking funnel for RDCs and PREs for 1RJJ and in case of all three NMR data types for 2M89 (see **Figure S8**). The same observation holds true for proteins 2L01 and 2JWK for which no folding funnel developed with PRE data. Another challenge for Fold-and-Dock guided by RDCs and PCSs represented the protein 2KOD, the C-terminal domain of HIV-1 capsid protein. The protein is a C2-symmetric non-interleaved dimer stabilized by an interface of hydrophobic residues from helix 9 of both subunits. In addition to the native topology, Fold-and-Dock sampled an interleaved dimer in which helix 8 is swapped between the two subunits and occupies the same position and orientation as in the native dimer. Due to the symmetric shape of the alignment and Δχ-tensors, the interleaved dimer fits the RDC and PCS data equally well and the simulation is misguided to the non-native dimer topology. In contrast, PREs offered a better discrimination and clearly favored the native dimer. We reason that this is most likely because of small differences in the inter-subunit distance between the two dimer topologies and the high sensitivity of the PRE to small distance variations.

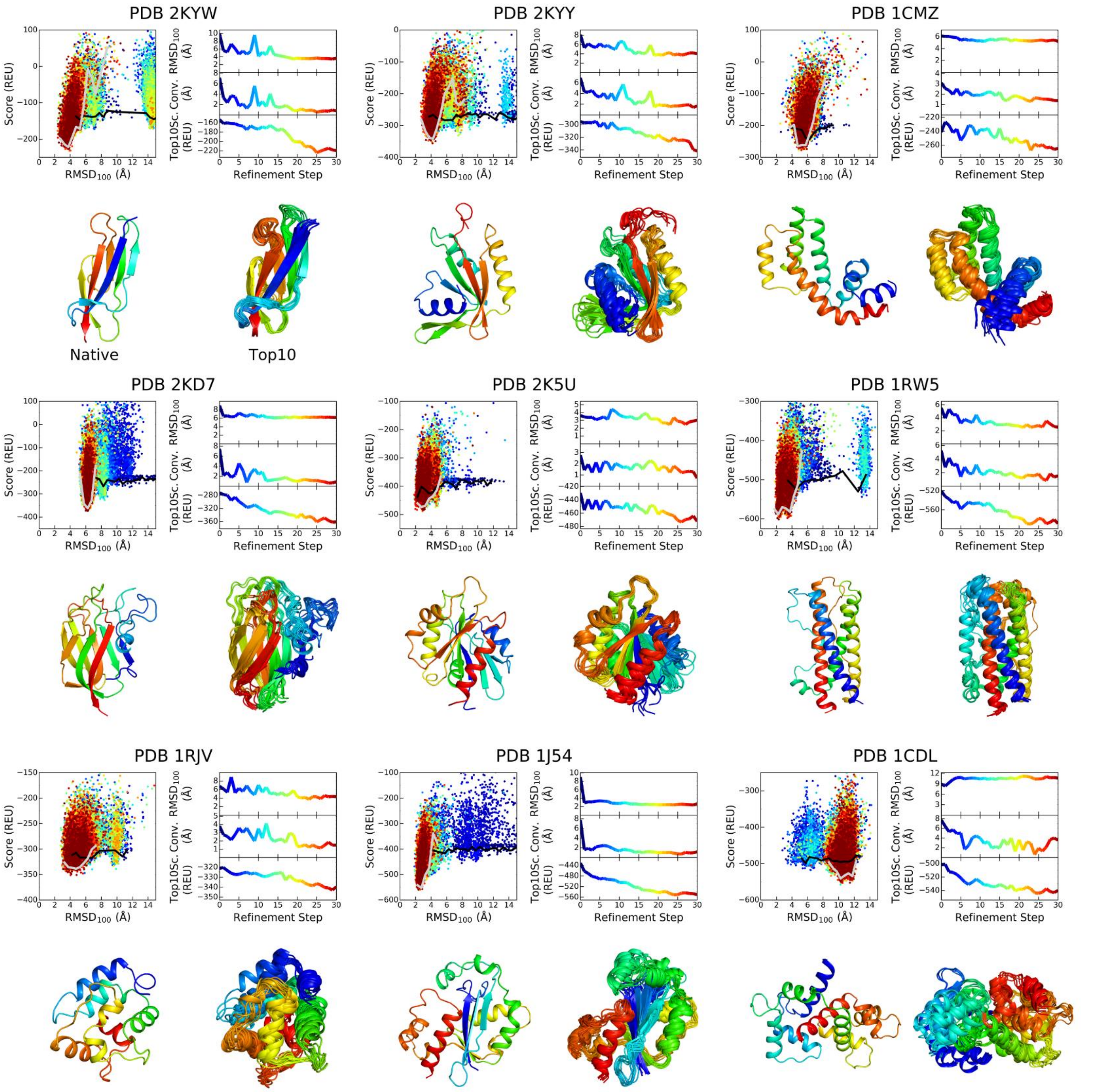


**Figure S7: Protein model refinement with Iterative Hybridize using RDC, PCS and PRE data; related to Table 1.** For each of the nine protein targets (labeled by their corresponding PDB IDs), 30 rounds of Iterative Hybridize were carried out and 720 models were created in each round. The panels display the following results: The upper left panel plots the combined Rosetta and NMR score versus the model's $RMSD_{100}$ relative to the experimental structure. The NMR score was calculated by scoring models with their respective RDC, PCS or PRE restraints. The color coding (blue to red) corresponds to the number of refinement steps. The black and gray line represent the lowest energy rim (i.e. the median of the 5 lowest-scoring models by RMSD bin) of the score-vs-RMSD plot of the model pool at step 0 and 30, respectively. Note that the pool at step 0 is comprised of models after the first Iterative Hybridize selection step. The upper right panel displays the average $RMSD_{100}$ (top row), convergence (middle row) and average score (bottom row) of the ten lowest scoring models after each refinement step. Convergence was calculated as average RMSD over all DSSP-assigned secondary structure regions in the protein. The values of successive refinement steps are colored from blue to red. The lower panel compares the native structure model (left) depicted as ribbon diagram and colored in rainbow with the ensemble of ten lowest-scoring models (Top10) obtained after the last refinement step. For clarity, flexible termini are omitted as described in the footnote to **Table S1**.

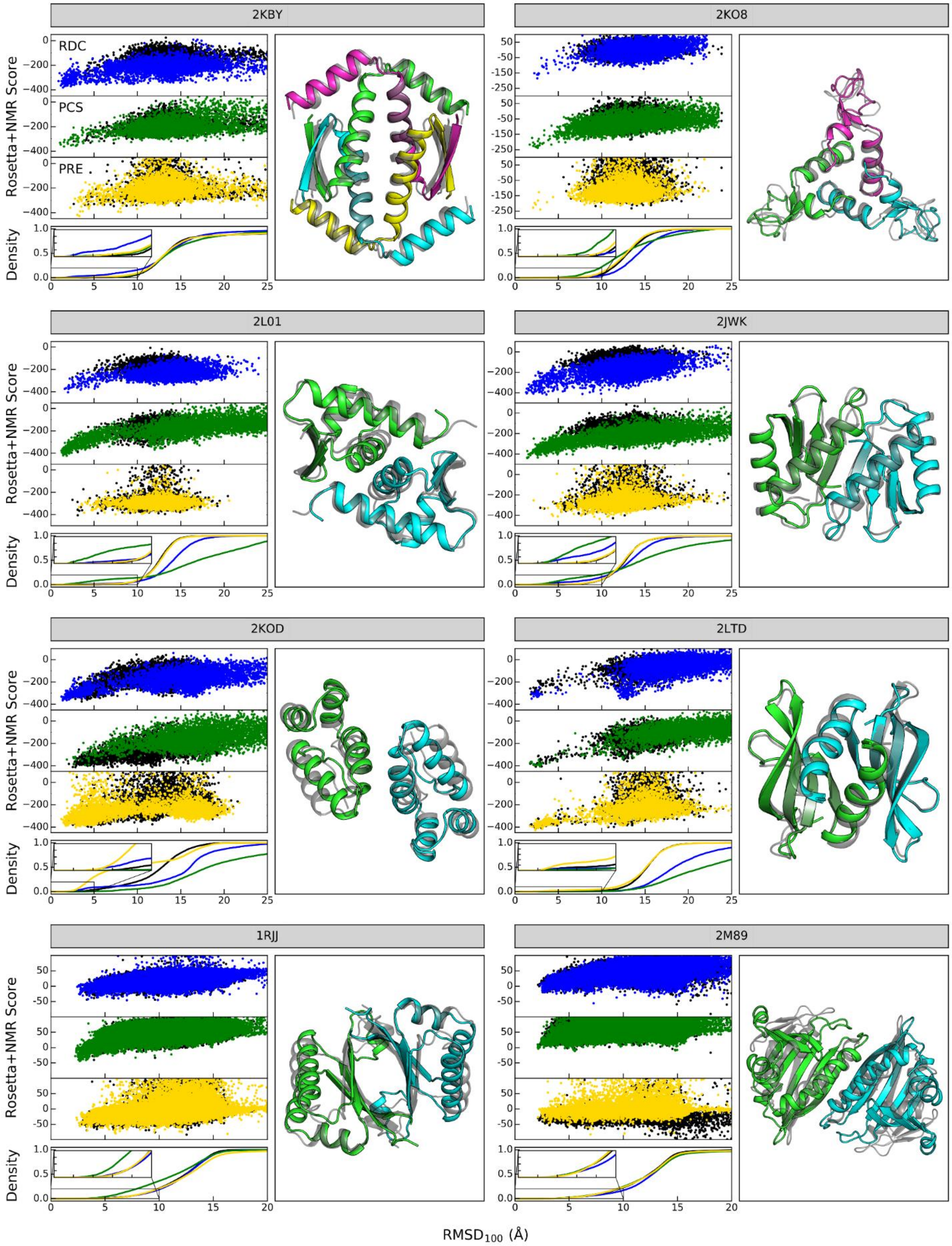


**Figure S8: Results of *de novo* structure prediction of symmetric proteins with experimental RDC and simulated PCS and PRE data; related to Figure 6 and Table 2.** For each protein target, the combined Rosetta and NMR score as well as the model density is plotted versus the $RMSD_{100}$ relative to the experimental structure. The NMR score was calculated from the respective NMR restraint type that had been used for structure prediction, i.e. RDCs (blue), PCSs (green) or PREs (yellow), respectively, and models that had been generated without restraints were rescored accordingly. In each case, a representative low-scoring model from one of the NMR calculations is displayed as ribbon diagram and compared with the experimental structure colored gray. The superposition and $RMSD_{100}$ calculation were done for residues in ordered regions as described in the footnote to **Table S1**. For proteins corresponding to PDB IDs 1RJJ and 2M89, only the results of the symmetric docking experiment are shown.

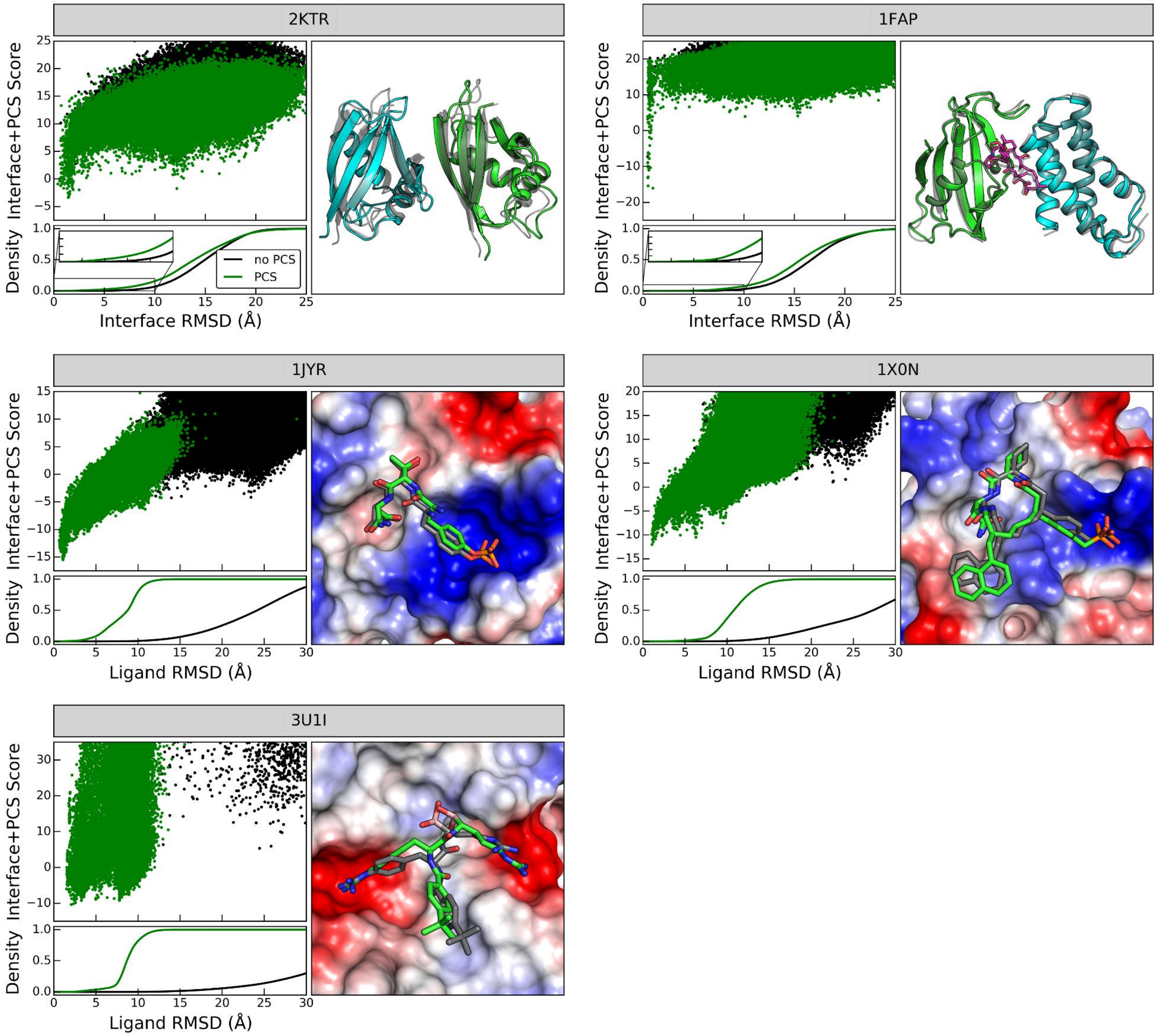


**Figure S9: Results of the PCS-assisted protein-protein and protein-ligand docking; related to Figure 7 and Table 3.** The first row displays the results of protein-protein docking with PCS data: docking of the p62 PB1 dimer (PDB ID 2KTR) and docking of the FKBP12-rapamycin-binding domain (FRB) to human FK506-binding protein complexed with rapamycin (PDB ID 1FAP). The second and third rows show the results of PCS-assisted ligand-docking: docking of a pYTN peptide and a high affinity inhibitor to the SH2 domain of human growth factor receptor binding protein (Grb2) (PDB IDs 1JYR and 1X0N), and docking of a high affinity inhibitor to dengue virus serotype 3 (DENpro) (PDB IDs 3U1I). For each test case, the combined Rosetta interface and PCS score as well as the fraction of models is plotted versus the RMSD relative to the experimental structure. The lowest-scoring model produced by docking with PCSs is displayed as a cartoon representation and compared with the experimental structure of the protein-protein or protein-ligand complex, respectively. In case of the DENV-3-inhibitor complex, the Rosetta model was compared with and the ligand RMSD was calculated to a previously published structural model (14) generated with AutoDock-Vina (15) using PCS and intermolecular NOE data.

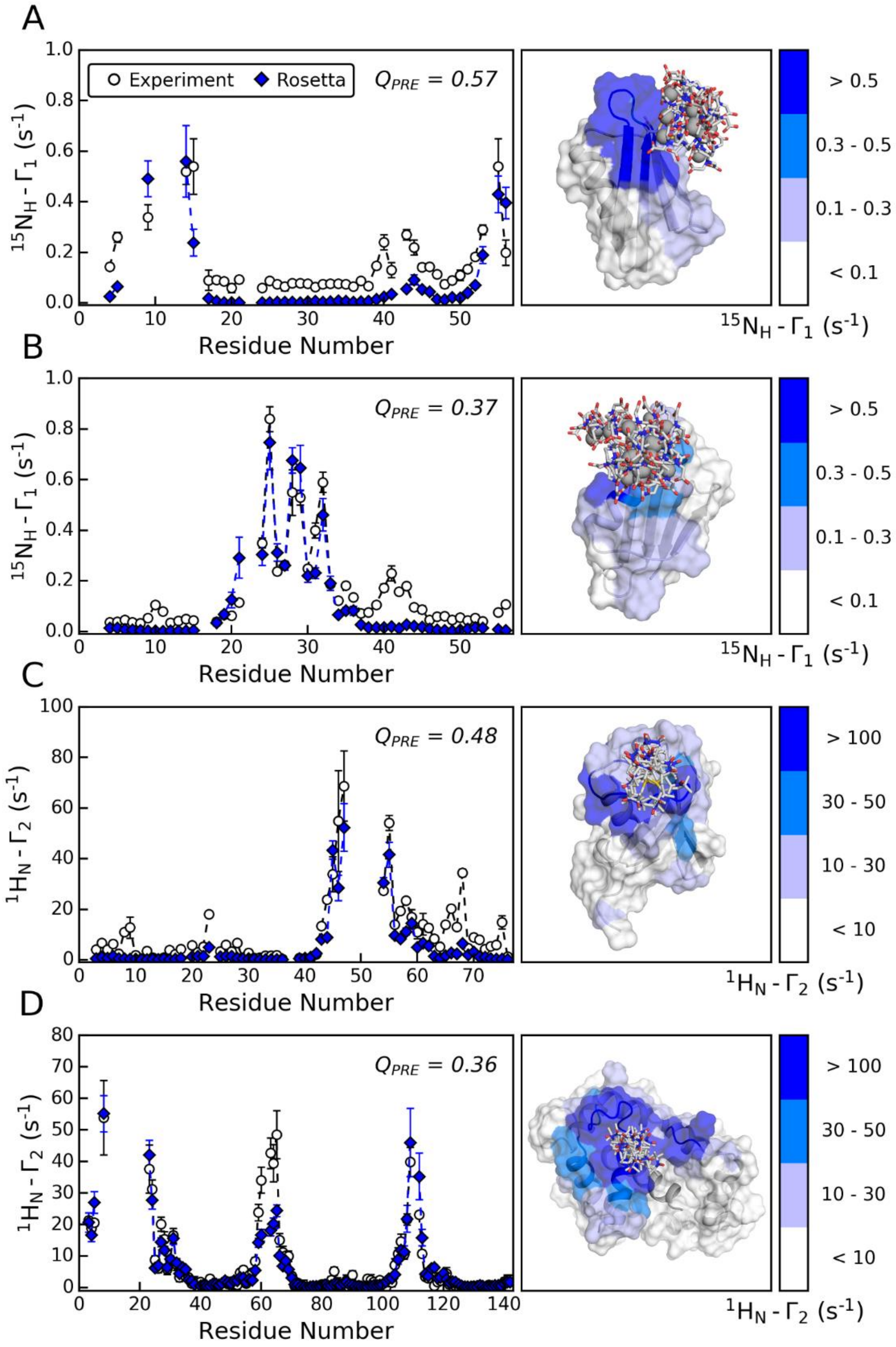


**Figure S10: Comparison of experimental and Rosetta-simulated PRE data for four proteins.** Experimental and calculated PRE values for $Cu^{2+}$-MTS-EDTA-labeled variants N8C **(A)** and K28C **(B)** of the B1 domain of protein G (16), MTSL-labeled ubiquitin variant K48C (17) **(C)**, and MTSL-labeled calmodulin variant S17C (18) **(D)**. PRE values were calculated with Rosetta's PRE scoring method using the relaxed native structure with PDB ID 3GB1 for protein G, 1UBQ for ubiquitin and 1CDL for MLCK-peptide-bound calmodulin. Error bars represent one standard deviation computed by a Monte-Carlo error estimation protocol in which 30% of the spin-label side-chain conformations were randomly deleted. A representation of the spin-label ensemble drawn in gray sticks is given on the righthand side of the figure, and the experimental PRE values are mapped onto the protein surface.

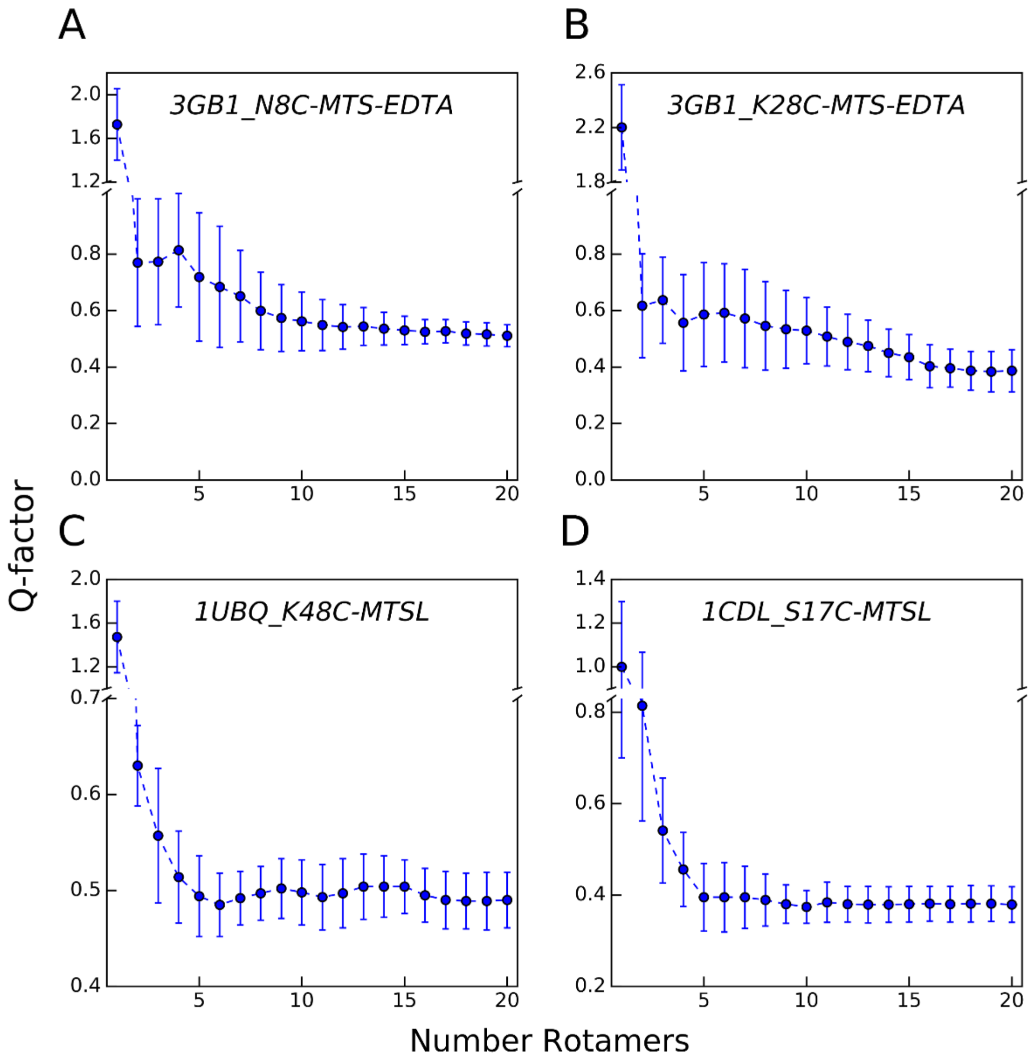


**Figure S11: Influence of the number of spin-label rotamers on the PRE Q-factor.** The Q-factor reports the agreement of the experimental PRE data with those predicted from the native protein structure using a spin-label model over a variable number of rotamers. The experimental datasets were the same as in **Figure S10**: $^{15}N_H$-$\Gamma_1$ data collected on variants N8C **(A)** and K28C **(B)** of the B1 domain of protein G conjugated with $Cu^{2+}$-MTS-EDTA (16), and $^{1}H_N$-$\Gamma_2$ data derived for MTSL-labeled ubiquitin variant K48C (17) **(C)** and MTSL-labeled calmodulin variant S17C (18) **(D)**. Error bars represent one standard deviation computed by a Monte-Carlo error estimation protocol in which one thousand random samples made up of only 70% of the spin-label rotamers were successively drawn. The samples were clustered by average linkage hierarchical clustering with a cluster number corresponding to the desired number of rotamers, and the PRE was simulated from those rotamers corresponding to the centroids of each cluster.

### Supporting Tables

**Table S1:** Type and source of NMR data used for structure prediction of monomeric and symmetric proteins; related to **Figure 2**, **3**, **4** and **5**, and **Table 1** and **2**.

| Protein Name | PDB ID* | BMRB ID | No. Chains | No. Residues† | No. RDC | Atom Types RDC | No. PCS | Atom Types PCS | No. PRE | Atom Types PRE | No. RDC Alignment Media | Spin-Label Site‡ | Metal Ions Used | Ref. |
|---|---|---|---|---|---|---|---|---|---|---|---|---|---|---|
| **Monomeric** | | | | | | | | | | | | | | |
| Z domain of staphylococcal protein A | 1Q2N | 5656 | 1 | 58 (1-58) | 126 | N-$H^N$, Cα-C', Cα-Hα | | | | | 1 | | | (19) |
| Protein CD1104.2 from C. difficile | 2L7K | 17359 | 1 | 76 (1-68) | 79 | N-$H^N$ | | | | | 2 | | | ** |
| Domain of adhesion exoprotein from P. pentosaceus | 2KYW | 16988 | 1 | 87 (1-79) | 132 | N-$H^N$ | | | | | 2 | | | ** |
| Ob fold of heme chaperone CCME from D. vulgaris | 2KCT | 16096 | 1 | 94 (51-127) | 49 | N-$H^N$ | | | | | 1 | | | ** |
| Second type III domain of human fibronectin | 2H45 | 7127 | 1 | 95 (20-95) | 50 | N-$H^N$ | | | | | 1 | | | (20) |
| Human ubiquitin-like domain of ubiquitin 1 | 2KLC | 16390 | 1 | 101 (23-97) | 75 | N-$H^N$, Cα-C' | | | | | 1 | | | ** |
| Ets-1 PNT domain | 2JV3 | 4205 | 1 | 110 (41-136) | 91 | N-$H^N$ | | | | | 1 | | | (21) |
| hSet2/HYPB SRI domain | 2A7O | 6834 | 1 | 112 (10-101) | 120 | N-$H^N$ | | | | | 1 | | | (22) |
| NESG target MrR121A | 2KCK | 16083 | 1 | 112 (4-104) | 144 | N-$H^N$ | | | | | 2 | | | ** |
| Integrase-like domain from B. cereus ordered locus BC_1272 | 2KD1 | 16102 | 1 | 118 (4-110) | 105 | N-$H^N$ | | | | | 1 | | | ** |
| G-alpha interacting protein (GAIP) | 1CMZ | 4407 | 1 | 128 (79-206) | 291 | N-$H^N$, Cα-C', Cα-Hα, N-C' | | | | | 2 | | | (23) |
| PBS linker domain of phycobilisome linker polypeptide from S. elongatus | 2L3W | 17207 | 1 | 143 (3-132) | 176 | N-$H^N$ | | | | | 2 | | | ** |
| N-ter domain of DNA helicase RecG-related protein from N. europaea | 2KYY | 16991 | 1 | 152 (2-122) | 81 | N-$H^N$ | | | | | 1 | | | ** |
| F5/8 type C-ter. domain of a putative chitobiase from B. thetaiotaomicron | 2KD7 | 16107 | 1 | 159 (4-151) | 137 | N-$H^N$ | | | | | 2 | | | ** |
| Myristoylated yeast ARF1 protein | 2K5U | 15809 | 1 | 181 (15-180) | 322 | N-$H^N$, N-C', C'-$H^N$ | | | | | 3 | | | (24) |
| Human prolactin | 1RW5 | 6643 | 1 | 199 (13-196) | 81 | N-$H^N$ | | | | | 1 | | | (25) |
| Θ-subunit E. coli DNA polymerase III | 2AE9 | 6571 | 1 | 76 (10-64) | | | 86 | $H^N$ | | | | D14 | $Dy^{3+}$, $Er^{3+}$ | (26, 27) |
| Ubiquitin | 1D3Z | 6457 | 1 | 76 (1-76) | | | 331 | $H^N$ | | | | E18<br>T66 | $Tb^{3+}$, $Tm^{3+}$<br>$Tb^{3+}$, $Tm^{3+}$ | (13, 28) |
| SH2 domain of growth factor receptor binding protein 2 | 1X0N | 11055 | 1 | 104 (65-150) | | | 227 | $H^N$ | | | | M73 | $Dy^{3+}$, $Tb^{3+}$, $Er^{3+}$, $Tm^{3+}$ | (29, 30) |
| C-terminal domain of rat ERp29 | 1G7D/ *2QC7*¶ | 4920 | 1 | 106 (155-228,230-250) | | | 276 | $H^N$ | | | | C157<br><br>S200<br>A218<br>Q241 | $Tm^{3+}$, $Tb^{3+}$, $Yb^{3+}$, $Dy^{3+}$<br>$Tb^{3+}$, $Tm^{3+}$<br>$Tb^{3+}$, $Tm^{3+}$<br>$Tb^{3+}$, $Tm^{3+}$ | (31, 32) |
| Sortase A from S. aureus | 1IJA | 4879 | 1 | 148 (11-148) | | | 41 | $H^N$ | | | | Q55 | $Tm^{3+}$ | (28, 33) |

| | | | | | | | | | | | | | | |
|---|---|---|---|---|---|---|---|---|---|---|---|---|---|---|
| N-terminal exonuclease domain of ε-subunit of E. coli DNA polymerase III | *1J54* | 6184 | 1 | 186 (7-180) | | | 480 | $H^N$, N | | | | D12 | $Dy^{3+}$, $Er^{3+}$, $Tb^{3+}$ | (34, 35) |
| NPr | 5T1N | 30158 | 1 | 85 (1-85) | 57 | N-$H^N$ | 102 | $H^N$, N | | | 1 | E45 | $Yb^{3+}$ | (36) |
| Parvalbumin | 1RJV | 6049 | 1 | 110 (2-109) | 46 | N-$H^N$ | 104 | $H^N$, N | | | 1 | D93 | $Dy^{3+}$ | (37) |
| Calmodulin in DAPK-peptide bound state | 2K61 | 15852 | 1 | 146 (3-148) | 328 | N-$H^N$ | 403 | $H^N$ | | | 4 | D60 | $Tm^{3+}$, $Tb^{3+}$, $Yb^{3+}$, $Dy^{3+}$ | (38) |
| B1 domain of streptococcal protein G | 3GB1 | 7280 | 1 | 56 (1-56) | | | | | 91 | N | | N8<br>N28 | $Cu^{2+}$<br>$Cu^{2+}$ | (16, 39) |
| Calmodulin bound to peptide analog of CaM-binding region of chicken MLCK | *1CDL*[§] | 1634 | 2 | 142 (5-146) + 19 (147-165) | | | | | 120 | $H^N$ | | S17 | Nitroxide[#] | (18, 40) |
| Cu,Zn Superoxide Dismutase | *1SOS* | 4202 | 1 | 153 (1-153) | | | | | 179 | N, C’ | | H48 | $Cu^{2+}$ | (41, 42) |
| **Symmetric** | | | | | | | | | | | | | | |
| Tetramerization domain of human p73 | 2KBY | 25958 | 4 | 4 x 50 (3-46) | 154 (Exp.) | N-$H^N$, Cα-Hα, Cα-C’, N-C’, C’-$H^N$ | 115 (Sim.) | $H^N$ | 69 (Sim.) | $H^N$ | 1 | M19<br>Q41 | $Tb^{3+}$, $Tm^{3+}$, Nitroxide[#] | (43) |
| Tryptophan RNA-binding attenuator protein inhibitory protein | 2KO8 | 16492 | 3 | 3 x 53 (2-53) | 43 (Exp.) | N-$H^N$ | 131 (Sim.) | $H^N$ | 125 (Sim.) | $H^N$ | 1 | E22<br>D45 | $Tb^{3+}$, $Tm^{3+}$, Nitroxide[#] | (44) |
| Periplasmic domain of TolR from H. influenzae | 2JWK | 15459 | 2 | 2 x 74 (20-92) | 263 (Exp.) | N-$N^H$, Cα-Hα, Cα-C’, N-C’ | 233 (Sim.) | $H^N$ | 168 (Sim.) | $H^N$ | 1 | E43<br>Q52 | $Tb^{3+}$, $Tm^{3+}$, Nitroxide[#] | (45) |
| Protein BVU3908 from B. vulgatus | 2L01 | - | 2 | 2 x 77 (1-69) | 97 (Exp.) | N-$H^N$ | 223 (Sim.) | $H^N$ | 174 (Sim.) | $H^N$ | 2 | E17<br>K32 | $Tb^{3+}$, $Tm^{3+}$, Nitroxide[#] | ** |
| Apo YdbC from L. lactis | 2LTD | 18469 | 2 | 2 x 80 (1-74) | 102 (Exp.) | N-$H^N$ | 224 (Sim.) | $H^N$ | 188 (Sim.) | $H^N$ | 2 | E12<br>K67 | $Tb^{3+}$, $Tm^{3+}$, Nitroxide[#] | ** |
| C-terminal domain of HIV-1 CA | 2KOD | 16555 | 2 | 2 x 88 (148-220) | 134 (Exp.) | N-$H^N$, Cα-Hα | 232 (Sim.) | $H^N$ | 173 (Sim.) | $H^N$ | 2 | K170<br>A204 | $Tb^{3+}$, $Tm^{3+}$, Nitroxide[#] | (46) |
| Protein At5g22580 from A. thaliana | 1RJJ | 6011 | 2 | 2 x 111 (1-111) | 105 (Exp.) | N-$H^N$ | 536 (Sim.) | $H^N$ | 359 (Sim.) | $H^N$ | 1 | K25<br>V71<br>D91 | $Tb^{3+}$, $Tm^{3+}$, Nitroxide[#] | (47) |
| Protein Aha1 from C. psychrerythraea | 2M89 | 19235 | 2 | 2 x 134 (1-134) | 92 (Exp.) | N-$H^N$ | 657 (Sim.) | $H^N$ | 437 (Sim.) | $H^N$ | 2 | Q19<br>C68<br>S124 | $Tb^{3+}$, $Tm^{3+}$, Nitroxide[#] | (48) |

* Proteins with reference X-ray structures are in italic; for proteins determined by NMR the first model of the NMR ensemble was used as reference structure.
[†] Total number of residues. The first and last residue numbers (in original PDB numbering) of the ordered region without the flexible termini are listed in parentheses; these and all intervening residues were used to superimpose models and calculate the $RMSD_{100}$ relative to the experimental structure.
[‡] Grid search of the metal ion position for PCS scoring or simulation of the spin-label ensemble for PRE scoring was centered around the reported experimental spin-label position (reported in original PDB numbering). For proteins 2AE9, 1RJV, 2K61, 1J54 and 1SOS with natural metal binding sites a conserved coordinating residue was used to define the center of the grid search.
[§] Protein 1CDL was a complex between calmodulin and a peptide from myosin light chain kinase. The model’s $RMSD_{100}$ was calculated over all residues in both polypeptide chains.
[¶] The model for protein 1G7D was compared to the X-ray structure of the homologous human protein (PDB 2QC7) because previous studies (31) uncovered flaws in the structure determined by NMR. The $RMSD_{100}$ was calculated over residues 155-228 and 230-250, because residue K229 is inserted into the human protein.
[#] The MTSL nitroxide spin-label was used for modeling.
** Proteins represent targets of the northeast structural genomics consortium.

**Table S2:** NMR quality factor and convergence among the ten lowest-scoring models of *de novo* predicted monomeric proteins; related to **Figure 2**, **3** and **4** and **Table 1**.

| | no paramagnetic NMR restraints* | | | | with paramagnetic NMR restraints† | | | | |
|---|---|---|---|---|---|---|---|---|---|
| | no chemical shifts | | with chemical shifts | | no chemical shifts | | with chemical shifts | | Native‡ |
| **PDB** | **Q-factor** | **Convergence§** | **Q-factor** | **Convergence** | **Q-factor** | **Convergence** | **Q-factor** | **Convergence** | **Q-factor** |
| **RDC** | | | | | | | | | |
| 1Q2N | 0.43 | 0.96 | 0.38 | 0.66 | 0.35 | 0.77 | 0.33 | 0.62 | 0.33 |
| 2L7K | 0.51 | 3.87 | 0.40 | 1.36 | 0.37 | 1.52 | 0.35 | 1.21 | 0.27 |
| 5T1N | 0.38 | 0.88 | 0.39 | 0.85 | 0.34 | 0.84 | 0.34 | 0.86 | 0.39 |
| 2KYW | 0.64 | 6.75 | 0.64 | 7.05 | 0.57 | 2.99 | 0.46 | 6.90 | 0.42 |
| 2KCT | 0.64 | 1.96 | 0.55 | 1.81 | 0.56 | 1.65 | 0.44 | 1.28 | 0.39 |
| 2H45 | 0.37 | 1.16 | 0.27 | 0.81 | 0.22 | 0.70 | 0.24 | 0.76 | 0.20 |
| 2KLC | 0.53 | 1.56 | 0.50 | 1.06 | 0.43 | 0.49 | 0.43 | 0.53 | 0.44 |
| 2JV3 | 0.58 | 3.48 | 0.55 | 2.26 | 0.48 | 2.96 | 0.50 | 1.69 | 0.53 |
| 1RJV | 0.64 | 6.85 | 0.55 | 3.80 | 0.49 | 5.69 | 0.34 | 3.52 | 0.44 |
| 2A7O | 0.64 | 5.87 | 0.40 | 1.88 | 0.43 | 2.53 | 0.37 | 1.18 | 0.28 |
| 2KCK | 0.31 | 0.65 | 0.34 | 1.18 | 0.28 | 0.73 | 0.25 | 0.32 | 0.50 |
| 2KD1 | 0.66 | 2.16 | 0.62 | 1.68 | 0.57 | 1.52 | 0.55 | 1.34 | 0.52 |
| 1CMZ | 0.68 | 3.73 | 0.64 | 2.53 | 0.55 | 4.98 | 0.48 | 2.73 | 0.36 |
| 2L3W | 0.70 | 7.30 | 0.65 | 4.83 | 0.46 | 3.66 | 0.40 | 1.43 | 0.28 |
| 2K61 | 0.56 | 2.10 | 0.48 | 1.51 | 0.33 | 0.98 | 0.38 | 1.07 | 0.45 |
| 2KYY | 0.75 | 6.20 | 0.68 | 8.10 | 0.46 | 8.16 | 0.47 | 6.61 | 0.49 |
| 2KD7 | 0.70 | 7.73 | 0.72 | 8.06 | 0.50 | 6.53 | 0.50 | 8.04 | 0.26 |
| 2K5U | 0.74 | 3.55 | 0.67 | 4.96 | 0.57 | 2.32 | 0.55 | 2.75 | 0.44 |
| 1RW5 | 0.52 | 9.15 | 0.44 | 6.13 | 0.39 | 7.98 | 0.36 | 8.45 | 0.34 |
| **PCS** | | | | | | | | | |
| 2AE9 | 0.12 | 1.86 | 0.07 | 0.66 | 0.08 | 0.98 | 0.07 | 0.53 | 0.09 |
| 1D3Z | 0.20 | 1.24 | 0.19 | 0.91 | 0.18 | 0.55 | 0.17 | 0.54 | 0.17 |
| 5T1N | 0.14 | 0.88 | 0.14 | 0.85 | 0.13 | 0.82 | 0.14 | 0.92 | 0.13 |
| 1X0N | 0.14 | 2.64 | 0.10 | 1.37 | 0.11 | 2.04 | 0.08 | 1.13 | 0.04 |
| 1G7D | 0.16 | 1.96 | 0.14 | 2.70 | 0.14 | 2.38 | 0.13 | 1.58 | 0.14 |
| 1RJV | 0.22 | 6.85 | 0.24 | 3.80 | 0.17 | 4.19 | 0.18 | 2.84 | 0.15 |
| 2K61 | 0.17 | 2.10 | 0.14 | 1.51 | 0.12 | 1.59 | 0.13 | 1.37 | 0.11 |
| 1IJA | 0.21 | 0.64 | 0.21 | 0.77 | 0.20 | 0.77 | 0.18 | 0.50 | 0.19 |
| 1J54 | 0.44 | 8.00 | 0.36 | 4.61 | 0.31 | 5.52 | 0.31 | 6.30 | 0.11 |
| **RDC + PCS** | | | | | | | | | |
| 5T1N | 0.38 / 0.14 | 0.88 | 0.39 / 0.14 | 0.85 | 0.35 / 0.13 | 0.98 | 0.37 / 0.14 | 0.80 | 0.39 / 0.13 |
| 1RJV | 0.64 / 0.22 | 6.85 | 0.55 / 0.24 | 3.80 | 0.50 / 0.19 | 2.86 | 0.33 / 0.19 | 2.65 | 0.44 / 0.15 |
| 2K61 | 0.56 / 0.17 | 2.10 | 0.48 / 0.14 | 1.51 | 0.33 / 0.10 | 0.88 | 0.39 / 0.12 | 0.83 | 0.45 / 0.11 |
| **PRE** | | | | | | | | | |
| 3GB1 | 0.52 | 0.30 | 0.52 | 0.31 | 0.52 | 0.25 | 0.52 | 0.35 | 0.53 |
| 1CDL | 0.46 | 8.58 | 0.44 | 8.01 | 0.35 | 6.64 | 0.35 | 6.94 | 0.36 |
| 1SOS | 0.90 | 6.18 | 0.86 | 2.95 | 0.88 | 6.79 | 0.85 | 2.42 | 0.92 |

* Models used to calculate convergence and the RDC, PCS or PRE Q-factor were identified by the Rosetta full-atom energy.
† Models used to calculate convergence and the RDC, PCS or PRE Q-factor were identified by the combined Rosetta full-atom energy and the respective paramagnetic NMR score term.
‡ ‘Native’ Q-factor was calculated as average over ten models obtained by refining the experimental structure with the Rosetta FastRelax protocol.
§ Converge is defined as the average Cα $RMSD_{100}$ (in Å) between the lowest-scoring model and the next nine low-scoring models, calculated over the secondary structure regions of the protein.

**Table S3:** Protein model refinement with paramagnetic NMR data and Iterative Hybridize; related to **Table 1**.

| PDB ID | Type NMR restraints | $RMSD_{100}$ before refinement | | | $RMSD_{100}$ after refinement | | |
|---|---|---|---|---|---|---|---|
| | | Best RMSD model (Å) | Best scoring model (Å) | Top 10 Score (Å) | Best RMSD model (Å) | Best scoring model (Å) | Top 10 Score (Å) |
| 2KYW | RDC | 4.2 | 5.2 | 9.2 | **2.5** | **3.8** | **3.6** |
| 2KYY | RDC | 3.3 | 5.2 | 9.6 | **2.8** | **3.9** | **4.0** |
| 1CMZ | RDC | 2.6 | 5.9 | 6.2 | 2.9 | **5.3** | **5.2** |
| 2KD7 | RDC | 6.2 | 6.2 | 9.6 | **5.3** | 6.2 | **6.2** |
| 2K5U | RDC | 2.2 | 3.6 | 3.5 | **2.0** | **3.0** | **3.0** |
| 1RW5 | RDC | 3.2 | 4.3 | 11.5 | **1.8** | **2.3** | **2.6** |
| 1RJV | PCS | 3.1 | 6.2 | 6.0 | **2.9** | **4.2** | **4.3** |
| 1J54 | PCS | 2.9 | 3.3 | 7.7 | **2.2** | **2.4** | **2.6** |
| 1CDL | PRE | 3.2 | 4.6 | 7.7 | 8.1 | 10.2 | 10.7 |

The backbone Cα atom $RMSD_{100}$ with respect to the experimental structure before refinement and after running Iterative Hybridize (49) is reported. The $RMSD_{100}$ of the best RMSD and best scoring model and the average $RMSD_{100}$ of the 10 lowest-scoring models was calculated. Improvements in the $RMSD_{100}$ are highlighted in bold font.

**Table S4:** NMR quality factor and convergence of the ten lowest-scoring models of *de novo* predicted symmetric proteins; related to **Figure 5** and **Table 2**.

| PDB ID | no NMR restraints* | | | | with RDC† | | with PCS‡ | | with PRE§ | | Native¶ | | |
|---|---|---|---|---|---|---|---|---|---|---|---|---|---|
| | $Q_{RDC}$ | $Q_{PCS}$ | $Q_{PRE}$ | Convergence# | $Q_{RDC}$ | Convergence | $Q_{PCS}$ | Convergence | $Q_{PRE}$ | Convergence | $Q_{RDC}$ | $Q_{PCS}$ | $Q_{PRE}$ |
| 2KBY | 0.76 | 0.20 | 0.59 | 3.58 | 0.40 | 0.87 | 0.13 | 1.85 | 0.43 | 2.11 | 0.27 | 0.10 | 0.42 |
| 2KO8 | 0.86 | 0.37 | 0.89 | 10.47 | 0.66 | 4.88 | 0.15 | 0.60 | 0.73 | 9.27 | 0.77 | 0.24 | 0.48 |
| 2JWK | 0.71 | 0.37 | 0.64 | 5.36 | 0.46 | 2.75 | 0.18 | 1.70 | 0.58 | 5.40 | 0.42 | 0.16 | 0.52 |
| 2L01 | 0.60 | 0.29 | 0.52 | 4.13 | 0.42 | 2.09 | 0.15 | 0.88 | 0.59 | 5.45 | 0.30 | 0.10 | 0.45 |
| 2LTD | 0.71 | 0.20 | 0.42 | 1.23 | 0.54 | 4.11 | 0.20 | 0.88 | 0.36 | 1.23 | 0.55 | 0.18 | 0.31 |
| 2KOD | 0.51 | 0.28 | 0.49 | 3.08 | 0.44 | 1.41 | 0.21 | 7.86 | 0.27 | 1.42 | 0.38 | 0.14 | 0.39 |
| 1RJJ** | 0.71 | 0.36 | 0.61 | 6.29 | 0.70 | 4.54 | 0.32 | 0.92 | 0.54 | 7.45 | 0.72 | 0.28 | 0.63 |
| 2M89** | 0.59 | 0.34 | 0.99 | 8.67 | 0.55 | 12.94 | 0.32 | 8.11 | 0.72 | 8.83 | 0.37 | 0.21 | 0.52 |

* Models used to calculate convergence and the RDC, PCS or PRE Q-factor were identified by the Rosetta full-atom score.
† Models used to calculate convergence and the RDC Q-factor were identified by the combined Rosetta full-atom and RDC score.
‡ Models used to calculate convergence and the PCS Q-factor were identified by the combined Rosetta full-atom and PCS score.
§ Models used to calculate convergence and the PRE Q-factor were identified by the combined Rosetta full-atom and PRE score.
¶ The 'native' RDC, PCS or PRE Q-factor was calculated as average over ten models obtained by refining the experimental structure with the Rosetta relax protocol.
### Converge is defined as the average Cα $RMSD_{100}$ (in Å) between the lowest-scoring model and the next nine low-scoring models, calculated over the secondary structure regions of the protein.
** For protein targets 1RJJ and 2M89, only the results of the symmetric docking are reported.

**Table S5:** Comparison of benchmark results to those of previous protein structure prediction studies; related to **Table 1** and **2**.

| PDB ID | Present study: Type NMR restraints | Present study: Best RMSD model (Å) | Present study: Best scoring model (Å) | Previous study: Type NMR restraints | Previous study: Best RMSD model (Å) | Previous study: Best scoring model (Å) | Method/ Reference |
|---|---|---|---|---|---|---|---|
| 1Q2N | RDC<br>CS+RDC | **1.0**<br>**1.1** | **1.6**<br>**1.6** | CS+RDC | 4.0 | 5.2* | BCL::Fold (50) |
| 2L7K | RDC<br>CS+RDC | **1.8**<br>**1.7** | **1.9**<br>**2.6** | CS+RDC | 3.3 | 7.4 | |
| 2KYW | RDC<br>CS+RDC<br>CS+RDC | 8.2<br>5.1<br>**2.6[§]** | 14.8<br>6.2<br>**4.2[§]** | CS+RDC | **4.8** | **5.4** | |
| 2KCT | RDC<br>CS+RDC | **2.1**<br>**1.6** | **5.8**<br>**1.8** | CS+RDC | 4.0 | 10.0 | |
| 2H45 | RDC<br>CS+RDC | **1.1**<br>**1.0** | **1.2**<br>**1.1** | CS+RDC | 4.1 | 10.2 | |
| 2KLC | RDC<br>CS+RDC | **1.4**<br>**1.5** | **1.3**<br>**1.9** | CS+RDC | 3.6 | 7.2 | |
| 2JV3 | RDC<br>CS+RDC | **1.8**<br>**1.6** | **2.9**<br>**2.7** | CS+RDC | 2.5 | 5.7 | |
| 2A7O | RDC<br>CS+RDC | **1.6**<br>**1.5** | 4.1<br>**2.7** | CS+RDC | 1.7 | 4.1 | |
| 2KCK | RDC<br>CS+RDC | **2.3**<br>**2.4** | **2.9**<br>**2.6** | CS+RDC | 3.0 | 6.3 | |
| 2KD1 | RDC<br>CS+RDC | **1.5**<br>**1.7** | **2.0**<br>**1.8** | CS+RDC | 2.6 | 5.0 | |
| 2L3W | RDC<br>CS+RDC | **2.4**<br>**2.0** | 3.4<br>**2.4** | CS+RDC | 2.8 | 3.4 | |
| 1CMZ | RDC<br>CS+RDC<br>CS+RDC | **4.2**<br>**2.7**<br>**2.8[§]** | 8,5<br>6.0<br>**5.1[§]** | CS+RDC | 4.4 | **5.7** | |
| 2KYY | RDC<br>CS+RDC<br>CS+RDC | 5.9<br>3.3<br>**2.9[§]** | 12.9<br>6.6<br>**3.0[§]** | CS+RDC | **3.2** | **4.8** | |
| 1RW5 | RDC<br>CS+RDC<br>CS+RDC | **1.5**<br>1.6<br>**1.3[§]** | **1.5**<br>**1.6**<br>**1.6[§]** | CS+RDC | 1.6 | 2.3 | |
| 2AE9 | PCS<br>CS+PCS | -<br>- | 1.7<br>**1.5** | PCS | - | **1.6[†]** | PCS-Rosetta (4) |
| 2K61 | PCS<br>CS+PCS | -<br>- | 2.8<br>3.4 | PCS | - | **2.8** | |
| 1RJV | PCS<br>CS+PCS<br>CS+PCS | -<br>-<br>- | **8.0**<br>**6.4**<br>**4.4[§]** | PCS | - | 11.3 | |
| 1J54 | PCS<br>CS+PCS<br>CS+PCS | -<br>-<br>- | **8.1**<br>**4.2**<br>**3.0[§]** | PCS | - | 20.6 | |
| 3GB1 | CS<br>CS+PRE | -<br>- | 0.7<br>0.7 | CS | - | 0.7[†] | CS-Rosetta (51) |
| 1D3Z | CS<br>CS+PCS | -<br>- | 1.0<br>0.9 | CS | - | **0.7** | |
| 2KCT | RDC<br>CS+RDC | -<br>- | 4.0<br>2.1 | CS+RDC | - | **1.4[‡]** | CS-RDC-Rosetta (52) |
| 2KD1 | RDC<br>CS+RDC | -<br>- | **3.2**<br>**2.5** | CS+RDC | - | 3.4 | |
| 2KCL | RDC<br>CS+RDC | -<br>- | 1.6<br>1.7 | CS+RDC | - | **1.4** | |
| 2K5U | RDC<br>CS+RDC<br>CS+RDC | -<br>-<br>- | 3.9<br>4.4<br>3.7[§] | CS+RDC<br>CS+RDC+NOE | -<br>- | **2.6**<br>**2.5[§]** | |
| 2KD7 | RDC<br>CS+RDC<br>CS+RDC | -<br>-<br>- | 11.2<br>9.9<br>7.3[§] | CS+RDC+NOE | - | **2.4[§]** | |
| 2JWK | RDC<br>PCS<br>PRE | -<br>-<br>- | 4.3<br>1.6<br>11.6 | CS+RDC | - | **1.5[¶]** | CS-RDC-RosettaSymmetry (53) |
| 2KOD | RDC<br>PCS | -<br>- | 1.6<br>4.6 | CS+RDC | - | **1.4** | |

| | | | | | | |
|---|---|---|---|---|---|---|
| | PRE | - | 3.6 | | | |
| 1RJJ | RDC | - | 9.8 | CS+RDC | - | **2.2** |
| | PCS | - | 3.7 | | | |
| | PRE | - | 8.3 | | | |

The RMSD of protein structure models generated in the present study is compared with the RMSD value of the top-scoring and, if available, top-RMSD model reported in previous studies. Lower RMSD values are highlighted in bold font.

* The RMSD reported in the BCL::Fold method (50) was the $RMSD_{100}$ computed over native secondary structure elements. We have recalculated the $RMSD_{100}$ of models in the present study for DSSP-assigned secondary structure regions.

† The RMSD reported in the PCS- (4) and CS-Rosetta (51) methods was the unnormalized RMSD computed for residues in the protein core without termini. We have recalculated the RMSD obtained in the present study for the range of protein core residues reported in the PCS- and CS-Rosetta publications.

‡ The RMSD reported in the CS-RDC-Rosetta method (52) was the median RMSD of the 10 lowest energy structures computed over residues in the protein core or in converged regions of the protein (i.e. those residues for which their backbone can be superimposed within 4 Å) in case of the iterative CS-RDC-Rosetta protocol. Since the authors did not specify the exact set of residues included in the RMSD calculation we compare their results here with the RMSD we obtained for the residue region described in **Table S1**.

§ RMSD obtained after iterative refinement protocol.

¶ The RMSD reported in the CS-RDC-Rosetta method for symmetric proteins (53) was the RMSD of the lowest energy dimer.

### Supporting References


1. Bertini I*, et al.* (2001) Paramagnetism-based versus classical constraints: an analysis of the solution structure of Ca Ln calbindin D9k. *J Biomol NMR* 21(2):85-98.
2. Iwahara J, Schwieters CD, & Clore GM (2004) Ensemble approach for NMR structure refinement against $^{1}$H paramagnetic relaxation enhancement data arising from a flexible paramagnetic group attached to a macromolecule. *J Am Chem Soc* 126(18):5879-5896.
3. Koehler J & Meiler J (2011) Expanding the utility of NMR restraints with paramagnetic compounds: background and practical aspects. *Prog Nucl Magn Reson Spectrosc* 59(4):360-389.
4. Schmitz C, Vernon R, Otting G, Baker D, & Huber T (2012) Protein structure determination from pseudocontact shifts using ROSETTA. *J Mol Biol* 416(5):668-677.
5. Losonczi JA, Andrec M, Fischer MWF, & Prestegard JH (1999) Order Matrix Analysis of Residual Dipolar Couplings Using Singular Value Decomposition. *J. Magn. Res.* 138:334-342.
6. Battiste JL & Wagner G (2000) Utilization of site-directed spin labeling and high-resolution heteronuclear nuclear magnetic resonance for global fold determination of large proteins with limited nuclear overhauser effect data. *Biochemistry* 39(18):5355-5365.
7. Clore GM & Iwahara J (2009) Theory, Practice, and Applications of Paramagnetic Relaxation Enhancement for the Characterization of Transient Low-Population States of Biological Macromolecules and Their Complexes. *Chemical Reviews* 109(9):4108-4139.
8. Lipari G & Szabo A (1982) Model-Free Approach to the Interpretation of Nucllear Resonance Relaxation in Macromolecules. 1. Theory and Range of Validity. *J. Am. Chem. Soc.* 104:4546-4559.
9. Lipari G & Szabo A (1982) Model-Free Approach to the Interpretation of Nucllear Resonance Relaxation in Macromolecules. 2. Analysis of Experimental Results. *J. Am. Chem. Soc.* 104:4559-4570.
10. Brueschweiler R, Roux B, Griesinger C, Karplus M, & Ernst RR (1992) Influence of Rapid Intramolecular Motion on NMR Cross-Relaxation Rates. A Molecular Dynamics Study of Antamanide in Solution. *J. Am. Chem. Soc.* 114:2289-2302.
11. Carugo O & Pongor S (2001) A normalized root-mean-square distance for comparing protein three-dimensional structures. *Protein Sci* 10(7):1470-1473.
12. Kabsch W & Sander C (1983) Dictionary of Protein Secondary Structure: Pattern Recognition of Hydrogen-Bonded and Geometrical Features. *Biopolymers* 22:2577-2637.
13. Cornilescu G, Marquardt JL, Ottiger M, & Bax A (1998) Validation of Protein Strcuture from Anisotropic Carbonyl Chemical Shifts in a Dilute Liquid Crystalline Phase. *J. Am. Chem. Soc.* 120:6836-6837.
14. Chen WN*, et al.* (2016) Sensitive NMR Approach for Determining the Binding Mode of Tightly Binding Ligand Molecules to Protein Targets. *J Am Chem Soc* 138(13):4539-4546.
15. Trott O & Olson AJ (2010) AutoDock Vina: improving the speed and accuracy of docking with a new scoring function, efficient optimization, and multithreading. *J Comput Chem* 31(2):455-461.
16. Nadaud PS, Helmus JJ, Sengupta I, & Jaroniec CP (2010) Rapid acquisition of multidimensional solid-state NMR spectra of proteins facilitated by covalently bound paramagnetic tags. *J Am Chem Soc* 132(28):9561-9563.
17. Olivieri C*, et al.* (2018) Simultaneous detection of intra- and inter-molecular paramagnetic relaxation enhancements in protein complexes. *J Biomol NMR* 70(3):133-140.
18. Fawzi NL*, et al.* (2011) A rigid disulfide-linked nitroxide side chain simplifies the quantitative analysis of PRE data. *J Biomol NMR* 51(1-2):105-114.
19. Zheng D, Aramini JM, & Montelione GT (2004) Validation of helical tilt angles in the solution NMR structure of the Z domain of Staphylococcal protein A by combined analysis of residual dipolar coupling and NOE data. *Protein Sci* 13(2):549-554.
20. Vakonakis I, Staunton D, Rooney LM, & Campbell ID (2007) Interdomain association in fibronectin: insight into cryptic sites and fibrillogenesis. *EMBO J* 26(10):2575-2583.
21. Slupsky CM*, et al.* (1998) Structure of the Ets-1 pointed domain and mitogen-activated protein kinase phosphorylation site. *Proc Natl Acad Sci U S A* 95(21):12129-12134.
22. Li M, Phatnani HP, Greenleaf AL, & Zhou P (2006) NMR assignment of the SRI domain of human Set2/HYPB. *J Biomol NMR* 36 Suppl 1:5.
23. de Alba E, de Vries L, Farquhar MG, & Tjandra N (1999) Solution Structure of Human GAIP (Galpha Interacting Protein): A Regulator of G Protein Signaling. *J. Mol. Biol.* 291:927-939.
24. Liu Y, Kahn RA, & Prestegard JH (2009) Structure and membrane interaction of myristoylated ARF1. *Structure* 17(1):79-87.
25. Teilum K*, et al.* (2005) Solution structure of human prolactin. *J Mol Biol* 351(4):810-823.
26. Mueller GA*, et al.* (2005) Nuclear magnetic resonance solution structure of the Escherichia coli DNA polymerase III theta subunit. *J Bacteriol* 187(20):7081-7089.
27. Schmitz C, Stanton-Cook MJ, Su XC, Otting G, & Huber T (2008) Numbat: an interactive software tool for fitting Deltachi-tensors to molecular coordinates using pseudocontact shifts. *J Biomol NMR* 41(3):179-189.

28. Loh CT, *et al.* (2013) Lanthanide tags for site-specific ligation to an unnatural amino acid and generation of pseudocontact shifts in proteins. *Bioconjug Chem* 24(2):260-268.
29. Ogura K, *et al.* (2008) Solution structure of the Grb2 SH2 domain complexed with a high-affinity inhibitor. *J Biomol NMR* 42(3):197-207.
30. Saio T, *et al.* (2011) An NMR strategy for fragment-based ligand screening utilizing a paramagnetic lanthanide probe. *J Biomol NMR* 51(3):395-408.
31. Yagi H, *et al.* (2013) Three-dimensional protein fold determination from backbone amide pseudocontact shifts generated by lanthanide tags at multiple sites. *Structure* 21(6):883-890.
32. Liepinsh E, *et al.* (2001) Thioredoxin fold as homodimerization module in the putative chaperone ERp29: NMR structures of the domains and experimental model of the 51 kDa dimer. *Structure* 9(6):457-471.
33. Ilangovan U, Ton-That H, Iwahara J, Schneewind O, & Clubb RT (2001) Structure of sortase, the transpeptidase that anchors proteins to the cell wall of Staphylococcus aureus. *Proc Natl Acad Sci U S A* 98(11):6056-6061.
34. DeRose EF, *et al.* (2002) Model for the catalytic domain of the proofreading epsilon subunit of Escherichia coli DNA polymerase III based on NMR structural data. *Biochemistry* 41(1):94-110.
35. Schmitz C, *et al.* (2006) Efficient chi-tensor determination and NH assignment of paramagnetic proteins. *J Biomol NMR* 35(2):79-87.
36. Strickland M, *et al.* (2016) Structure of the NPr:EIN(Ntr) Complex: Mechanism for Specificity in Paralogous Phosphotransferase Systems. *Structure* 24(12):2127-2137.
37. Baig I, *et al.* (2004) Paramagnetism-based refinement strategy for the solution structure of human alpha-parvalbumin. *Biochemistry* 43(18):5562-5573.
38. Bertini I, *et al.* (2009) Accurate solution structures of proteins from X-ray data and a minimal set of NMR data: calmodulin-peptide complexes as examples. *J Am Chem Soc* 131(14):5134-5144.
39. Wilton DJ, Tunnicliffe RB, Kamatari YO, Akasaka K, & Williamson MP (2008) Pressure-induced changes in the solution structure of the GB1 domain of protein G. *Proteins* 71(3):1432-1440.
40. Ikura M, Kay LE, Krinks M, & Bax A (1991) Triple-resonance multidimensional NMR study of calmodulin complexed with the binding domain of skeletal muscle myosin light-chain kinase: indication of a conformational change in the central helix. *Biochemistry* 30(22):5498-5504.
41. Banci L, *et al.* (1998) Solution structure of reduced monomeric Q133M2 copper, zinc superoxide dismutase (SOD). Why is SOD a dimeric enzyme? *Biochemistry* 37(34):11780-11791.
42. Knight MJ, *et al.* (2012) Structure and backbone dynamics of a microcrystalline metalloprotein by solid-state NMR. *Proc Natl Acad Sci U S A* 109(28):11095-11100.
43. Coutandin D, *et al.* (2009) Conformational stability and activity of p73 require a second helix in the tetramerization domain. *Cell Death Differ* 16(12):1582-1589.
44. Sachleben JR, McElroy CA, Gollnick P, & Foster MP (2010) Mechanism for pH-dependent gene regulation by amino-terminus-mediated homooligomerization of Bacillus subtilis anti-trp RNA-binding attenuation protein. *Proc Natl Acad Sci U S A* 107(35):15385-15390.
45. Parsons LM, Grishaev A, & Bax A (2008) The periplasmic domain of TolR from Haemophilus influenzae forms a dimer with a large hydrophobic groove: NMR solution structure and comparison to SAXS data. *Biochemistry* 47(10):3131-3142.
46. Byeon IJ, *et al.* (2009) Structural convergence between Cryo-EM and NMR reveals intersubunit interactions critical for HIV-1 capsid function. *Cell* 139(4):780-790.
47. Cornilescu G, *et al.* (2004) Solution structure of a homodimeric hypothetical protein, At5g22580, a structural genomics target from Arabidopsis thaliana. *J Biomol NMR* 29(3):387-390.
48. Rossi P, *et al.* (2015) A hybrid NMR/SAXS-based approach for discriminating oligomeric protein interfaces using Rosetta. *Proteins* 83(2):309-317.
49. Ovchinnikov S, *et al.* (2017) Protein structure determination using metagenome sequence data. *Science* 355(6322):294-298.
50. Weiner BE, *et al.* (2014) BCL::Fold--protein topology determination from limited NMR restraints. *Proteins* 82(4):587-595.
51. Shen Y, *et al.* (2008) Consistent blind protein structure generation from NMR chemical shift data. *Proceedings of the National Academy of Sciences of the United States of America* 105(12):4685-4690.
52. Raman S, *et al.* (2010) NMR structure determination for larger proteins using backbone-only data. *Science* 327(5968):1014-1018.
53. Sgourakis NG, *et al.* (2011) Determination of the structures of symmetric protein oligomers from NMR chemical shifts and residual dipolar couplings. *J Am Chem Soc* 133(16):6288-6298.
