## Supplementary material for "Integrative protein modeling in RosettaNMR from sparse paramagnetic restraints": RosettaNMR Protocol Capture

### Protocol capture to the manuscript "Integrative protein modeling in RosettaNMR from sparse paramagnetic restraints"

#### Overview

This document is a protocol capture to the manuscript "Integrative protein modeling in RosettaNMR from sparse paramagnetic restraints" by Kuenze G. et al. For following the steps in this protocol capture, it is recommended that the user downloads the weekly release version **v2019.02-dev60579**, which is available at https://www.rosettacommons.org/software/license-and-download.

Instructions on how to install Rosetta, information on software dependencies and install times, and how to get started with the program can be found on the Rosetta Commons documentation webpages: https://www.rosettacommons.org/docs/latest/build_documentation/Build-Documentation and https://www.rosettacommons.org/docs/latest/getting_started/Getting-Started.

This protocol capture contains instructions for applying paramagnetic restraints in the RosettaNMR framework to four different modeling tasks: protein *de no* structure prediction (*protocol 1*), modeling of symmetric proteins (*protocol 2*), protein-ligand docking (*protocol 3*) and protein-protein docking (*protocol 4*). All required input files for running those RosettaNMR calculations can be found in the directory accompanying this protocol capture. Copy the file ***protocol_capture_rosettanmr_inputfiles.tar.gz*** to your desired location and extract and unpack the input file directory by running the command:

```
tar -xzvf protocol_capture_rosettanmr_inputfiles.tar.gz
```

The following conventions are used throughout the protocol capture: ***Names of directories are highlighted in bold and italicized font.*** *File names are written in italicized font.* `And fixed width text means that this is a command that should be typed into the terminal.`

#### 1) De novo structure prediction

This protocol introduces protein *de novo* structure prediction in the RosettaNMR framework with PCS, RDC and PRE data. The protein ubiquitin (PDB ID 1D3Z), for which experimental PCS, RDC [1] and PRE data [2] were published, is used as an example system.
Navigate to the folder ***1_denovo***. There are six sub-folders located, one ***input*** and one ***output*** folder and four additional folders, one for each of the four steps of this protocol:

- fragment picking with chemical shifts (***1_fragments***)
- creating decoys with Rosetta AbRelax without NMR data (***2_abrelax***)
- scoring of decoys with PCS, RDC and PRE data and calculation of the restraint score weight in both centroid and full-atom mode (***3_rescore***)
- structure prediction with Rosetta AbRelax with PCS, RDC and PRE data (***4_abrelax_nmr***)

In the ***input*** folder, the following files are prepared for you:

- Fragment files (*1d3z.200.3mers*, *1d3z.200.9mers*) (will be created during step 1.1)
- Blast checkpoint file (*1d3z.checkpoint*) (will be created during step 1.1)
- List of PDB IDs of homologous proteins; this list can be used to exclude proteins from fragment picking e.g. for benchmarking purposes (*1d3z.deniedpdbs.txt*)
- 1D3Z sequence in Fasta format (*1d3z.fasta*)
- 1D3Z structure in PDB format; only the first model of the NMR ensemble (*1d3z.pdb*)
- Topology broker setup file (*1d3z.tbp*)
- PCS input file (*1d3z.pcs.inp*)

- RDC input file (*1d3z.rdc.inp*)
- PRE input file (*1d3z.pre.inp*)
- PCS data files (in Rosetta format); located in sub-directory ***pcs***; one data file for each spin-labeled variant (T12C and S57C), and each lanthanide ion ($Tb^{3+}$ and $Tm^{3+}$)
- RDC data files (in Rosetta format); located in sub-directory ***rdc***; one data file for each spin-labeled variant (T12C and S57C), and each lanthanide ion ($Tb^{3+}$ and $Tm^{3+}$)
- PRE data file (in Rosetta format); located in directory ***pre***; one $^{1}H$-$R_2$ PRE data file for MTSL spin-labeled variant K48C
- Score function patch files for PCSs, RDCs and PREs (*pcs.wts_patch*, *rdc.wts_patch*, *pre.wts_patch*) (will be created during step 1.3)
- Score function patch file for all NMR data (*nmr.wts_patch*) (will be created in step 1.5)
- Chemical shift table in TALOS+ format (*cs/1d3z.tab*); information on the table format can be found under https://spin.niddk.nih.gov/NMRPipe/talos/
- TALOS+ secondary structure prediction file (*cs/1d3z.predSS.tab*) (will be created during step 1.1)
- TALOS+ phi/psi torsion angle prediction file (*cs/1d3z.pred.tab*) (will be created during step 1.1)

The ***output*** folder contains:
- Binary silent file containing 1000 decoys of 1D3Z (*1d3z_abrelax.out*) generated without NMR data to be used for scoring and NMR score weight calculation in step 1.3.

##### 1.1) Fragment picking with chemical shifts

- Change into the ***1_fragments*** sub-folder. An options file (*fragment_picker.options*) and a weights file (*fragment_picker.wts*) for the Rosetta fragment picker application are provided in this directory. In the options file, adjust the paths to the Rosetta database and to the vall fragment database to the location where they are found on your system. Make sure that the path to all other files in the ***input*** directory is correctly set too.

- Notice that the PSI-BLAST checkpoint file is provided in the ***input*** directory for you. This file can be created by running the two commands below, using blastpgp version 2.2.18 and assuming that PSI-BLAST is correctly installed and the path to the PSI-BLAST database correctly set up.

```
blastpgp -b 0 -j 3 -h 0.001 -d path_to_PSI-BLAST_database -i 1d3z.fasta -C
1d3z.chk -Q 1d3z.ascii
../../scripts/convert_chk.pl 1d3z.fasta 1d3z.chk
```

  The last command converts the checkpoint file from a binary format (*1d3z.chk*) to a human-readable format (*1d3z.checkpoint*) that can be read by Rosetta.

- The secondary structure and phi/psi angle prediction files are located in the ***input*** directory as well. If you want to prepare those files from the chemical shift table yourself, use the program TALOS+ which is free for download (https://spin.niddk.nih.gov/NMRPipe/talos/). Assuming TALOS+ is correctly installed one can simply run the following command in the terminal:

```
talos+ -in 1d3z.tab
```

  Prediction results can be inspected and further refined through the graphical interface rama+.

```
rama+ -in 1d3z.tab
```

  Further information how to select consistent predictions can be found under: https://spin.niddk.nih.gov/NMRPipe/talos/. Alternatively, one can also submit the chemical shift file to the TALOS+ web-server of the Bax group (https://spin.niddk.nih.gov/bax/nmrserver/talos/) for creating secondary structure and phi/psi angle prediction files.

- From the ***1_fragments*** folder, run the Rosetta fragment picker application by typing:

```
~/Rosetta/main/source/bin/fragment_picker.linuxgccrelease @
fragment_picker.options
```

- This will create 3mer and 9mer fragment files (*1d3z.200.3mers*, *1d3z.200.9mers*) and fragment score files (*1d3z.fsc.200.3mers*, *1d3z.fsc.200.9mers*). For convenience, those files are also provided in the ***input*** directory for you. More information on the fragment picker options can be found in [3].

##### 1.2) Creating decoys with Rosetta AbRelax without NMR data

- Change into the ***2_abrelax*** sub-folder. An options file (*abrelax.options*) is provided in this directory. Using the provided settings, one output model will be created. Change the "-out:nstruct" flag to the desired number of models if you wish to create more models. Typically, for *de novo* structure prediction, more than 1000 models are created and calculations are run on multiple CPUs on a cluster.

- Run the Minirosetta application by typing:

```
~/Rosetta/main/source/bin/minirosetta.linuxgccrelease -database
~/Rosetta/main/database/ @ abrelax.options
```

- This will create one output model in Rosetta silent file format (*1d3z_abrelax.out*) and a Rosetta score file (*1d3z_abrelax.sc*). Extract the model of 1D3Z as PDB file by typing:

```
~/Rosetta/main/source/bin/extract_pdbs.linuxgccrelease -in:file:silent
1d3z_abrelax.out
```

- Note, a Rosetta silent file (*1d3z_abrelax.out*) containing 1000 models of 1D3Z created without paramagnetic restraints but with fragments selected from chemical shifts is provided in the ***output*** directory. If you wish, you can use this file to proceed to the next step.

##### 1.3) Scoring of decoys with PCS, RDC and PRE data

- Change into the sub-folder ***3_rescore***. Individual settings files for scoring with either PCSs (*rescore.pcs.options*), RDCs (*rescore.rdc.options*) or PREs (*rescore.pre.options*) can be found in this directory. In addition, two Rosetta XML scripts are provided: For scoring with the score3 centroid-atom energy function use the script *rescore.cen.xml*, and for scoring with the Rosetta ref2015 full-atom energy function use the script *rescore.fa.xml*.
  For running structure prediction with the AbRelax protocol (step 1.4), we will use a weight of the PCS, RDC and PRE score that is optimized against the score3 energy, and for final model scoring and selection the weight will be adjusted relative to the total ref2015 energy. The following formula is used for the weight calculation:

$$w_{NMR} = \frac{E_{Rosetta}^{high} - E_{Rosetta}^{low}}{E_{NMR}^{high} - E_{NMR}^{low}} \quad (1)$$

where $E_{Rosetta}^{high}$ and $E_{Rosetta}^{low}$ are the average highest and lowest 10% of the values of the Rosetta score, and $E_{NMR}^{high}$ and $E_{NMR}^{low}$ are the average of the highest and lowest 10% of the values of the PCS, RDC or PRE score, respectively.

- Have a look at the PCS, RDC and PRE options files by opening these files in a text editor. The most important parameters that control how the PCS, RDC or PRE score is calculated are written in separate input files (named *1d3z.pcs.inp*, *1d3z.rdc.inp* and *1d3z.pre.inp*, respectively, and located in the ***input*** directory). Take a look at these input files by opening them in another text editor window. PCS and PRE datasets that were measured at different spin-label sites or with different types of spin-labels and RDC datasets collected with different alignment media are grouped in separate sections in the input file. Each section must start with the keyword "**MULTISET**" and end with the word "**END**". The calculation parameters are specified as key-value pairs, and each pair must be separated by an equal ("=") sign. Read the comments in the input files for a short description of the meaning and format of the computation parameters. Note, blank lines and comments (starting with a # sign) are allowed within the MULTISET section but not on the same line as a computation parameter. The format of each type of input file and the complete list of parameters is explained in more detail in **Table 1** in the **Appendix** at the end of this document. Furthermore, a collection of optional flags provide additional control over how the respective scores are calculated. Those options and their meaning are summarized in **Table 2** in the **Appendix**.

- The experimental PCS, RDC and PRE values can be found in the data files in the ***input/pcs***, ***input/rdc*** and ***input/pre*** directory, respectively. The data files follow a specific Rosetta format which is explained in the respective file header. Briefly, each experimental value must be placed on a new line. The atom to which a measured PCS, RDC or PRE value refers is defined by its name, it's residue number and chain ID. This atom definition must be enclosed in parentheses and must appear in front of the experimental value. The last two elements in each line are the experimental value and error in ppm (PCSs) or Hz (RDCs or PREs).

- Notice that the PCS, RDC and PRE options files contain a flag called "multiset_weights" which is followed by a vector of floating point numbers. Those numbers represent weighting factors which specify how much the score values that are calculated for the individual tagging sites or alignment media will contribute to the overall score. Thus, different experimental datasets can be assigned a higher or lower weight in order to reflect our decisions regarding their quality or importance for the calculation. Two important remarks about the input format of this flag must be mentioned: (1) the vector size must be equal the number of PCS, PRE or RDC MULTISETs, i.e. 2, 1 and 4 for this example, and (2) the order by which the weighting factors are applied will be the same as the order in which MULTISETs appear in the NMR input file.

- For scoring, the RosettaScripts application is used. Run the following command in the terminal:

```
~/Rosetta/main/source/bin/rosetta_scripts.linuxgccrelease -database
~/Rosetta/main/database/ -parser:protocol rescore.cen.xml @ rescore.pcs.options
-in:file:silent 1d3z_abrelax.out -out:file:scorefile 1d3z_abrelax_pcs_score3.sc
```

- By adding the "-in:file:tags" flag to the command above followed by one or more model tags, specific models can be selected for scoring (e.g. `-in:file:tags 1d3z_abrelax_1_S_0001`).

- Repeat scoring with the score3 weights for PREs and RDCs.

- Repeat scoring with the ref2015 score function. Therefore, in the options file, remove the pound sign (#) in front of the ref2015-specific flags and comment out those flags specific for the score3 score function. For example, the corresponding lines in your PCS options file should look like this:

```
# Use the following options for scoring with Rosetta high resolution score
function
-parser:script_vars sfxn=ref2015 nmr_sc_type=nmr_pcs nmr_sc_wt=1.0
-score:weights ref2015.wts

# Use the following options for scoring with Rosetta low resolution score
function
#-parser:script_vars sfxn=score3.wts nmr_sc_type=nmr_pcs nmr_sc_wt=1.0
#-score:weights score3.wts
```

- Now, type the following command:

```
~/Rosetta/main/source/bin/rosetta_scripts.linuxgccrelease -database
~/Rosetta/main/database/ -parser:protocol rescore.fa.xml @ rescore.pcs.options -
in:file:silent 1d3z_abrelax.out -out:file:scorefile 1d3z_abrelax_pcs_ref2015.sc
```

- Repeat scoring with the ref2015 weights for PREs and RDCs.

- *Optional step:* If you have split up the *de novo* structure calculation over multiple CPUs, created multiple silent output files and scored each of them separately with the commands above, concatenate all separate score files into one single text file that contains the scores of all models. For example:

```
cat 1d3z_abrelax_*_pcs_ref2015.sc > 1d3z_abrelax_pcs_ref2015.sc
```

- Calculate the PCS, RDC or PRE weight for the AbRelax protocol by running the *calc_nmr_wt.py* script from the scripts directory with the score files obtained by scoring with the score3 weight set. The script requires as arguments the name of the Rosetta score file and at least one NMR score type (*pcs*, *rdc* or *pre*).

```
../../scripts/calc_nmr_wt.py 1d3z_abrelax_pcs_score3.sc --nmr_scoretypes pcs
../../scripts/calc_nmr_wt.py 1d3z_abrelax_rdc_score3.sc --nmr_scoretypes rdc
../../scripts/calc_nmr_wt.py 1d3z_abrelax_pre_score3.sc --nmr_scoretypes pre
```

Note: when copying this command, make sure that the editor does not replace the en dash (-) with the em dash (–) in front of the "--nmr_scoretypes" flag as this might cause an error in the script.

A short message reporting the calculated weight will be written to the terminal, e.g:

```
nmr_rdc = 0.0727
```

- Additionally, calculate the PCS, RDC or PRE weight to be used for final model scoring and selection by running the *calc_nmr_wt.py* script with the score files obtained by scoring with the ref2015 weight set:

```
../../scripts/calc_nmr_wt.py 1d3z_abrelax_pcs_ref2015.sc --nmr_scoretypes pcs
../../scripts/calc_nmr_wt.py 1d3z_abrelax_rdc_ref2015.sc --nmr_scoretypes rdc
../../scripts/calc_nmr_wt.py 1d3z_abrelax_pre_ref2015.sc --nmr_scoretypes pre
```

**1.4) *De novo* structure prediction with Rosetta AbRelax and PCSs, RDCs and PREs**

- Navigate to the sub-folder ***4_abrelax_nmr***. There are eight options files located in this directory: four for running AbRelax and four for rescoring. The options files named *abrelax.pcs.options*,

*abrelax.rdc.options* and *abrelax.pre.options*, are for running AbRelax with PCSs, RDCs and PREs, respectively, and the options file named *abrelax.nmr.options* can be used for structure prediction with all three types of NMR data applied together.
Take a look at these files by opening them in a text editor and compare them to the *abrelax.options* file located in the ***2_abrelax*** folder. PCS, RDC and PRE restraints are applied in Abinitio stages 1 – 4 by setting the weight of these score terms in the Rosetta score function. We do this by changing their weights in the accompanying patch files (*pcs.wts_patch*, *rdc.wts_patch*, *pre.wts_patch* in the ***input*** folder) to those values that were determined for centroid mode in the previous step. For example, the line in the *pcs.wts_patch* file:

```
nmr_pcs = 3.0
```

sets the weight of the PCS score to 3.0.

- After modifying the respective score function patch files, run the Minirosetta application with PCS, RDC or PRE restraints by typing:

```
~/Rosetta/main/source/bin/minirosetta.linuxgccrelease -database
~/Rosetta/main/database/ @ abrelax.pcs.options
```

- This will create one output model in Rosetta silent file format (*1d3z_abrelax_pcs.out*) and a Rosetta score file (*1d3z_abrelax_pcs.sc*). Increase the value of the "-out:nstruct" flag to create more models. Typically, more than 1000 models are created for *de novo* structure prediction and calculations are run on multiple CPUs on a cluster.

- For final model selection, rescore models with PCS, RDC or PRE data. In this step, we can use the Rosetta *score_jd2* command line application. Within the provided options files (*rescore.pcs.options*, *rescore.rdc.options*, *rescore.pre.options*), change the PCS, RDC or PRE score weight to the value determined for the Rosetta ref2015 score function in step 1.3. For example, the following line in the *rescore.pcs.options* file sets the weight for the PCS score to 2.6:

```
-score:set_weights nmr_pcs 2.6
```

- Then, run the following commands in the terminal:

```
~/Rosetta/main/source/bin/score_jd2.linuxgccrelease -database
~/Rosetta/main/database/ @ rescore.pcs.options -in:file:silent
1d3z_abrelax_pcs.out -out:file:scorefile 1d3z_abrelax_pcs_rescored.sc

~/Rosetta/main/source/bin/score_jd2.linuxgccrelease -database
~/Rosetta/main/database/ @ rescore.rdc.options -in:file:silent
1d3z_abrelax_rdc.out -out:file:scorefile 1d3z_abrelax_rdc_rescored.sc

~/Rosetta/main/source/bin/score_jd2.linuxgccrelease -database
~/Rosetta/main/database/ @ rescore.pre.options -in:file:silent
1d3z_abrelax_pre.out -out:file:scorefile 1d3z_abrelax_pre_rescored.sc
```

- Finally, the Rosetta and NMR score and the model's RMSD relative to the experimental structure of 1D3Z can be extracted from the score file by using the python script *extract_scores.py* from the script directory. The script writes a comma-separated data table for easy creation of Score-vs-RMSD plots (see **Figure 1**). It requires two command line arguments as inputs: 1) the name of the score file and 2) a list of column labels to be extracted from the score file.

```
../../scripts/extract_scores.py 1d3z_abrelax_pcs_rescored.sc --columns score
nmr_pcs rms description && mv scores.csv scores_pcs.csv
../../scripts/extract_scores.py 1d3z_abrelax_rdc_rescored.sc --columns score
nmr_rdc rms description && mv scores.csv scores_rdc.csv
../../scripts/extract_scores.py 1d3z_abrelax_pcs_rescored.sc --columns score
nmr_pre rms description && mv scores.csv scores_pre.csv
```

Note: when copying this command, make sure that the editor does not replace the en dash (-) with the em dash (–) in front of the "--columns" flag as this might cause an error in the script.

- Alternatively, the lowest-scoring model can be selected and used as reference model for the RMSD calculation. For example, assuming the tag of the best-scoring model is 1d3z_abrelax_pcs_S_0010 the following command will extract this model as a PDB file from the respective Rosetta silent file:

```
~/Rosetta/main/source/bin/extract_pdbs.linuxgccrelease -in:file:silent
1d3z_abrelax_pcs.out -in:file:tags 1d3z_abrelax_pcs_S_0010
```

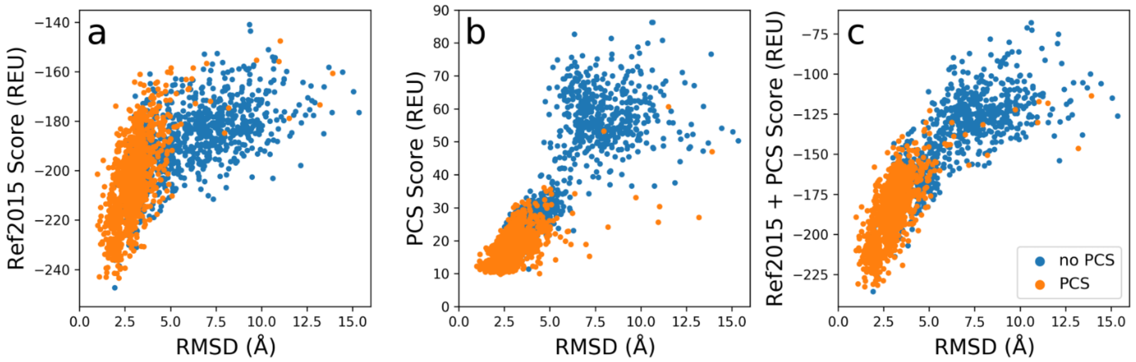


**Figure 1: Structure prediction of 1D3Z with and without PCS data.** The Rosetta ref2015 score (**a**), PCS score (**b**) and combined ref2015 and PCS score (**c**) is plotted versus the Cα-RMSD relative to the experimental structure of 1D3Z.

##### *1.5) Optional step: Repeat structure prediction with Rosetta AbRelax using all NMR data together*

- Combining PCSs, RDCs and PREs in the RosettaNMR framework is as easy as combining their respective options. For running AbRelax with all NMR data, another options file, named *abrelax.nmr.options*, has been prepared for you in ***4_abrelax_nmr***. In addition, within the ***input*** directory, create a new patch file, called *nmr.wts_patch*, which contains the weight of each of the three score terms to be used for the AbRelax calculation:

```
nmr_pcs = 1.000
nmr_rdc = 0.024
nmr_pre = 0.005
```

- Different ways exist of how to split the weights between the three score terms. Here, we choose to set the weight of each NMR score to 1/3 of its original value that was used when PCSs, RDCs and PREs were applied separately. Another possibility is to adjust the weights such that their ratio is proportional to the logarithm of the number of PCS, RDC and PRE values. Furthermore, you may decide adjusting those weights to reflect your considerations on other factors such as confidence in the experimental data or the structural information content of each NMR data type.

- Rerun Rosetta AbRelax with PCSs, RDCs and PREs, and rescore models with the ref2015 score function and NMR data:

```
~/Rosetta/main/source/bin/minirosetta.linuxgccrelease -database
~/Rosetta/main/database/ @ abrelax.nmr.options

~/Rosetta/main/source/bin/score_jd2.linuxgccrelease -database
~/Rosetta/main/database/ @ rescore.nmr.options -score:set_weights nmr_pcs 0.870
nmr_rdc 0.021 nmr_pre 0.006 -in:file:silent 1d3z_abrelax_nmr.out -
out:file:scorefile 1d3z_abrelax_nmr_rescored.sc
```

#### 2) Modeling symmetric proteins

This protocol introduces *de novo* structure prediction of symmetric proteins with the RosettaNMR framework using chemical shifts and RDC data. As example, the C2-symmetric protein 2JWK, for which experimental RDC data were published [4], is used.
Navigate to the folder ***2_symmetry***. There are six sub-folders located, one ***input*** and one ***output*** folder and four additional folders, one for each of the four steps of this protocol:

- fragment picking with chemical shifts (***1_fragments***)
- creating decoys with Rosetta Fold-and-Dock (***2_folddock***)
- scoring of decoys with RDCs and calculation of the RDC score weight (***3_rescore***)
- structure prediction with Rosetta Fold-and-Dock and RDCs (***4_folddock_nmr***)

In the ***input*** folder, the following files are prepared for you:

- Fragment files (*2jwk.200.3mers*, *2jwk.200.9mers*) (will be created during step 2.1)
- Blast checkpoint file (*2jwk.checkpoint*) (will be created during step 2.1)
- List of PDB IDs of homologous proteins; those proteins can be excluded from fragment picking e.g. for benchmarking purposes (*2jwk.deniedpdbs.txt*)
- Sequence of 2JWK chain A in Fasta format (*2jwk.fasta*)
- 2JWK structure in PDB format; only the first model of the NMR ensemble (*2jwk.AB.pdb*)
- Topology broker setup file (*2jwk.tbp*)
- Symmetry definition file for C2-symmetry (*2jwk.symm*) (will be created during step 2.2)
- RDC input file (*2jwk.rdc.inp*)
- RDC data files (in Rosetta format); located in sub-directory ***rdc***; one RDC data file for each of the following atom types: CA-CO, N-H, CA-HA and N-CO.
- Chemical shift file in TALOS+ format (*cs/2jwk.tab*); information on the table format can be found under https://spin.niddk.nih.gov/NMRPipe/talos/
- TALOS+ secondary structure prediction file (*cs/2jwk.predSS.tab*) (will be created during step 2.1)
- TALOS+ phi/psi torsion angle prediction file (*cs/2jwk.pred.tab*) (will be created during step 2.1)
- Score function patch file for RDCs (*rdc.wts_patch*) (will be created during step 2.3)

The ***output*** folder contains:

- Binary silent file containing 1000 decoys of 2JWK (*2jwk_folddock.out*) generated without NMR data to be used for scoring and RDC score weight calculation in step 2.2.

##### 2.1) Fragment picking with chemical shifts

- Change into the ***1_fragments*** sub-folder. An options file (*fragment_picker.options*) and weights file (*fragment_picker.wts*) are provided in this directory. In the options file, adjust the paths to the Rosetta database and to the vall fragment database to the location where they are found on your system. Make sure that the paths to all other files in the ***input*** directory are correctly set too. The secondary structure and phi/psi angle prediction files were created with the help of chemical shifts using the program TALOS+ as explained in the previous protocol (step 1.1). The PSI-BLAST checkpoint file can be created by running the following two commands, assuming that PSI-BLAST is correctly installed and the path to the PSI-BLAST database correctly set up. For convenience, this file has been prepared in the ***input*** directory too.

```
blastpgp -b 0 -j 3 -h 0.001 -d path_to_PSI-BLAST_database -i 1d3z.fasta -C
1d3z.chk -Q 1d3z.ascii
../../scripts/convert_chk.pl 1d3z.fasta 1d3z.chk
```

- Run the Rosetta fragment picker application:

```
~/Rosetta/main/source/bin/fragment_picker.linuxgccrelease @
fragment_picker.options
```

- This will create 3mer and 9mer fragment files (*2jwk.200.3mers*, *2jwk.200.9mers*) and fragment score files (*2jwk.fsc.200.3mers*, *2jwk.fsc.200.9mers*).

**2.2) Creating decoys with Rosetta Fold-and-Dock**

- Navigate to the sub-directory ***2_folddock***. Have a look at the options file that is located in this directory. The last option at the bottom of the file is the path to the symmetry definition file which is needed for modeling symmetric proteins. It contains all the information that Rosetta needs to know about the symmetry of the system, e.g. how to score the symmetric structure, how to maintain symmetry in rigid body perturbations, what degrees of freedom are allowed to move, how to initially setup the symmetric system and how to perturb the system.

- Create a C2-symmetry definition file for a protein with two subunits and cyclical symmetry by running the following python command. Rename the output file to *2jwk.symm*.

```
~/Rosetta/main/source/src/apps/public/symmetry/make_symmdef_file_denovo.py -
symm_type cn -nsub 2
```

  Execute this script without any arguments to see a full list of all possible options. For further information on Rosetta Symmetry the user is referred to reference [5].

- For structure prediction, the Rosetta Fold-and-Dock protocol is used which can model proteins with intertwined topology. Start Fold-and-Dock by running the Minirosetta application:

```
~/Rosetta/main/source/bin/minirosetta.linuxgccrelease -database
~/Rosetta/main/database/ @ folddock.options
```

- This will create one output model in the Rosetta silent file format. If you wish to produce more models increase the value of the “-out:nstruct” flag. Typically, more than 1000 models are created for *de novo* structure prediction and calculations are run on multiple CPUs on a cluster.

- A Rosetta silent file (*2jwk_folddock.out*) containing 1000 models of 2JWK created without RDCs but with fragments selected with chemical shifts is provided in the ***output*** directory.

**2.3) Scoring of decoys with RDC data**

- Change to the ***3_rescore*** directory. An options file (*rescore.rdc.sym.options*) and two Rosetta XML scripts (*rescore.cen.sym.xml, rescore.fa.sym.xml*) are provided for determining the RDC score as well as the Rosetta ref2015 and score3 energy. The weight of the RDC score is calculated according to equation 1 as explained in the previous protocol. For structure prediction with Fold-and-Dock it will be adjusted relative to the score3 energy and for final model scoring and model selection relative to the ref2015 energy.

- In the RosettaNMR framework, the RDC score of symmetric proteins can be calculated by enforcing that one axis of the alignment tensor coincides with the symmetry axis of the system. This assumption is met for proteins with cyclical and dihedral symmetry. In this case, the RDC score is calculated by considering all symmetric copies of the protomer. However, since identical residues in symmetric subunits will share the same experimental RDC value, only RDCs of the asymmetric unit need to be included in the data input file. Have a look at one of the RDC data files (e.g. *2jwk.nh.dat* in ***input/rdc***) to make sure that RDC values are assigned only to residues of chain A. In addition, the "-nmr:rdc:use_symmetry_calc" flag should be set to true in the options file which turns on the symmetric mode of the RDC calculation.

- Symmetric RDC score calculation requires that the orientation of the symmetry axis and the rigid body transformation between symmetric subunits is known. Unfortunately, this information cannot be easily recovered from the Rosetta silent file once a model has been output. To restore this information, we employ the DetectSymmetry Mover in the Rosetta script which initializes a symmetric system from a PDB file. It currently works for all cyclic symmetries from C2 to C99.

- First, extract all protein models from the silent file and create PDB files:

```
~/Rosetta/main/source/bin/extract_pdbs.linuxgccrelease -in:file:silent
2jwk_folddock.out
```

- Then, run the RosettaScripts application for model rescoring. Use the provided options file *rescore.rdc.sym.options.* Make sure to remove any pound signs in front of the following lines:

```
-parser:script_vars sfxn=score3 nmr_sc_type=nmr_rdc nmr_sc_wt=1.0
-score:weights score3.wts
```

  and comment out the respective lines for the ref2015 score function. For scoring with the score3 centroid score function type:

```
~/Rosetta/main/source/bin/rosetta_scripts.linuxgccrelease -parser:protocol
rescore.cen.sym.xml @ rescore.rdc.sym.options -in:file:s 2jwk_folddock_*.pdb -
out:file:scorefile 2jwk_folddock_rdc_score3.sc
```

  For scoring with the ref2015 all-atom score function, make sure to remove any pound signs in front of the following lines in the options file:

```
-parser:script_vars sfxn=ref2015 nmr_sc_type=nmr_rdc nmr_sc_wt=1.0
-score:weights ref2015.wts
```

  and comment out the respective lines for the score3 score function. Then type

```
~/Rosetta/main/source/bin/rosetta_scripts.linuxgccrelease -parser:protocol
rescore.fa.sym.xml @ rescore.rdc.sym.options -in:file:s 2jwk_folddock_*.pdb -
out:file:scorefile 2jwk_folddock_rdc_ref2015.sc
```

- Calculate the RDC weight for the Fold-and-Dock protocol by running the *calc_nmr_wt.py* script from the scripts directory with the score file obtained by scoring with the score3 weight set. The script requires two arguments: 1) the name of the Rosetta score file and 2) the type of NMR restraints (*rdc*).

```
../../scripts/calc_nmr_wt.py 2jwk_folddock_rdc_score3.sc --nmr_scoretypes rdc
```

- Additionally, calculate the RDC weight to be used for final model scoring and selection by running the *calc_nmr_wt.py* script with the score file obtained by scoring with the ref2015 score function:

```
../../scripts/calc_nmr_wt.py 2jwk_folddock_rdc_ref2015.sc --nmr_scoretypes rdc
```

**2.4) *De novo* structure prediction with Rosetta Fold-and-Dock and RDCs**

- Change into the ***4_folddock_nmr*** sub-directory. Have a look at the provided options file (*folddock.rdc.options*). RDCs are applied in Abinitio stages 1 – 4 by setting their score weight in the Rosetta score function to that value determined for the centroid mode in the previous step. You need to modify the accompanying patch file *rdc.wts_patch* in the ***input*** directory to reflect this change, e.g.:

```
nmr_rdc = 0.020
```

- Run the Minirosetta application to start Fold-and-Dock with RDCs.

```
~/Rosetta/main/source/bin/minirosetta.linuxgccrelease -database
~/Rosetta/main/database/ @ folddock.rdc.options
```

- The output Rosetta silent file (*2jwk_folddock_rdc.out*) and score file (*2jwk_folddock_rdc.sc*) will contain one model. Increase the "-out:nstruct" flag to create more models if desired.

- After the structure prediction run is completed, rescore models with RDCs as described in step 2.3). Copy the *rescore.fa.sym.xml* and *rescore.rdc.sym.options* files from folder ***3_rescore*** to the current directory. Make sure to change the RDC weight to that value determined relative to the ref2015 weight set. For example, the following line in the options file sets the weight for the RDC score to 0.02:

```
-parser:script_vars sfxn=ref2015 nmr_sc_type=nmr_rdc nmr_sc_wt=0.02
```

- First, extract models from the silent file and create PDB files:

```
~/Rosetta/main/source/bin/extract_pdbs.linuxgccrelease -in:file:silent
2jwk_folddock_rdc.out
```

- Then, run the RosettaScripts application:

```
~/Rosetta/main/source/bin/rosetta_scripts.linuxgccrelease -parser:protocol
rescore.fa.sym.xml @ rescore.rdc.sym.options -in:file:s 2jwk_folddock_rdc_*.pdb
-out:file:scorefile 2jwk_folddock_rdc_rescored.sc
```

- Finally, the Rosetta and RDC score and the model's symmetric RMSD relative to the experimental structure of 2JWK can be extracted from the score file by using the python script *extract_scores.py* which writes a comma-separated data table for easy creation of Score-vs-RMSD plots (see **Figure 2**). The script requires two command line arguments as inputs: 1) the Rosetta score file and 2) a list of column labels for scores or other metrics that should be extracted.

```
../../scripts/extract_scores.py 2jwk_folddock_rdc_rescored.sc --columns score
nmr_rdc symmetric_rms description
```

- Alternatively, the lowest-scoring model can be selected and used as reference model for the RMSD calculation. E.g., assuming the tag of the lowest-scoring model is 2jwk_folddock_rdc_S_0010 the following command will extract this model as a PDB file from the respective Rosetta silent file:

```
~/Rosetta/main/source/bin/extract_pdbs.linuxgccrelease -in:file:silent
2jwk_folddock_rdc.out -in:file:tags 2jwk_folddock_rdc_S_0010
```

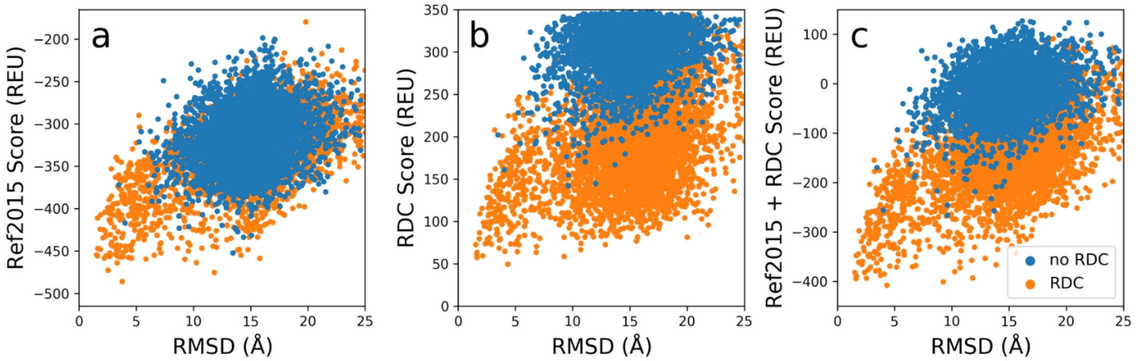


**Figure 2: Structure prediction of the C2-symmetric protein 2JWK with and without RDC data.** The Rosetta ref2015 score (**a**), RDC score (**b**) and combined ref2015 and RDC score (**c**) is plotted versus the Cα-RMSD relative to the experimental structure of 2JWK.

**3) Protein-Ligand Docking**

This protocol introduces how to use RosettaNMR together with PCSs for protein-ligand docking. A complex between the SH2 domain of protein GRB2 and a phosphorylated tripeptide pYTN serves as example system. For comparison, the X-ray structure of GRB2 SH2 with a related phosphorylated nona-peptide bearing the same 3-residue recognition motif (PDB ID 1JYR) is used.

Navigate to the folder ***3_ligand_docking***. There are six sub-folders located in this directory, one ***input*** and one ***output*** folder and four additional folders, one for each of the four steps of this protocol:

- creation of ligand parameters (***1_ligand_params***)
- determination of the Δχ-tensor (***2_tensor_fit***)
- ligand rigid-body grid search with PCSs (***3_rbsearch***)
- full flexible protein-ligand docking with PCSs (***4_docking***)

The ***input*** directory provides the following input files:

- X-ray structure of GRB2 SH2 without pYTN ligand (*GRB2.pdb*); use this for docking runs
- X-ray structure of GRB2 SH2 with pYTN ligand (*GRB2_YTN.pdb*); use this to compare docking results and calculate the ligand RMSD; this PDB was created by superimposing pYTN onto the corresponding residues of the nona-peptide ligand in the experimental structure (PDB ID 1JYR) and replacing the native peptide ligand. The ligand residue is labeled with a different chain ID than the protein, typically chain X.
- PDB file of pYTN (*YTN.pdb*)
- SDF file of pYTN (*YTN.sdf*)
- SDF file containing 1000 pYTN conformations (*YTN_confs.sdf*) (will be created during step 3.1)
- Rosetta residue parameter file for pYTN (*YTN.params*) (will be created during step 3.1)
- Rosetta conformer library for pYTN (*YTN_conformers.pdb*) (will be created during step 3.1)
- Extra parameters for improper torsions (optional) (*YTN.tors*) (will be created during step 3.1)
- PCS input file for GRB2 SH2 (*grb2_pcs.inp*) (Δχ-tensor parameters will be determined in step 3.2)
- PCS input file for pYTN (*ytn_pcs.inp*) (Δχ-tensor parameters will be determined in step 3.2)

- PCS data files (in Rosetta format); located in sub-directory ***pcs***; four PCS data files for GRB2 SH2 (one for each lanthanide: $Tm^{3+}$, $Tb^{3+}$, $Dy^{3+}$, $Er^{3+}$) and three PCS data files for pYTN (one for each lanthanide: $Tm^{3+}$, $Tb^{3+}$, $Dy^{3+}$)

The ***output*** directory contains:

- $\Delta\chi$-tensor values for lanthanides $Tm^{3+}$, $Tb^{3+}$, $Dy^{3+}$, $Er^{3+}$ (*pcs_tensor.info*)
- PDB file containing the coordinates of the pYTN center-of-mass after running the PCS grid search (*YTN_rbseach_centroids.pdb*); this file is used as input for RosettaLigand and parsed to the StartFrom Mover.

##### 3.1) Creation of ligand parameters

- Navigate to the ***1_ligand_params*** directory. Create an SDF file containing the pYTN conformations (*YTN_confs.sdf*). This file is also provided in the input directory. This set of conformations was created with the Meiler lab BioChemicalLibrary (BCL v3.6.1, http://www.meilerlab.org/bclcommons) by running the command below. Notice that the BCL version may differ on your system, and use the help command to print the list of input flags for the ConformerGenerator application.

```
~/BCL/bcl.exe molecule:ConformerGenerator -rotamer_library cod -top_models 1000
-max_iterations 30000 -add_h -ensemble_filenames ../input/YTN.sdf -
conformation_comparer SymmetryRMSD 0.250 -generate_3D -conformers_single_file
YTN_confs.sdf
```

- Parameter files contain information about Rosetta residue types such as atom types, atom partial charges, atom connectivity and the residue's conformation in internal coordinates. Type the following command into the terminal:

```
~/Rosetta/main/source/scripts/python/public/molfile_to_params.py -n YTN -p YTN -
-conformers-in-one-file ../input/YTN_confs.sdf --extra_torsion_output --
recharge=-1
```

  This will create a parameter file (*YTN.params*) for a residue with three-letter code YTN. The program reads all pYTN ligand conformations stored in the file *YTN_confs.sdf* and creates a conformer library in Rosetta format (*YTN_conformers.pdb*). The first ligand conformation is written to a PDB file named *YTN.pdb*. In addition, a text file with extra improper torsions (*YTN.tors*) is generated. The total charge of the ligand is set to -1 and all atom charges are offset accordingly.

- For convenience, all files created in this step are also provided for you in the ***input*** directory so that you can proceed with the subsequent steps in case there are any problems in creating ligand parameters.

##### 3.2) $\Delta\chi$-tensor determination from protein PCSs

- The position of the ligand during docking can be scored by calculating its PCSs at the current coordinates using a predefined $\Delta\chi$-tensor as input and comparing the calculated PCSs with the experimental ones. The values of the $\Delta\chi$-tensor are determined in this step by fitting PCSs to the known structure of the protein. This has the advantage that PCSs of the protein are more manifold than ligand PCSs and thus the $\Delta\chi$-tensor can be determined more accurately.

- Change into the sub-folder ***2_tensor_fit*** and run the following command line application with the options file that is provided in this directory. Make sure that the path to the PDB and PCS input file of GRB2 SH2, which are both located in the ***input*** directory, is correct.

```
~/Rosetta/main/source/bin/calc_nmr_tensor.linuxgccrelease @ fit.options
```

To display a list of all relevant options that this application can take, rerun the command above without the options file but only the "`-help`" flag.
The application creates two output files: a table of experimental-vs.-calculated PCS values is written to file *GRB2_pcs_pred.txt* and the Δχ-tensor values for each lanthanide ion are written to file *pcs_tensor.info* (see below).

| Position | Metal | Experiments | PCSs | Xax | Xrh | alpha | beta | gamma | xM | yM | zM |
|---|---|---|---|---|---|---|---|---|---|---|---|
| 18 | Tb | 1 | 56 | 23.966 | 15.288 | 79.312 | 68.199 | 151.578 | 7.585 | 24.964 | 8.048 |
| 18 | Tm | 1 | 56 | 19.071 | 6.842 | 162.631 | 113.645 | 170.187 | 7.585 | 24.964 | 8.048 |
| 18 | Dy | 1 | 55 | -21.164 | -11.060 | 131.562 | 136.607 | 140.271 | 7.585 | 24.964 | 8.048 |
| 18 | Er | 1 | 57 | 8.102 | 3.451 | 151.963 | 121.910 | 174.273 | 7.585 | 24.964 | 8.048 |

$\chi_{ax}$ and $\chi_{rh}$ are the axial and rhombic component of the Δχ-tensor, α, β and γ are the three Euler angles that orient the tensor frame with respect to the PDB frame, and $x_M$, $y_M$ and $z_M$ are the Cartesian coordinates of the metal ion in the PDB frame.
In addition, another text file (*GRB2_pcs_pred.txt*) containing the predicted PCS values of GRB2 will be created. The last column in this prediction file lists the deviation between predicted and experimental values and indicates positions where the deviation is larger than the error which was provided in the PCS data file.

- In the ligand PCS input file (*ytn_pcs.inp*), change now the Δχ-tensor values for $Tm^{3+}$, $Tb^{3+}$ and $Dy^{3+}$ in the dataset vector to those values determined by the fit. The Δχ-tensor values correspond to the last eight fields in the dataset vector and have the following order from left to right: $x_M$, $y_M$, $z_M$, $\chi_{ax}$, $\chi_{rh}$, α, β, γ. Notice that the "fixed_tensor" keyword in the input file is set to true which means that the Δχ-tensor values are read from the input file and no fitting will be done during ligand docking. The PCS input file for the ligand should look like this:

```
MULTISET

spinlabel_position = 18
chain_id = A
gridsearch = [ CA, CB, 10.0, 4.0, 0.0, 20.0 ]
fixed_tensor = TRUE

dataset = [ ../input/pcs/ytn_tb_pcs.dat, Tb, 1.0, CONST, MEAN, SVD, 7.585, 24.964, 8.048,  23.966,
15.288,  79.312,  68.199, 151.578 ]

dataset = [ ../input/pcs/ytn_tm_pcs.dat, Tm, 1.0, CONST, MEAN, SVD, 7.585, 24.964, 8.048,  19.071,
6.842, 162.631, 113.645, 170.187 ]

dataset = [ ../input/pcs/ytn_dy_pcs.dat, Dy, 1.0, CONST, MEAN, SVD, 7.585, 24.964, 8.048, -21.164, -
11.060, 131.562, 136.607, 140.271 ]

END
```

##### 3.3) Ligand rigid body grid search with PCSs

- In this protocol, global ligand docking is attempted, which is a notoriously difficult task. In order to leverage the full potential of the PCS in restricting the conformational search space and narrow down ligand docking to a few favorable binding regions with low PCS score, a grid search of the ligand's rigid body degrees of freedom in a shell around the protein is performed. Moreover, different ligand conformations (20 in this protocol capture) are tested. The grid nodes corresponding to centers-of-mass of the ligand with the lowest PCS score are written to a PDB file which then serves as input for RosettaLigand in the next step of the protocol. Optionally, the rigid body grid search can be followed by a short high-resolution docking step aiming to create a physically realistic all-atom model of the protein-ligand encounter complex.

- Change into the sub-directory ***3_rbsearch*** and execute the *ligand_transform_with_pcs* command line application.

```
~/Rosetta/main/source/bin/ligand_transform_with_pcs.linuxgccrelease -help
```

This will print the relevant options for this application together with their descriptions and default values. Now run the application with the provided options file (*rbsearch.options*). Make sure that the path to the PDB, PCS and residue parameter file of pYTN, which are all three located in the ***input*** directory, is correct.

```
~/Rosetta/main/source/bin/ligand_transform_with_pcs.linuxgccrelease @
rbsearch.options
```

By default, 20 independent trajectories with translational and rotational magnitudes of 4 Å and 20°, respectively, are performed and the GRB2 and pYTN coordinates are written out as PDB files. Furthermore, the centers-of-mass (COM) of pYTN docking poses at the end of each calculation are stored in the PDB file *YTN_centroids_pcs_dock.pdb* as HETATM records. Notice, that the pYTN COMs are located close together indicating that the PCS grid search has found one favorable docking position with low PCS score.

##### 3.4) Run Rosetta protein-ligand docking with PCSs

- Change into the sub-directory ***4_docking*** from which you will run RosettaLigand. Use the default settings given in the options file (*ligdock.options*) and Rosetta XML script (*ligdock.pcs.xml*). For a detailed description of the various components of the RosettaLigand protocol as implemented in Rosetta scripts we refer the reader to references [6] and [7]. Make sure that the path to the PDB file containing the pYTN centers-of-mass from the previous step is correctly parsed to the StartFrom Mover. Furthermore, the path to the PCS input file of pYTN must be provided as argument to the PCSMultiGrid in the <SCORINGGRIDS> section of the Rosetta script. The weight of the PCS ligand grid score (e.g. 20 in this protocol capture) was determined similar to protocols 1) and 2) by optimizing it against the total ligand grid score (“total_score_X”). The weight of the PCS score in the ligand full-atom scoring function, as defined at the beginning of the Rosetta XML script in the <SCOREFXNS> section, is set to 1/10 of the PCS ligand grid score:

```
<Reweight scoretype=”nmr_pcs” weight=”2.00”/>
```

- Start RosettaLigand by typing into the terminal:

```
~/Rosetta/main/source/bin/rosetta_scripts.linuxgccrelease -parser:protocol
ligdock.pcs.xml @ ligdock.options
```

- A Rosetta silent file (*GRB2_YTN_dock_pcs.out*) and score file (*GRB2_YTN_dock_pcs.sc*) containing 10 models of GRB2 with pYTN will be created. Increase the value of the “-out:nstruct” flag if you wish to create more models. Typically, 10,000 – 100,000 models are generated for protein-ligand docking and calculations are run on multiple CPUs on a cluster. If you wish to extract docking models as PDB files from the silent file you can again use Rosetta’s *extract_pdbs* application. This time, you also need to provide the pYTN params file with the “-in:file:extra_res_fa” flag.

- Finally, the ligand interface score, the PCS score and the ligand RMSD without superimposition relative to the X-ray structure of GRB2-pYTN can be extracted from the score file by using the python script *extract_scores.py.* Values can then be plotted to create Score-vs-RMSD plots (see **Figure 3**).

```
../../scripts/extract_scores.py GRB2_YTN_dock_pcs.sc --columns interface_delta_X
nmr_pcs ligand_rms_no_super_X description
```

Alternatively, the ligand RMSD can be calculated relative to the lowest-scoring docking solution by extracting the best model from the silent file and parsing it as native PDB to the InterfaceScoreCalculator Mover in the Rosetta XML script.

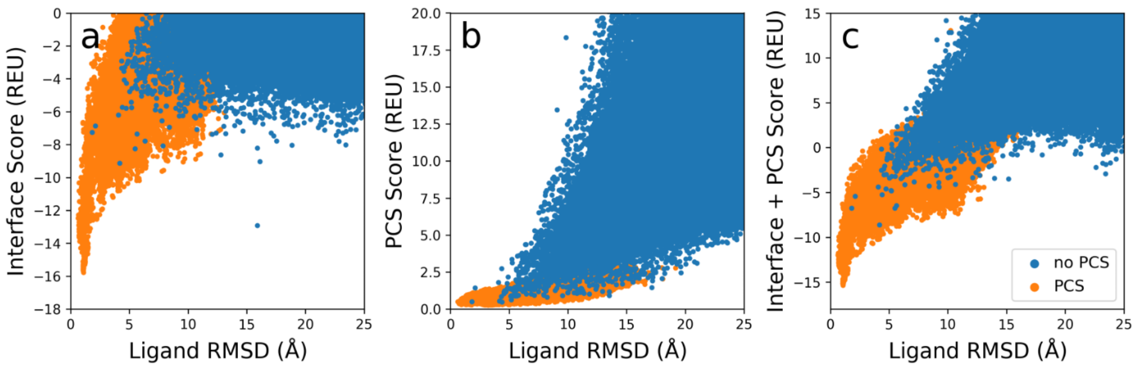


**Figure 3: Docking of a pYTN tripeptide ligand to the SH2 domain of GRB2 with and without PCS data.** The Rosetta interface score (**a**), PCS score (**b**) and combined interface and PCS score (**c**) is plotted versus the ligand RMSD relative to the experimental structure calculated without superimposition.

**4) Protein-protein docking**

This protocol introduces how to use RosettaNMR together with PCSs for protein-protein docking. A heterodimeric protein complex consisting of the ε and θ subunits of *E. coli* DNA polymerase III serves as example system. A structural model of the complex (PDB 2XY8) was previously determined from PCS data [8]. The protein exhibits a natural metal-binding site formed by residues D12, E14 and D167 which was employed for lanthanide ion tagging.

Navigate to the folder ***4_protein_docking***. There are four sub-folders located in this directory, one ***input*** folder and three additional folders, one for each of the three steps of this protocol:

- pre-packing of protein docking partners (***1_prepack***)
- protein-protein docking with PCSs (***2_docking***)
- re-scoring of docking models with PCSs (***3_rescore***)

The ***input*** directory contains the following files:

- 2XY8 structure in PDB format; only the first model of the NMR ensemble (*2xy8.pdb*)
  Note that the protein sequence has been renumbered to comply with Rosetta's pose numbering scheme according to which the protein sequence starts at position 1 and all residues are numbered consecutively. The ε subunit is chain A (residues 1-174) and the θ subunit is chain B (residues 175-235). The calcium ion in the PDB file has been removed because of current difficulties in running Rosetta docking in the low-resolution centroid stage with metal ion residues. Removing metal ions and other ligands from the PDB file and renumbering of residues can be done with Rosetta’s

*clean_pdb.py* script which can be found in the Rosetta tools repository under ***~/Rosetta/tools/protein_tools/scripts/clean_pdb.py***). For scoring with PCSs, it is not necessary that the metal ion is part of the structural model, and we can define a coordinating protein residue as grid search center in determining the $\Delta\chi$-tensor position.

- PCS input file (*2xy8.pcs.inp*)
- PCS data files (in Rosetta format); located in directory ***pcs***; one data file for each of the three lanthanides ($Tb^{3+}$, $Dy^{3+}$, $Er^{3+}$)

**4.1) Pre-packing of protein docking partners**

- To prepare the two protein chains for docking with Rosetta, the Docking Prepack application is run, which optimizes side chain conformations by rotamer packing. This is to ensure that the side chains outside of the docking interface have low energy conformations which is essential for model scoring.

- Change into the sub-directory ***1_prepack*** and run Docking Prepack by typing the following command into the terminal:

```
~/Rosetta/main/source/bin/docking_prepack_protocol.linuxgccrelease @
prepack.options
```

  Use the options provided in the text file *prepack.options*.

**4.2) Run Rosetta protein-protein docking with PCSs**

- Change into the directory ***2_docking***. For running Rosetta protein-protein docking with PCSs, the Rosetta Scripts application is used. An input Rosetta XML script (*docking.pcs.xml*) as well as an options file (*docking.options*) have been prepared for you in this directory. Have a look at the options file. Since we are running global docking, the orientation of the two docking partners, as specified with the "-docking:partners" flag, is randomized at the beginning of each trajectory. The flags "-docking:randomize1" and "-docking:randomize2" in the options file randomize the orientation of the first and second docking partner, respectively. If you do not wish to perform global protein-protein docking, do not use those flags. In addition, another initial rigid-body perturbation of the two protein subunits with a translational and rotational magnitude of 3 Å and 8°, respectively, is performed (see the "-docking:dock_pert 3 8" flag in the options file).

- Use the weight of the PCS score as provided in the <SCOREFXNS> section of the Rosetta XML Script. It has been determined in similar manner as described before in protocols 1) and 2) by first creating 10000 decoys without PCSs and afterward re-scoring them with PCSs. The PCS score weight was thereby optimized against the binding energy after separating and re-packing both docking partners (i.e. the ddg score in Rosetta) using the ref2015 weight set.

- Run Rosetta protein-protein docking by typing the following command into the terminal:

```
~/Rosetta/main/source/bin/rosetta_scripts.linuxgccrelease -parser:protocol
docking.pcs.xml @ docking.options
```

  which will run docking on the pre-packed model from step 4.1) and create 3 output models (*2xy8_docking.out*). Alternatively, docking can be initiated from an ensemble of different input structures as specified by a list file ("-in:file:l"), and the number of output models can be increased by setting the "-out:nstruct" flag to a larger value. Typically, between 10,000 to 100,000 models are created for protein-protein docking and calculations are run on multiple CPUs on a cluster.

##### 4.3) Rescoring of docking models with PCSs

- After the docking run is completed, score the output models with PCS data again. This step is necessary because the PCS score which was reported in the score file at the end of the docking protocol was evaluated when calculating the Rosetta binding energy (ddg). Since the ddg-calculation is done by separating the two binding partners the PCS score is incorrect because it must be evaluated on the complete protein complex in the bound state.
  For re-scoring docking models with PCSs, use the Rosetta *score_jd2* application:

```
~/Rosetta/main/source/bin/score_jd2.linuxgccrelease @ rescore.pcs.options -
in:file:silent ../2_docking/2xy8_docking.out -out:file:scorefile
2xy8_docking_pcs_rescored.sc
```

- Finally, the binding energy (ddg), the PCS score and the model's interface RMSD relative to the experimental structure of 2XY8 can be extracted from the score file by using the python script *extract_scores.py,* and values are plotted to create Score-vs.-RMSD plots (see **Figure 4**).

```
../../scripts/extract_scores.py 2xy8_docking_pcs_rescored.sc --columns ddg
nmr_pcs Irmsd description
```

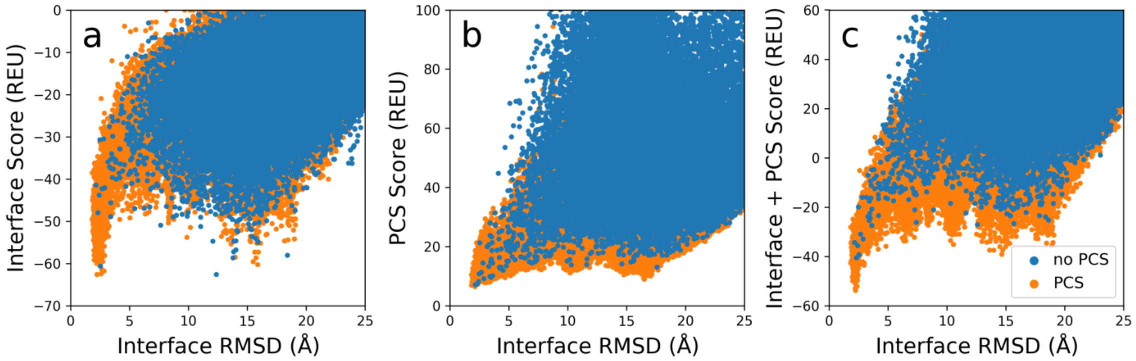


**Figure 4: Docking of the ε and θ subunits of *E. coli* DNA polymerase III with and without PCS data.** The Rosetta interface score (**a**), PCS score (**b**) and combined interface and PCS score (**c**) is plotted versus the interface RMSD relative to the experimental structure.

#### 5) Appendix

##### 5.1) Description of the PCS, RDC and PRE input file format

PCS as well as PRE datasets that were measured with different types of spin-labels or at different spin-label sites, and RDC datasets that were collected with different alignment media (or metal ions) must be placed into separate sections in the input file. Each section must start with the keyword “**MULTISET**” and end with the keyword “**END**”. PCS datasets obtained with different metal ions but at the same spin-label site should be grouped within the same MULTISET section because during the calculation their metal ion coordinates are optimized together. In case of RDCs, datasets for different atom types (e.g. N-H, CA-HA) that were measured with the same type of alignment medium should be grouped in the same MULTISET section because they share the same alignment tensor. And for PREs, the MULTISET section includes PRE datasets that were collected with the same metal ion or radical at the same spin-label site. It is also recommended to place $R_1$ and $R_2$ relaxation data into separate MULTISETs or apply pre-scaling because their values usually differ by several orders of magnitude. Furthermore, relaxation data measured under different experimental conditions (temperature, protein mass, viscosity) should be placed into separate MULTISET sections too, since the spin-label and protein correlation times will be different.

Each MULTISET section contains a collection of keyword-value pairs. Each pair must be separated by an equal sign (=) and placed on a new line. The value can be a numerical value, character, literal string or a list of any of the latter types. Multiple values are grouped in a vector, enclosed by brackets ([ ]) and separated by commas. A list of the possible MULTISET keywords for each of the three types of input files is provided in **Table 1**.

One additional remark on the types of weights that can be applied to the input data in scoring. In addition to the global weights for PCSs, RDCs or PREs which are set through the Rosetta scoring function, weighting of the input NMR data is possible on three additional levels:

- MULTISET weights (this option is available for PCSs, RDCs and PREs and weights are assigned through a command line flag, e.g. "-nmr:pcs:multiset")
- SINGLESET weights (Single PCS or PRE datasets/experiments that were obtained with different meal ions but belong to the same MULTISET can be assigned different weights through a positional argument in the dataset vector in the input file; see **Table 1** below)
- Each single data point is assigned a weight that depends on its error and magnitude (possible modes are CONST, SIGMA or OBSIG; see explanations in the footnote to **Table 1**)

**Table 1: Keywords of the PCS, RDC and PRE input file.**

| PCS input file | | | |
|---|---|---|---|
| **Keyword** | **Explanation** | **Required?** | **Example** |
| spinlabel_position | Residue number of the spin-label site. This residue becomes the anchor point for the grid search of the metal ion coordinates. Alternatively, if a value for the keyword *spinlabel_type* is provided, the grid search will be replaced by a dummy spin-label residue that is modeled at this protein position in order to determine the metal ion coordinates. | Yes | spinlabel_position = 58 |
| chain_id | The ID of the protein chain which includes the spin-label site. | Yes | chain_id = A |
| gridsearch | Vector of parameters that define the metal ion grid search. The elements of this vector are in the following order: names of atoms 1 and 2 which are used to define the grid search center, the distance between atom 1 and the grid search center (in Å), the stepsize of the grid search (in Å), and the minimal and maximal radius of the grid search (in Å). The center of the grid search lies on a line running through atoms 1 and 2 and is located at a defined distance from atom 1. The metal ion position with the lowest PCS score is searched within a range between the minimal and maximal radius around the grid search center. See also **Figure 5** for an explanation of the grid search parameters. | Yes, but can be replaced by the *spinlabel_type* keyword | gridsearch = [ CA, CB, 10.0, 4.0, 0.0, 20.0 ] |
| spinlabel_type | Three-letter code of the spin-label residue type as stored in the Rosetta database. For example, R1A is the code of the MTSL spin-label residue. (see | Yes, but can be replaced by *gridsearch* keyword | spinlabel_type = R1A |

| | | | |
|---|---|---|---|
| | ~/Rosetta/main/database/scoring/nmr/spinlabel/spinlabel_properties.txt for a list of currently available spin-label residue types in Rosetta) | | |
| dataset | Vector of input parameters for one single PCS dataset collected from one metal ion. The vector must contain exactly 14 elements in the following order: Name and location of PCS data file, metal ion label, weighting factor for this PCS dataset, type of weighting single PCS values (see explanations below*), averaging type for a PCS value that is assigned to a group of atoms (MEAN or SUM), type of $\Delta\chi$-tensor fit (SVD = singular value decomposition, NLS = non-linear least squares fitting), and the eight $\Delta\chi$-tensor parameters ($x_M$, $y_M$, $z_M$, $\chi_{ax}$, $\chi_{rh}$, $\alpha$, $\beta$, $\gamma$). In many cases e.g. de novo structure prediction, the tensor parameters can be set to random values as they will be determined automatically during the calculation. However, other protocols, e.g. ligand docking with PCSs, require that the tensor parameters are determined prior to the Rosetta calculation and entered in the dataset vector. | Yes | dataset = [ 2k61_dapk_exp_pcs_dy.txt, Dy, 1.0, CONST, MEAN, SVD, 10.0, 10.0, 10.0, 40.0, 4.0, 10.0, 10.0, 10.0 ] |
| fixed_tensor | Do not fit the $\Delta\chi$-tensor but calculate the PCS score from the tensor values entered into the dataset list. This is needed for protein-ligand docking (Default: false). | Optional | fixed_tensor = true |

**PRE input file**

| Keyword | Explanation | Required? | Example |
|---|---|---|---|
| spinlabel_position | See explanation of this keyword in PCS input file above. | Yes | spinlabel_position = 58 |
| chain_id | See explanation of this keyword in PCS input file above. | Yes | chain_id = A |
| spinlabel_type | See explanation of this keyword in PCS input file above. | Yes, but can be replaced by *gridsearch* keyword | spinlabel_type = R1A |
| gridsearch | See explanation of this keyword in PCS input file above. See also **Figure 5** for an explanation of the grid search parameter. | Yes, but can be replaced by the *spinlabel_type* keyword | gridsearch = [ CA, CB, 10.0, 4.0, 0.0, 20.0 ] |
| ion_type | Name of radical atom or paramagnetic ion as stored in the Rosetta database (see ~/Rosetta/main/database/chemical/element_sets/default/para_ion_properties.txt for reference) | Yes | ion_type = Cu2+ |
| protein_mass | Molecular mass of the protein (in kDa). | Yes | protein_mass = 6.2 |
| temperature | Temperature at which the PRE datasets were recorded (in Kelvin). | Yes | temperature = 278.0 |
| dataset | Vector of input parameters for one | Yes | dataset = |

| | | | |
|---|---|---|---|
| | single PRE dataset. Must contain exactly five elements in the following order: Name and location of PRE data file, weighting factor for this PRE dataset, type of weighting single PRE values (see explanations below*), type of relaxation data (R1 or R2), magnetic field strength in MHz. | | [ 8c_edta_cu2_nr1_alt.txt, 1.0, CONST, R1, 500 ] |
| averaging | Averaging type for a PRE value that is assigned to a group of atoms. Possible values are MEAN and SUM (Default: MEAN). | Optional | averaging = MEAN |
| tauc_min | Lower bound for tauc$ (in ns). | Optional | tauc_min = 1.0 |
| tauc_max | Upper bound for tauc$(in ns). | Optional | tauc_max = 15.0 |

**RDC input file**

| Keyword | Explanation | Required? | Example |
|---|---|---|---|
| alignment_medium | Name of the alignment medium. This value is not actually used in the calculation but has a purely descriptive purpose. | Yes | alignment_medium = phage |
| computation_type | Type of the alignment tensor fit. Possible choices are: SVD (singular value decomposition), NLS (non-linear least squares fitting), NLSDA (NLS with fixed value for alignment magnitude Da), NLSR (NLS with fixed value for rhombicity R), NLSDAR (NLS with fixed value for Da and R). Notice that for computation types NLSDA, NLSR and NLSDAR the alignment magnitude Da and/or rhombicity R must be provided in the input file too. | Yes | computation_type = SVD |
| dataset | Vector of input parameters for one single RDC dataset collected for one atom type. The vector must contain exactly three elements in the following order: Name and location of RDC data file, weighting factor for this RDC dataset, type of weighting single RDC values (see explanations below*). | Yes | dataset = [ 2k61_dapk_nh.txt, 1.0, SIGMA ] |
| alignment_tensor | Vector of the five alignment tensor values in the following order: Da, R, α, β, γ. If one of the computation types NLSDA, NLSR or NLSDAR is chosen the value of Da and/or R will be read from this vector. All other values will be re-determined during the calculation and can be set to random start values. | Yes | alignment_tensor = [ -14.217, 0.530, 10.0, 10.0, 10.0 ] |
| averaging | Type of averaging an RDC value if assignment spans a group of atom pairs. Possible values are MEAN and SUM (Default: MEAN). | Optional | averaging = MEAN |
| fixed_tensor | Do not fit the alignment tensor but calculate the RDC score from the values provided in the *alignment_tensor* list (Default: false). | Optional | fixed_tensor = true |

*Three choices for weighting the contribution of single data points to the total score exist:
CONST: all data points have equal weight, i.e. $w = 1.0$
SIGMA: data points have a weight proportional to the inverse of their error, i.e. $w = \frac{1}{\sigma^2}$
OBSIG: data points have a weight proportional to the inverse of their error and their magnitude relative to the maximal observed value, i.e. $w = \frac{1}{\sigma^2} \times \frac{\delta^{obs}}{\delta^{obs,max}}$
$See reference [9] for definition of these correlation times.

**5.2) Description of additional PCS, RDC and PRE options**

**Table 2: PCS, RDC and PRE options.**

| **PCS options** | |
|---|---|
| -nmr:pcs:input_file | Name of PCS input file (the only required flag; see format description above). |
| -nmr:pcs:optimize_tensor | Optimize the metal ion position and the other values of the Δχ-tensor after initial grid search (Default: false). |
| -nmr:pcs:nls_repeats | Number of repeats in non-linear least squares fitting of the Δχ-tensor (Default: 5). |
| -nmr:pcs:multiset_weights | Vector of weights by which the scores from different PCS multi-sets are multiplied. One multi-set includes all PCS datasets that were collected at the same spin-label site but with different metal ions. Defaults to a vector of 1.0 if not explicitly set. |
| -nmr:pcs:normalize_data | Normalize PCS values of every dataset by their standard deviation (Default: false). |
| -nmr:pcs:use_symmetry_calc | Calculate the PCS score by considering the contributions from all symmetric subunits (Default: false). PCS values should be assigned only to the atoms of the asymmetric subunit in the data input file. Note, this option was developed for cases of Cn- and Dn-symmetry but not tested for systems with other types of symmetry. |
| -nmr:pcs:show_info | Show Δχ-tensor and a table of experimental vs. calculated PCS values at every scoring step (Default: false). Note, to print this information to the screen or the log file it is also required that the tracer output level ("-out:level") is set to 500 (debug mode). Be careful, this will make the log file very large! |
| **RDC options** | |
| -nmr:rdc:input_file | Name of RDC input file (the only required flag; see format description above). |
| -nmr:rdc:nls_repeats | Number of repeats in non-linear least squares fitting of alignment tensor (Default: 5). |
| -nmr:rdc:multiset_weights | Vector of weights by which the scores from different RDC multi-sets are multiplied. One multi-set includes all RDC datasets that were collected with the same alignment medium or lanthanide ion and multiple different atom types (e.g. N-H, CA-HA). Defaults to a vector of 1.0 if not explicitly set. |
| -nmr:rdc:normalization_type | Apply scaling to the input RDC values with respect to those of a chosen atom type. Possible options are NH, CH or none. By default, RDCs are not expected to be normalized and will be scaled relative to the NH dipolar coupling constant (option "NH"). Alternatively, the input RDCs can be scaled relative to the CA-HA dipolar coupling constant (option "CH") or remain unmodified (option "none"). |
| -nmr:rdc:correct_sign | Use the correct sign of the $^{15}N$ gyromagnetic ratio and thus of the dipolar coupling constants i.e. positive for NC and NH and negative for CH (Default: false). Use this option if input couplings have different signs. By default, the $^{15}N$ gyromagnetic ratio is treated as positive and the dipolar coupling constants have the same sign. |
| -nmr:rdc:use_symmetry_calc | Calculate the RDC score by considering the contributions from all symmetric subunits (Default: false). RDC values should be assigned only to the atoms of the asymmetric subunit in the data input file. Note, this option was developed for cases with Cn- and Dn-symmetry but not tested for systems with other types of symmetry. |
| -nmr:rdc:show_info | Show alignment tensor and a table of experimental vs. calculated RDC values at every scoring step (Default: false). Note, to print this information to the screen or the log file it is also required that the tracer output level ("-out:level") is set to 500 (debug mode). Be careful, this will make the log file very large! |

| PRE options | |
|---|---|
| -nmr:pre:input_file | Name of PRE input file (the only required flag; see format description above). |
| -nmr:pre:nls_repeats | Number of repeats in non-linear least squares fitting of spin-label correlation time (Default: 3). |
| -nmr:pre:multiset_weights | Vector of weights by which the scores from different PRE multi-sets are multiplied. One multi-set includes all PRE datasets that were collected at the same spin-label site with the same type of spin-label or metal ion. Defaults to a vector of 1.0 if not explicitly set. |
| -nmr:pre:normalize_data | Normalize PRE values of every dataset by their standard deviation (Default: false). |
| -nmr:pre:use_symmetry_calc | Calculate the PRE score by considering the contributions from all symmetric subunits (Default: false). PRE values should be assigned only to the atoms of the asymmetric subunit in the data input file. Note, this option was developed for cases with Cn- and Dn-symmetry but not tested for systems with other types of symmetry. |
| -nmr:pre:show_info | Show a table of experimental vs. calculated PRE values at every scoring step (Default: false). Note, to print this information to the screen or the log file it is also required that the tracer output level ("-out:level") is set to 500 (debug mode). Be careful, this will make the log file very large! |
| **Spin-label options** | |
| -nmr:spinlabel:max_ensemble_size | Maximum number of spin-label rotamers to represent the ensemble (Default: 20). If the ensemble size is higher than this cutoff, rotamers are binned by their side chain RMSD. This is to speed up the calculation. |
| -nmr:spinlabel: highres_conformer_filter_type | Type of detecting clashes of spin-label conformers with neighboring protein residues in Rosetta's full-atom mode. Possible values are DISTANCE and BUMP_ENERGY. In the first case, conformers are treated as having a clash if any of their side chain heavy atoms is within the neighbor radius of any of the neighboring amino acid residues. In the second method, clashes are detected by calculating the bump energy of a spin-label conformer with its protein environment. The default value is BUMP_ENERGY, whereas in Rosetta's centroid stage the DISTANCE method is the only one that is available. (Default: BUMP_ENERGY). |
| -nmr:spinlabel: elaborate_rotamer_clash_check | Perform an elaborate clash check of every heavy atom of all spin-label rotamers with the heavy atoms in surrounding residues (Default: false). If all rotamers produce at least one clash, return that rotamer with the lowest number of atom clashes. If false, consider only heavy atoms in non-clashing rotamers and return a random spin-label rotamer in case there are no remaining non-clashing rotamers. |
| -nmr:spinlabel:boltzmann_kt | Scaling factor for Boltzmann weighting of the spin-label conformer probabilities (Default: 2.0). |

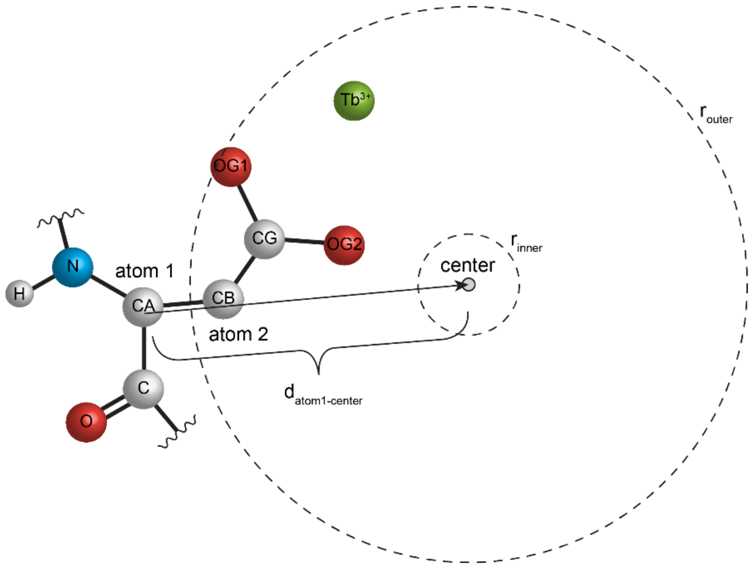


**Figure 5: Explanation of the metal grid search parameters.** The grid search center is defined relative to a protein residue that is in close proximity to the metal ion (e.g. $Tb^{3+}$). The center lies on a vector running through atoms 1 and 2, and is located at a given distance $d_{atom1\text{-}center}$ from atom 1. Usually, the CA and CB atom are chosen as atoms 1 and 2 because they are present Rosetta's centroid and all-atom residue types. The grid search is performed within the volume confined by two spheres which have radii $r_{inner}$ and $r_{outer}$.

#### 6) References


1. Peters, F., et al., *Cys-Ph-TAHA: a lanthanide binding tag for RDC and PCS enhanced protein NMR.* J Biomol NMR, 2011. **51**(3): p. 329-37.
2. Olivieri, C., et al., *Simultaneous detection of intra- and inter-molecular paramagnetic relaxation enhancements in protein complexes.* J Biomol NMR, 2018. **70**(3): p. 133-140.
3. Gront, D., et al., *Generalized fragment picking in Rosetta: design, protocols and applications.* PLoS One, 2011. **6**(8): p. e23294.
4. Parsons, L.M., A. Grishaev, and A. Bax, *The periplasmic domain of TolR from Haemophilus influenzae forms a dimer with a large hydrophobic groove: NMR solution structure and comparison to SAXS data.* Biochemistry, 2008. **47**(10): p. 3131-42.
5. Dimaio, F., et al., *Modeling symmetric macromolecular structures in rosetta3.* PLoS One, 2011. **6**(6): p. e20450.
6. Lemmon, G. and J. Meiler, *Rosetta Ligand docking with flexible XML protocols.* Methods Mol Biol, 2012. **819**: p. 143-55.
7. DeLuca, S., K. Khar, and J. Meiler, *Fully Flexible Docking of Medium Sized Ligand Libraries with RosettaLigand.* PLoS One, 2015. **10**(7): p. e0132508.
8. Schmitz, C. and A.M. Bonvin, *Protein-protein HADDocking using exclusively pseudocontact shifts.* J Biomol NMR, 2011. **50**(3): p. 263-6.
9. Iwahara, J., C.D. Schwieters, and G.M. Clore, *Ensemble approach for NMR structure refinement against $^{1}H$ paramagnetic relaxation enhancement data arising from a flexible paramagnetic group attached to a macromolecule.* J Am Chem Soc, 2004. **126**(18): p. 5879-96.
